# Archetype analysis of multiplexed antibody imaging reveals interpretable morphological programs in optical pooled screens

**DOI:** 10.64898/2026.10.07.757470

**Authors:** Paula Pinto Coelho, Ana Carolina Leote, Heming Yao, Ping Wu, Taka Kudo, Hanchen Wang, Vladimir Ermakov, Alisha Saboowala, Taro Makino, Russell Littman, Romain Lopez, Joseph Kim, Tonia Hafner, Orit Rozenblatt-Rosen, David Richmond, Levi Garraway, Avtar Singh, John Marioni, Aviv Regev

## Abstract

High-content imaging-based screens with small molecule or genetic perturbations rely on rich optical phenotypes, yet how distinct subcellular markers contribute complementary biological information remains unclear. Here, we introduce CellMAPP, a multiplexed organelle imaging platform that uses optical pooled screens with iterative antibody staining to map cellular architecture under perturbation. Using 17 markers across 524 genetic perturbations in MCF-7 cells, we systematically examined how individual, and combinations of, markers capture diverse morphological responses. We find that markers provide complementary views of perturbation effects, with increased multiplexing improving signal and the breadth of hit recovery, although some phenotypes are better recovered from a subset of features rather than a full profile. Because non-molecular image-based profiles can be challenging to link with interpretable biology, we developed CHART (available at https://github.com/Genentech/CHART), an archetype analysis framework that identifies distinct reference morphological states for each marker and uses multimodal large language models (MLLMs) to automatically annotate them, connecting cellular images to recognizable organelle phenotypes. Integrating archetypes across multiple markers revealed modular morphological programs that describe coordinated organelle reorganization in response to perturbation. This framework identified regulators of defined organelle states such as micronuclei, nominated novel gene functions and revealed complex patterns of multi-organelle reposition. Together, optical profiling with MAPP and CHART provides an interpretable, scalable framework for image-based screens that sheds light on rules governing cellular structural reprogramming.

## INTRODUCTION

Cellular morphology is an information-rich readout of functional states, making image-based profiling a powerful platform for high-throughput biological screens of genetic perturbations and small molecules. Recently, Optical Pooled Screening (OPS) has enabled large-scale genetic screens with morphological readouts by combining imaging with targeted *in situ* sequencing (ISS) of perturbation barcodes, from bacteria^1,2^ to mammals^3^. OPS has been used for investigation of specific biological phenotypes (e.g., transcription factor translocation, viral infection, mitosis, synaptogenesis, cell growth and replication dynamics)^4–8^ as well as hypothesis-free morphological profiling of thousands of genetic perturbations using small (≤5) marker panels^9^. The rapid adoption of these approaches, together with steadily growing datasets, is positioning image-based profiling as a scalable and increasingly central modality for functional genomics.

While the number and nature of markers and features we can measure continue to grow, we do not generally understand their contributions as distinct phenotypes. Open questions include which compartments capture non-redundant biological information, in terms of the relation between perturbations; whether integrating across stains for different compartments helps resolve distinct cellular states; and how to translate complex imaging profiles with visual features into interpretable biological insights^10,11^. Current approaches rely on high-dimensional representations derived from manually-engineered features^12^ or deep learning embeddings^13–17^. Although simple measurements such as molecular intensity are readily interpretable, complex features (e.g., Haralick textures, Hu moments) and learned embeddings capture variation in ways that are difficult to map onto concrete biological phenotypes. As a result, these approaches can robustly quantify phenotypic effects without readily explaining them in biologically interpretable terms. Tackling these challenges requires both experimental advances to generate imaging-based perturbation screens with sufficient marker complexity and computational frameworks to map the performance of different marker combinations and link single-cell embeddings with biologically interpretable phenotypes.

Here, we developed CellMAPP (Cell Multiplexed Antibody Phenotypic Profiling), a platform combining 17-plex iterative immunofluorescence of subcellular structures present in most cell types with OPS, and use it to study MCF-7 breast cancer cells. Systematic evaluation of how markers recover perturbation effects, both individually and across 16,530 channel combinations revealed complementary biological information across compartments and improved detection sensitivity with increasing panel complexity. We then developed CHART (Cellular High-content imaging Archetype Response Toolkit), an AI-driven computational framework where imaging features are transformed into interpretable morphological archetypes for each marker, and then a scaled vision-language model workflow converts archetype images into textual descriptions via structured prompting, facilitating interpretation of perturbation effects. Finally, sparse PCA of the recovered archetypes defined latent morphological programs that capture coordinated phenotypic changes across multiple markers.

The 188 recovered archetypes enabled multiple biological discoveries including 1) the identification of regulators of specific organelle phenotypes, such as micronucleus formation, 2) recovery of unexpected gene functions such as a role for the splicing factor ACIN1 in endosomal trafficking, and 3) resolution of modularity in morphological response, where functionally distinct perturbation classes drive tailored cellular responses by impacting different combinations of programs of morphological changes. Overall, our approach helped decipher how distinct cellular pathways actively maintain and reconfigure subcellular spatial organization.

## RESULTS

### High-content antibody multiplexing for morphological profiling of genetic perturbations

Phenotyping in image-based screens has traditionally relied on small marker panels, most commonly Cell Painting^10,18^, which uses 5 dyes staining general cellular structures, and has produced informative maps of genetic and chemical perturbation relationships. A recent expansion to seven dyes targeting nine compartments enabled recovery of novel mechanisms of action by explicitly attributing perturbation effects to added compartments^19^, suggesting that increased plexity may improve not only detection sensitivity but also interpretability by linking effects to defined markers.

To study the relationship between panel complexity and biological understanding, we developed CellMAPP and applied it in an OPS of MCF-7, an ERα-positive breast cancer cell line. We designed a 17-marker antibody-based panel (**Fig. 1a**) targeting structures present across most cell types that can integrate signals from many pathways: 13 organelle markers (*e.g.*, TOM20:mitochondria, CALR:endoplasmic reticulum, CAT:peroxisome), two markers for general processes (Ki67:proliferation, phospho-RPS6:translation), as well as two system-specific markers (ERα:estrogen signaling CTNNB1:adherens junctions) (**Fig. 1a)**. We reasoned that unlike Cell Painting dyes, antibody-based staining provides greater molecular specificity, and thus should enable tractable interrogation of how individual compartments respond to perturbation. The screen library consisted of guides targeting 524 genes (4 guides/gene), curated based on known morphological effects^6,13^ or functional connections on individual imaged compartments (**Supplementary Table 1**). We transduced a high-efficiency knockout clonal MCF-7 line, seeded cells onto glass-bottom plates and imaged them across four rounds of staining with iterative bleaching^20^, before barcode detection using PerturbView^21^ (**Supplementary Fig. 1**, **Methods**). We mapped *in situ* sequencing (ISS) reads and extracted 11,804 morphological features using SCALLOPS^22^, then applied a custom analysis pipeline for tabular imaging data (**Fig. 1b, Supplementary Fig. 2a**) with feature and object filtering, followed by quantile normalization to eliminate well-level batch effects without the low standard deviation (SD) bias introduced by z-scoring (**Supplementary Fig. 2b, Supplementary Table 2**). After aberrant guide and outlier cell removal, this yielded 874,882 high-quality cell profiles.

**Figure 1.**
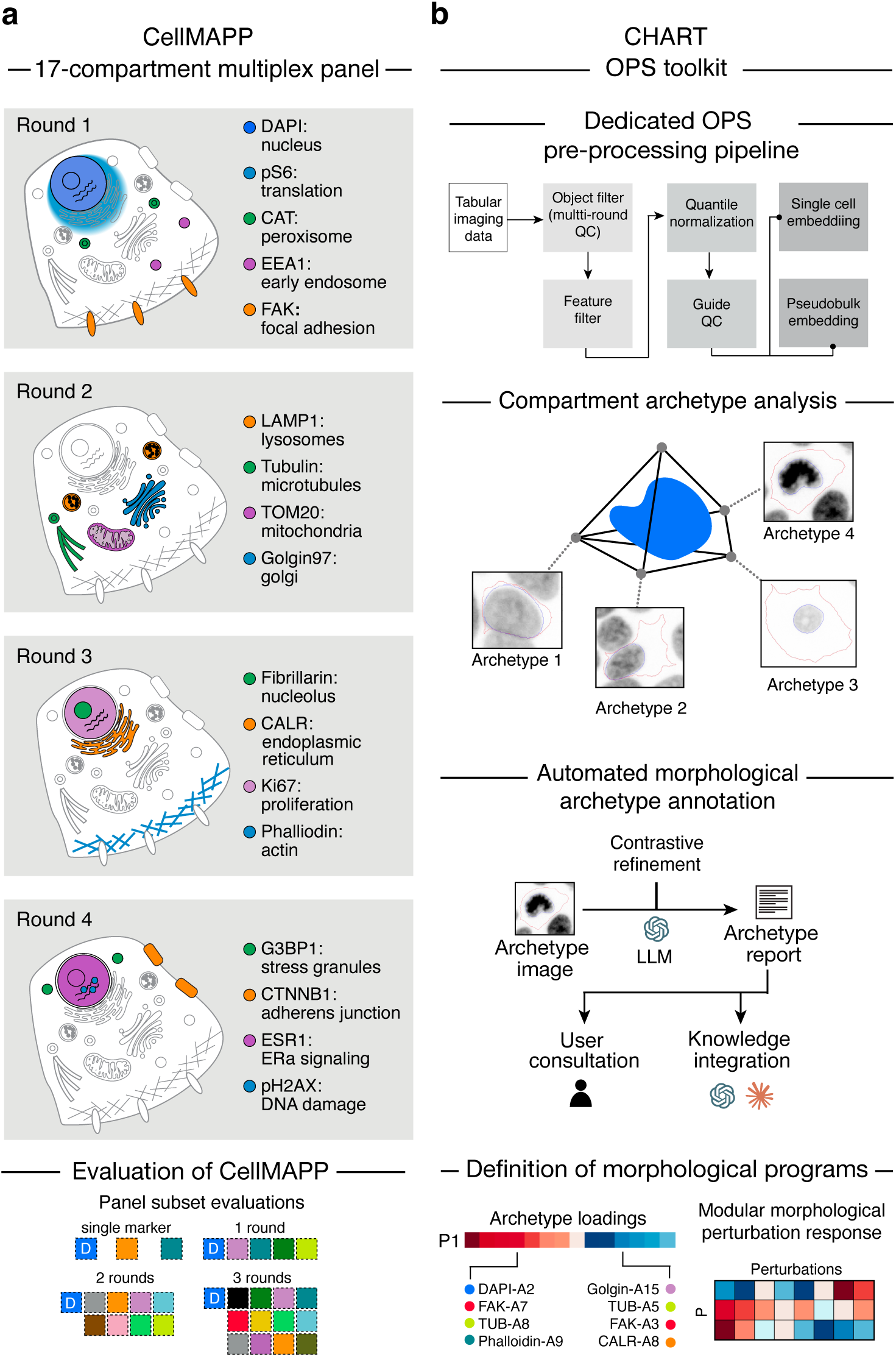
High-content antibody multiplexing for morphological profiling of genetic perturbations. **a.** CellMAPP experimental overview. Schematic of each of the four rounds of multiplexed organelle profiling workflow (top) and of panel subsetting used to evaluate CellMAPP-markers in OPS (bottom). **b.** CHART computational approach. Following pre-processing (top) archetype analysis with MLLM-based annotation (middle) identifies the key morphological phenotypes uncovered by each marker under the perturbations. Archetypes are then grouped into co-regulated morphological programs regulated by co-functional perturbation modules (bottom), which reveal coordinated cell biological effects.

### Multiplexed imaging integrates complementary signals to robustly capture biological relationships

Clustering of perturbation-level profiles (aggregated from single-cell profiles, **Methods**) derived from multiplexed organelle staining recovered coherent sets of genes involved in related biological processes or multi-molecular complexes, including translation, vesicular transport, cytoskeletal organization the ribosome and the proteasome (**Fig. 2a**).

**Figure 2.**
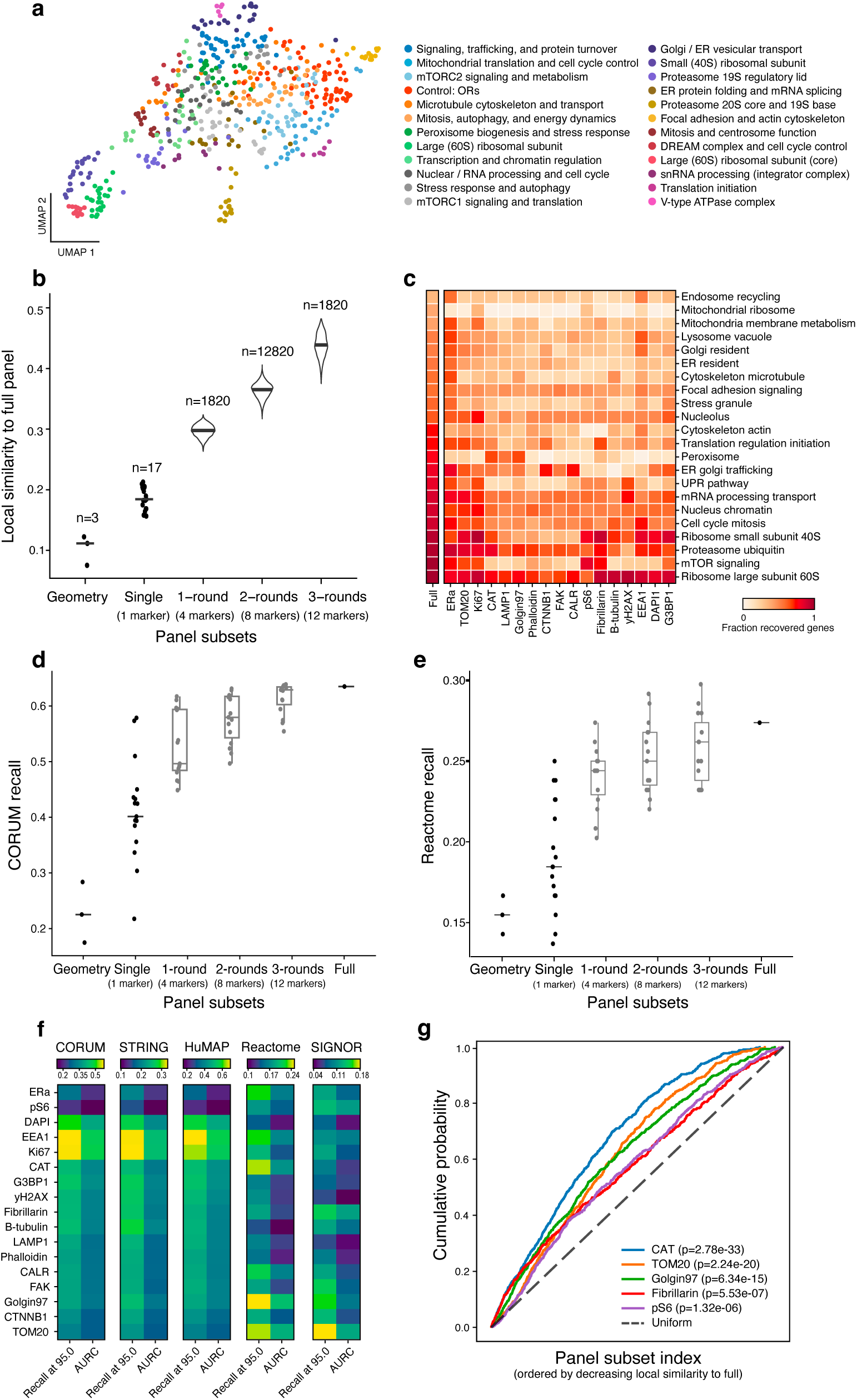
Multiplexed imaging integrates complementary signals to capture biological relationships. **a.** Multiplex imaging groups co-functional perturbations. UMAP embedding of aggregate perturbation imaging profiles (dots, **Methods**) based on 17 organelle markers, colored by clustering and annotated by coherent biological functions (color legend). **b.** Distribution of local similarity to the full panel embedding (y axis, partial AURC@30 of full embedding neighbor recall) for embeddings generated on each class of panel subset (x axis). **c.** Fraction of recovered perturbation effects (color bar) for each functional perturbation group (rows) based on each marker (columns). **d,e** Performance of sub-sampled panel embeddings in recovering known biological relationships in different databases. Recall (fraction of edges from the database reference graph recovered within the 5% strongest positive and negative similarities in the corresponding embedding (recall at 95.0, two-sided)) (y-axis) for 15 random panels selected for the 1, 2 or 3 round categories (x-axis). Random recovery corresponds to 0.1. **e,** Recall from CORUM. **f,** Recall from Reactome **f.** Ability of single markers to recover known biological relationships. Recall (defined as in e) and AURC (average recall across thresholds from 95% to 99% (two-sided)) (columns) for each database (panels, labeled on top) based on embeddings from each individual marker (rows). **g.** Marker enrichment in best performing 1 round panels. Empirical cumulative distribution functions for markers significantly enriched (1-sample KS: p < 0.01) in top performing panels (high local similarity to full embedding). Dashed line: uniform distribution.

To understand how these relationships depend on the composition of the phenotypic panel, we generated perturbation-level PCA embeddings from profiles consisting of each of 16,530 panel subsets spanning 17 single-marker, 3 geometry-feature combinations, and each possible marker-combination configuration across one (r1), two (r2) or three (r3) staining rounds, while retaining DAPI, which is often used to register cells across imaging rounds. We quantified each embedding’s local similarity to the embedding generated with the full panel as the partial area under the neighbor-recall curve (pAUC) with up to 30 neighbors for each gene, using neighbors in the full multiplexed embedding as ground truth (**Methods**, **Fig. 2b**). Local similarity increased monotonously with panel complexity, indicating that additional markers progressively restore the perturbation neighborhoods recovered by the full panel.

Individual markers showed variable sensitivity for detecting different types of perturbations, quantified as the fraction of perturbations in each annotated class that produced significant morphological differences from control (**Fig. 2c**, **Methods**). Perturbations targeting essential complexes, such as the 60S ribosome or proteasome, were detected as impactful by most markers, while other classes, such as mTOR signaling, the peroxisome, or the unfolded protein response (UPR), showed stronger marker specificity. In many cases, markers were most sensitive to perturbations affecting their cognate structures and processes - for example, EEA1 to perturbations in endosome recycling genes and β-tubulin to perturbations in microtubule genes. Most markers, however, also captured several indirect effects. For example, ERα staining was impacted by perturbations in endosome recycling and Ki67 was affected by mitochondrial ribosome perturbations.

We next benchmarked how recovery of known biological relationships scaled with plexity. Since we could not evaluate all 16,530 panel subsets due to computational limitations, we selected 15 random panel subsamples for each level of plexity, to a total of 45. The full panel outperformed the average of any other plexity across five curated gene-gene association databases spanning pathway and signalling relationships (Reactome^23^, SIGNOR^24^) and protein complexes and associations (STRING^25^, CORUM^26^, HuMAP^27^) (**Fig 2d,e, Extended Data Fig. 1a**). Complex/association databases (mean recall based on number of rounds: r1: 0.481, r2: 0.524, r3: 0.553, r4: 0.567) were overall recovered better than pathway/signaling ones (mean recall r1: 0.197, r2: 0.202, r3: 0.212, r4: 0.233). Additionally, we observed a saturation in performance, such that less improvement was achieved with each additional imaging round, especially for complex based databases. However, the best performing panels in pathway-based settings often performed poorly in complex-based settings, and vice versa (**Extended Data Fig. 1b-d)**, with clearer differences observed in 1-round panels. This indicates that higher plexity provides an advantage by enabling simultaneous recovery of different types of effects.

Biological recall of associations from these knowledge-bases was driven by distinct markers, with single-marker performance varying substantially across databases. Golgin97 and TOM20 performed best on pathway/signaling relationships, consistent with sensitivity to organelle-level metabolic responses, while EEA1, DAPI, and Ki67 performed best on recovering complex/association relationships, likely reflecting broader cellular reorganization and cell-cycle effects following perturbation of essential protein complexes (**Fig. 2f**). Consistently, including Golgin97 in panels typically improved performance in pathway-based settings but not complex-based ones (**Extended Data Fig. 1e)**, with the reverse observed for Ki67 (**Extended Data Fig. 1f)**.

Larger panels also resulted in more similar intra-perturbation profiles that were clearly distinct from those under other perturbations and controls (**Extended Data Fig. 2a**, ‘Phenoprint rate’). This indicates that increased plexity improves the ability to detect perturbation signals, while broadening the range of detectable responses. Nonetheless, performance varied markedly across random panels of the same size, underscoring the importance of marker choice. Because panels with high local similarity to the impact of the full panel also generally performed well against external ground truth (**Extended Data Fig. 2b**), we used this metric to further rank all panel configurations and identify markers associated with high performance. Among 1-round panels, top-ranked configurations were enriched for CAT, TOM20, Golgin97, fibrillarin, and pRPS6 (p-value < 0.05, KS test; **Fig. 2g**), several of which were not among the strongest single-channel performers. Together, these results suggest that higher plexity helps capture distinct types of relationships between perturbed genes and enables detection of unexpected and indirect effects. Increasing panel plexity broadens the range of effects that can be detected, a key goal of hypothesis-free morphological profiling.

Despite this overall gain, some marker-specific signals were obscured in the full panel embedding. In particular, 34.8% (72/207) of CORUM protein complexes with members in our perturbation library were recovered by at least one single-marker embedding but not captured by the full panel, and 26.6% (55/207) complexes fully captured by a single marker showed reduced recall with the full panel (**Extended Data Fig. 2c,d**). Thus, although multiplexing aids detection of perturbation effects and generally improves breadth of recovery, using the full profiles as the sole source of interpreting relationships can mask strong marker-specific signals.

### Archetype analysis and automated annotation reveal interpretable organelle morphological states

Because interpretation of perturbation effects requires relating morphological changes to meaningful biological concepts, we next devised a per-marker Archetype Analysis (AA)^28–31^ to define distinctive reference morphological states (archetypes) for each imaged cellular compartment (**Fig. 3a, Methods**) as part of the CHART framework. Unlike clustering, which enforces discrete partitioning, AA represents each cell as a convex combination of archetypes, allowing each cell to be described as a mixture of these reference states. For each marker, we generated single-cell embeddings, constructed a *k*-nearest neighbor (*k*NN) cell graph using the top embedding principal components (PCs), and aggregated similar cells into metacells^32,33^ to reduce noise. We then applied AA to identify the vertices of the polytopes encompassing each marker’s phenotypic space (**Fig. 3a**). These vertices (archetypes) represent clearly distinct morphological states for each marker that can be systematically annotated and serve as interpretable reference points. Within this framework, each cell is modelled as a weighted combination of the archetypes, reflecting its proximity to the reference states.

**Figure 3.**
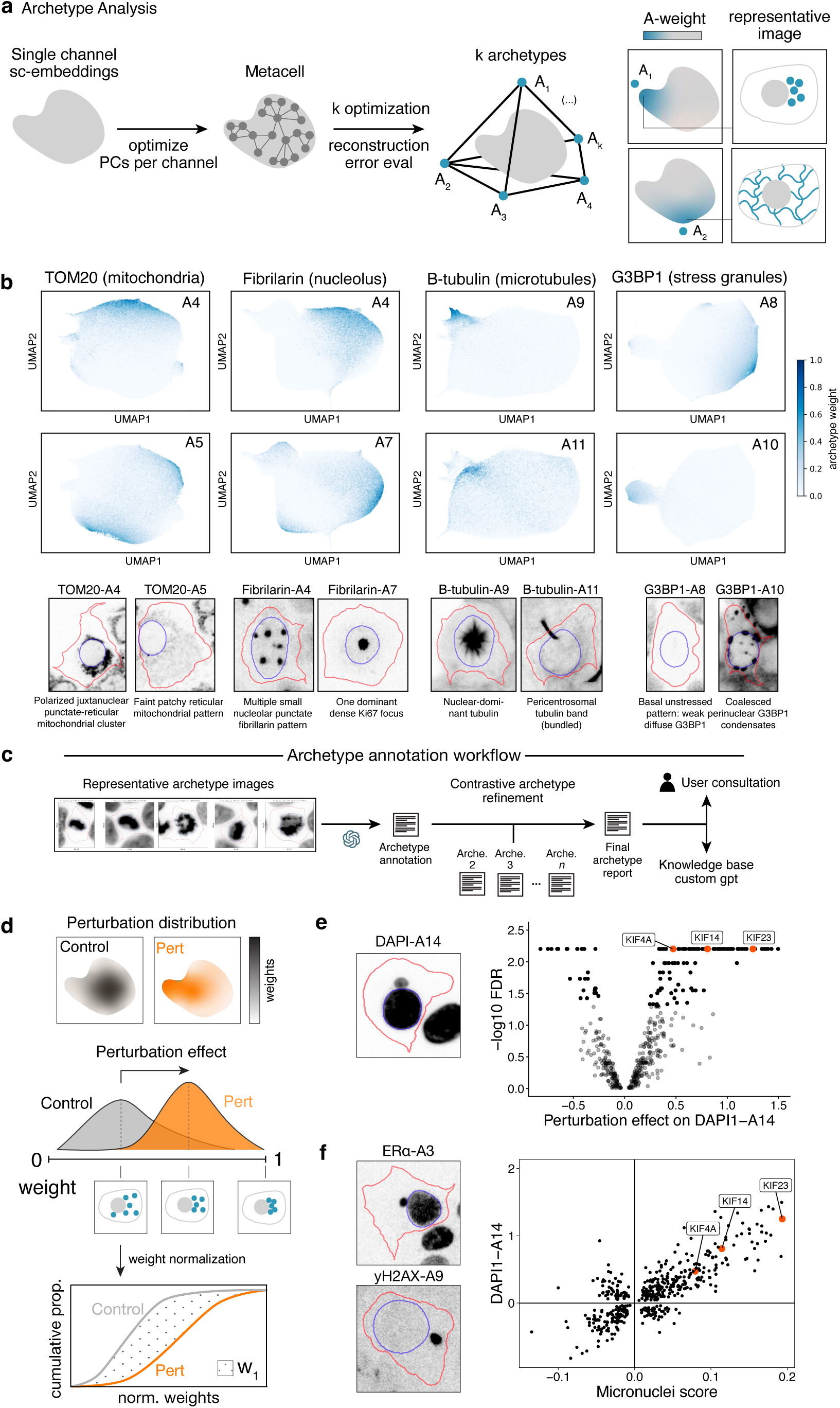
Archetype analysis and automated annotation reveal interpretable organelle morphological states. **a.** Archetype analysis (AA) approach. From left. A cell embedding was created for each marker (left), top PCs were selected (arrow), followed by aggregation into metacells (second left), and archetype analysis (second right) (**Methods**). Inspection of crops from cells with the heaviest archetype weights reveals characteristic marker phenotypes (right). **b.** Example archetypes from select markers. Single marker based embeddings of cell profiles (dots) (top two panels) colored by archetype weights (blue) and representative cell image selected from cells l among the top 10 with heaviest archetype weights (bottom) for two illustrative archetypes for each of TOM20 (mitochondria, left), fibrillarin (nucleoli, second left), B-tubulin (microtubules, second right) and G3BP1 (stress granules, right). **c.** Archetype annotation workflow. From left: images grids of high-scoring archetype cells for all markers were provided to a MLLM (GPT-5) pipeline that iteratively generated and refined structured textual archetype annotations through a contrastive comparison across other archetypes of the same marker. The resulting archetype reports can be used directly for analysis or as a knowledge base to create a customizable interactive LLM-based agents. **d.** Computation of perturbation effects on archetypes. Top: Archetype weights (color density) on perturbed (right, orange) and control (left, grey) cells. Middle: 1-D Wasserstein distance captures the overall displacement (magnitude and sign) between archetype weight distributions in the control and perturbed cells. The magnitude reflects the overall displacement between the two distributions. **e,f.** Regulators of the DAPI-A14 archetype reflecting micronuclei. **e,** Left: Representative image of DAPI-A14. Right. Effect size (x axis) and significance (-log_10_(FDR), permutation/Monte Carlo test, n = 1,000 iterations) of the impact of each perturbation (dot) on the DAPI-A14 archetype. **f.** Left: Representative images of ERɑ-A3 (top) and ɣH2AX-A9 (bottom). Right: DAPI-A14 perturbation effects (y axis) and effect on micronuclei (x axis; 1D-Wasserstein distance between perturbations and control cells for mean of channel-standardized weights per cell of 3 channels with micronuclei structures: DAPI, ERa and yH2Ax) for each perturbation (dot).

This approach yielded 188 archetypes across the 17 markers (8-14 archetypes per marker), excluding 31 archetypes corresponding to staining and segmentation errors (**Methods**). The retained archetypes captured meaningful morphological phenomena (*e.g.*, **Fig. 3b**), including changes in localization (*e.g.*, perinuclear mitochondrial accumulation, TOM20 A(rchetype)4; network expansion, TOM20 A5), puncta distribution (multiple small puncta, fibrillarin A4; single large aggregate, fibrillarin A7), distinctive structural states (mitotic spindles and intercellular bridges, β-tubulin A9 and A11, respectively), and altered marker accumulation (stress granule variance, G3BP1 A8 *vs*. A10). The global structure of perturbation relationships in the archetype space, defined as cosine similarity between perturbation profile pairs, was highly correlated to the perturbation relationship structure in the engineered feature space (Mantel r_M_ = 0.868, p < 0.001, **Extended Data Fig. 3a)**.

Because the markers in our panel are widely used and many of their associated phenotypes are well-characterized in the scientific literature, we automated the annotation of each archetype using multimodal large language models (MLLMs). We developed a multi-stage vision-language workflow that converted representative archetype images into standardized textual descriptions using appropriate biological terminology (**Fig. 3c**, **Supplementary Fig. 3**). Following an initial archetype annotation by the MLLM, our workflow ensured distinctive and informative descriptions by first contrasting images from each archetype against a baseline for that marker (defined as the 1-2 most dominant archetypes), with an additional side-by-side re-analysis in a new MLLM-iteration automatically triggered for semantically similar reports (**Methods**). This resulted in internally consistent descriptive archetype reports, linking AA-defined phenotypes to interpretable patterns of morphological variation (**Supplementary Fig. 4, 5**). For example, Fibrillarin-A7 was described as “a dominant nuclear punctate pattern characterized by a single, sharply circumscribed, intranuclear focus per cell (…) positioned centrally” (**Supplementary Fig. 5, Supplementary files)**. These reports can also serve as a knowledge base for other MLLM-based agents^34^, supporting interactive data exploration and rapid integration with external knowledge.

Next, using annotated archetypes as reference morphological states, we quantified how genetic perturbations shifted cell populations towards or away from those reference states. For each archetype, we quantified the difference between control and perturbed cell images using the 1-D Wasserstein distance on the Centered Log Ratio (CLR)-transformed archetype score, which can be interpreted as the area between the two empirical CDFs and thus measures the overall magnitude of distributional displacement (**Fig. 3d**).

Perturbations with well-established effects produced expected archetype shifts (**Extended Data Fig. 3b-e**). For example, loss of mTORC1 components (MTOR, RHEB, RPTOR, PDPK1) depleted the high-cytoplasmic pS6 A7 archetype and enriched for the low-signal state A1 archetype, consistent with reduced S6 phosphorylation following loss of mTORC1 signaling^35^. Deletion of HOPS complex member VPS18 promoted large aggregates of EEA1 archetype A7, reflecting expected disruption of early-to-late endosome transition and enlarged EEA1-positive endosomes^36^. G3BP1 knockout enriched for the weak diffuse staining G3BP1 A8 archetype, confirming protein depletion following knockout. Loss of the fusion factor MFN2 depleted the dispersed mitochondrial network archetype TOM20-A5, while enriching for compact juxtanuclear mitochondria TOM20-A7, consistent with increased mitochondrial fragmentation^37^.

This analysis allowed us to identify regulators of morphological phenotypes overlooked in traditional feature-based multichannel embeddings, such as DAPI-A14 (**Fig. 3e**, left), which captures micronuclei, small, extra-nuclear DNA-containing structures used as biomarkers of tumor grade and increasingly implicated in tumor progression^38,39^, which are generally associated with DNA damage and chromosome instability. Based on this archetype, we identified 106 functionally diverse genes whose perturbations promote micronucleus formation (FDR < 0.05) (**Fig. 3e**, right). Because micronuclei-like structures were also captured by archetypes for ERα (A3, **Fig. 3f**, top left) and γH2AX (A9, **Fig. 3f**, bottom left), we computed a composite ’micronuclei score’ by aggregating cell-level weights across all three archetypes. This score correlated with DAPI-A14 effect size (**Fig. 3f**, right: Spearman ρ = 0.75, P < 0.001), indicating joint regulation of these archetypes. The γH2AX association is consistent with localized genomic instability and chromothripsis, while ERα co-occurrence in micronuclei has not, to our knowledge, been previously described, and may warrant further investigation. The perturbation yielding the highest micronuclei score was KIF23, with KIF14 and KIF4A also among the high-ranked hits, possibly due to the role of KIFs in proper chromosomal segregation during mitosis^40^.

### Multi-compartment archetype analysis reveals coordinated morphological programs

Perturbations often cause shifts across multiple cellular archetypes (**Extended Data Fig. 4**). To identify recurrent and coordinated patterns of perturbation-induced archetype shifts, we decomposed the perturbation effect matrix using sparse principal component analysis (sPCA; **Fig. 4a**). This yielded 18 latent morphological programs (P1-P18), each defined by a sparse set of archetypes with positive and negative loadings and interpretable as a phenotypic axis combining distinct morphological states. Because these programs were derived from perturbation-level (pseudobulk) effects rather than individual-cell archetype scores, we next assessed whether their constituent archetypes shifted together within individual cells. Among cells at the extremes of each program-score distribution, archetypes belonging to the same program mostly showed concordant weight shifts, suggesting these programs capture co-occurring morphological changes in the same individual cells (**Extended Data Fig. 5, Supplementary Fig. 6**). The sparsity constraint produced compact programs that were well suited for MLLM-based annotation using the archetype loadings alongside the previously generated archetype reports (**Fig. 4b,c**).

**Figure 4.**
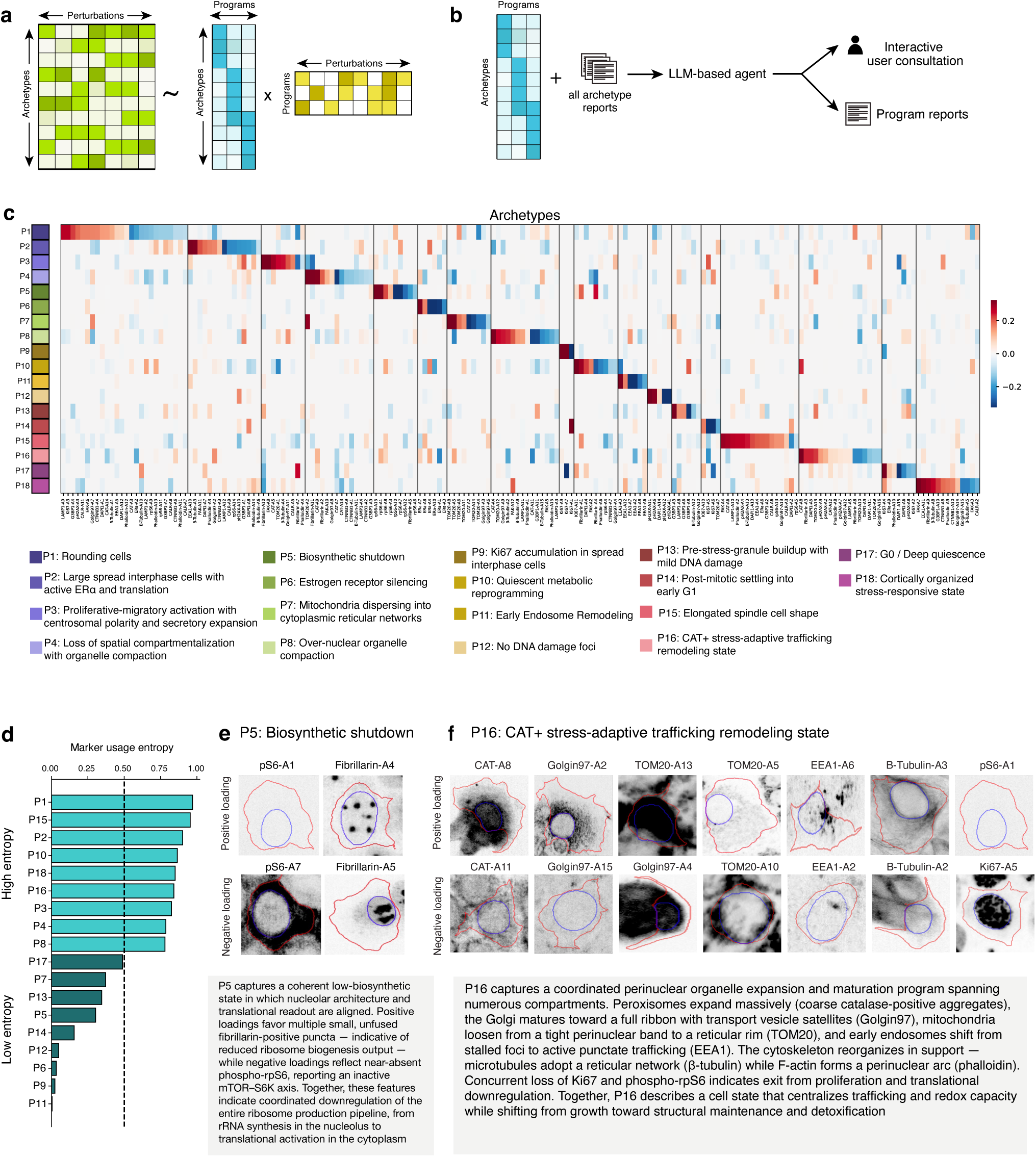
Coordinated morphological programs across compartments. **a.** Identification of morphological programs. Decomposition of the matrix of the effect of perturbations (left matrix, columns) on each archetype (left matrix, rows) (1D-Wasserstein distances between perturbed and control cells) using sparse PCA (sPCA), identifying latent morphological programs (middle matrix, columns) consisting of multiple coordinated archetypes (middle matrix, rows), and a regulation matrix of the impact of each perturbation (right matrix, columns) on each morphological program (right matrix, rows). **b.** MLLM-based annotation of morphological programs. Per-archetype visual reports and archetype-program weight matrices were provided to a knowledge-base LLM agent, which generated structured program annotations from a user-defined prompt. Annotations were refined through iterative user consultation with the agent. **c.** Morphological programs in our screen. Loadings (color bar) of each morphological program (rows) for each archetypes (columns). Bottom: program annotations. **d.** Marker usage entropy per program. Loadings were aggregated per marker and Shannon entropy was used to quantify high entropy (broad marker usage) and low entropy (few or single marker usage) programs (**Methods**). **e. ,f.** Example morphological programs. Representative images of top archetypes with positive and negative loadings (top) and automated (AI generated) textual descriptions for P5 (‘Biosynthetic shutdown’, e) and P16 (‘ERɑ+/CAT+ stress-adaptive trafficking remodelling state’, f).

While some programs were dominated by changes in a single or few markers (low Shannon entropy across channel-aggregated loadings, **Methods**), others involved coordinated co-variation across multiple markers (high entropy) (**Fig. 4d**). Low entropy programs included P7, which captures a shift from perinuclear to network-like mitochondria based on TOM20 (**Supplementary Fig. 7)**, and P11, which captures a shift in endosome distribution based on EEA1 (**Supplementary Fig. 8**). P5, another low entropy program, captured a decrease in phospho-S6 together with a shift in nucleolar architecture from fewer large nucleoli toward multiple smaller fibrillarin-positive puncta (**Fig. 4e, Supplementary Fig. 9**), linking reduced translational output and altered nucleolar organization. The nucleolus is the main site of ribosome biogenesis, and larger nucleoli are associated with greater protein-synthesis capacity^41–43^. This program, recovered solely through unsupervised image analysis and MLLM-based annotation, highlighted a link between nucleolar structure and translational activity that would typically be identified through biochemical assays. High entropy programs included P16, which captured coordinated changes across a broad set of compartments, including peroxisome accumulation (increase in CAT-A8, decrease in CAT-A11), mitochondrial network loosening (increase in TOM20-A5, decrease in TOM20-A10), Golgi maturation into a full ribbon (increase in Golgin97-A2, decrease in Golgin-A15/A4), maintained endosomal trafficking (increase in EEA1-A6), and cytoskeletal reorganization (increase in B-Tubulin-A3 and Phalloidin-A8, decrease in B-Tubulin-A2) supporting a perinuclear hub, alongside loss of proliferation (decrease in Ki67-A5), and translation (increase in pS6-A1) (**Fig. 4f**, **Supplementary Fig. 10**). P16 thus captures a cell state shifting from growth toward structural maintenance, active trafficking and detoxification. Overall, by integrating multicompartment changes within single programs, we capture complex morphological patterns and provide a framework for understanding perturbation effects.

Critically, several programs are directly regulated by perturbations in functionally coherent genes (**Fig. 5a**). For example, perturbation of mTORC-related genes (MTOR, RPTOR, PDPK1, RHEB, WDR24) increase P5 (translational shutdown); of γ-tubulin complex genes (TUBGCP6, TUBGCP3, TUBGCP4, TUBG1, TUBGCP2) increase P8 (marked by MTOC collapse); of translation initiation and G1/S regulators (EIF3B, EIF3A, EIF3I) increase P9 (interphase arrest) and of endosomal regulator (PIKFYVE) increases P11 (early endosome disorganization). This concordance between known gene function and morphological program identity indicates that morphological programs capture biologically meaningful responses.

**Figure 5.**
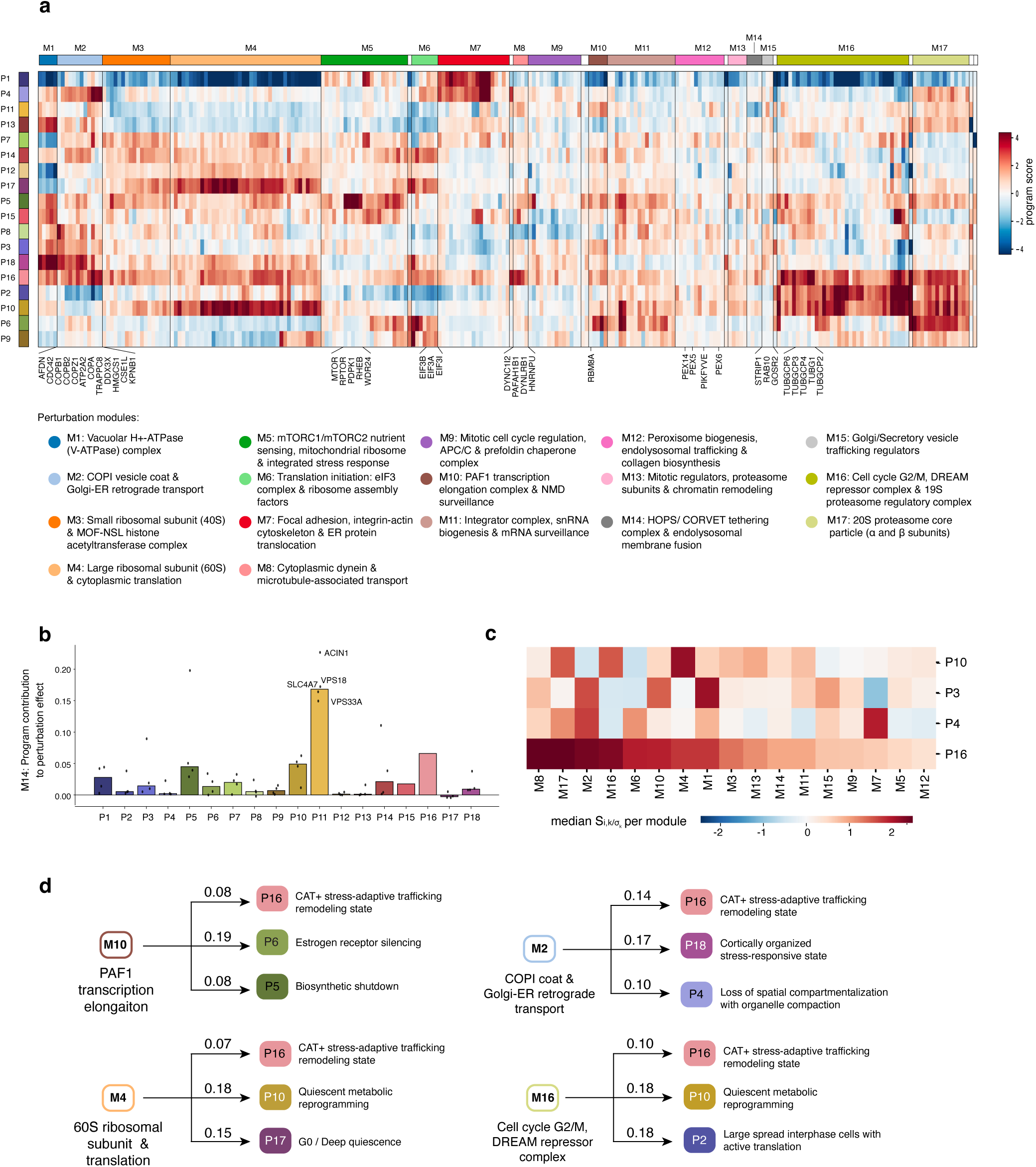
Morphological programs provide an interpretable framework for biological discovery. **a.** Regulation of morphological programs. Effects (color bar) of each perturbation (columns) on each morphological program (rows). Perturbations are clustered (top color bar) into modules using a consensus approach and annotated (**Methods**, bottom). **b.** Program contribution to the reconstruction of M14 perturbation effects. Program contributions (y-axis) reflect the share of perturbation effect that is attributable to each program. Contributions are shown individually for each gene in M14 (dots) and aggregated to the module median (bars). **c.** Effects of each perturbation module (columns) on subset morphological programs (rows) affecting similar sets of markers. Perturbation module scores were obtained by aggregating to the median across module members and scaled to program variance across modules. **d.** Schematic of morphological programs activated in conjunction with P16 by distinct modules. The top three programs activated by each module (M11, M2, M4, and M17) are shown, with the magnitude of the explained effect indicated above each module-to-program arrow.

### Morphological programs provide an interpretable framework for biological discovery

To define higher-order structure in our data, we clustered perturbations based on their morphological program scores (**Fig. 5a**), forming co-functional modules of perturbations with similar morphological effects. We quantified the share of morphological perturbation effect that is attributable to each program (‘program contribution’) using the archetype loadings and program scores and subsequently used both program contributions (**Extended Data Fig. 6a**) and scores to investigate gene function, link subcellular architectures to perturbation classes, and refine functional modules, which we illustrate with three key examples.

#### Morphological programs implicate ACIN1 in endosomal trafficking

Module M14 comprises KO of SLC4A7, ACIN1, VPS18, and VPS33A, and is characterized by a singular strong decrease (**Fig. 5a**) and contribution (**Fig. 5b, Methods**) from P11 (“Early endosome remodeling”, median contribution of 0.17 across M14 perturbations, a 2.55 ratio over the second top-contributing program P16 at 0.07). Thus, P11 is primarily defined by modulation of endocytic trafficking. VPS18 and VPS33A are members of the HOPS/CORVET complex, which regulates fusion in the endolysosomal system^44^, and SLC4A7 regulates intracellular pH and localize to endocytic membranes^45,46^. The presence of ACIN1 in M14, however, was more surprising, because it is mostly known as a peripheral factor of the exon junction complex (EJC), which binds mRNA upstream from exon-exon junctions regulating nonsense-mediated mRNA decay, alternative splicing and translation efficiency^47^. Its membership in M14 implicates a novel functional effect in modulation of endosomal trafficking, as reflected by the defining restructuring of the endosomal compartment captured by P11. While not previously described in mammalian cells, a *Drosophila* screen for endocytic trafficking defects identified the ACIN1 ortholog *Acinus* as an early endosome stabilization factor^48^. Together, these findings nominate ACIN1 as a candidate regulator of endosomal trafficking and raise the possibility that mechanisms linked to RNA processing or splicing intersect with endocytic control.

#### Cytoskeletal and trafficking perturbations drive distinct subcellular architectures through opposing morphological programs

Several multi-compartment morphological programs reveal how the same organelle repertoire can be reorganized into distinct spatial distributions with different functional implications (**Fig. 4d, Extended Data Fig. 6b**). For example, P10, P3, P4 and P16 all captured coordinated changes across peroxisomes, mitochondria and nucleoli among others, each resulting in distinct states with divergent functional implications (**Extended Data Fig. 6b**). In addition to the previously described P16, P10 is characterized by large cells with low proliferation and translation, an expansion in peroxisomes, active lysosomes and moderate mitochondria distribution, suggesting an exit from the cell-cycle shift towards a catabolic maintenance state (**Supplementary Fig. 11**). P4 is marked by loss of center-to-edge organization and transition from a spread, adherent morphology to a compact, rounded state with densely and uniformly distributed organelles, consistent with reduced spatial specialization during de-spreading or stress-adaptive remodeling (**Supplementary Fig. 12, 14**). By contrast, P3 defines a structured radial architecture in spread, adherent cells, with mitochondria, microtubules and peroxisomes arranged concentrically around the nucleus while FAK-positive adhesions and CTNNB1 junctions remain polarized at the cell edge (**Supplementary Fig. 13, 14**). This creates higher spatial modularity, combining a nucleus-encircling organelle-rich zone with a peripheral adhesion-signaling domain. Thus, while P10 and P16 are largely marked by expansion or contraction of select markers, P3 and P4 capture broad re-positioning of multiple organelles simultaneously.

The fundamentally distinct morphological states captured in P3 and P4 are in turn driven by distinct co-functional modules, linking pathway functions to specific subcellular spatial outcomes (**Fig. 5c**). M7 (focal adhesion and integrin-actin cytoskeleton) activates P4 (**Fig. 5c, Extended Data Fig. 6c**). Thus, adhesion and cytoskeletal pathways not only impact migration, but are also necessary to provide the structural scaffold upon which subcellular spatial specialization depends. Their disruption collapses cells into the spatially homogeneous P4 state, altering not only motility but the functional independence of organellar signaling domains. In contrast, M1 (V-ATPase complex) preferentially activates the centrosome-anchored P3 architecture (**Fig. 5c, Extended Data Fig. 6c**). V-ATPases regulate luminal acidification of multiple compartments including lysosomes, endosomes and Golgi-derived and secretory vesicles^49^. Moreover, genetic variants of its subunits are associated with congenital disorders marked by altered Golgi function^50^. This suggests that impaired acidification, and resulting abnormal membrane traffic^51,52^ may promote organelle compaction around the nucleus while peripheral adhesion signaling persists. Thus, normal trafficking may be required to maintain organelle distribution across multiple subcellular zones. In support of this, M15, a small module comprising regulators of secretory traffic, including GOSR2^53^ and RAB10^54,55^, also preferentially activated P3 (**Extended Data Fig. 6c**).

Together, these results demonstrate that subcellular spatial organization is actively maintained by the balance between cytoskeletal anchoring and vesicular trafficking; the former establishes the spread morphology that permits spatial segregation and organelle anchoring, while the latter distributes organelles across it. Perturbations to either system reshape the same organelle repertoire into architectures with fundamentally different structure-function relationships.

#### Combinatorial morphological programs reveal shared and divergent adaptation states

Co-functional modules often combine multiple morphological programs that enable tailored cellular response. For example, multiple modules active the P16 trafficking stress-adaptation program (**Fig. 5c**), but each also regulated a combination of other programs, yielding a distinct phenotypic response overall. For instance, the perturbations in M10 (PAF1 Complex mediated transcription) coupled P16 to a decrease in ERa levels (P6) and biosynthetic shutdown (P5), whereas those in M2 (Golgi-ER retrograde transport) promoted broader cellular re-organization (cytoskeletal changes in P18, and P4 to a lesser extent) (**Fig. 5d**, **Supplementary Fig. 9, 12, 15, 16**). Modules involving perturbations in 60S ribosome (M4) and cell-cycle and proteostasis (M16) genes were more similar, both combining P16 with the cell-cycle-exit and catabolic metabolic reprogramming of P10 (**Fig. 5d, Supplementary Fig. 11**). M4 in addition further activated P17 (**Supplementary Fig. 17)**, a quiescence-like program, with features present in either quiescent or senescent cells, such as absence of Ki67^56,57^, and a single nucleolus^58–60^. M16 instead activated P2 (**Supplementary Fig. 18**), which captures large spread interphase cells, but with organized cytoskeleton, active lamellipodia and high biosynthetic capacity. Thus, both perturbation modules lead to exit from the cell cycle (P10), rely on adaptation features of trafficking (P16) and metabolic reprogramming (P10), but ribosomal perturbations (M4) activate quiescent features and cell-cycle perturbations (M16) lead to more actively signaling interphase cells with organized cytoskeleton. This provides insight into both shared and diverging mechanisms of cellular adaptation.

Together, these examples demonstrate that morphological programs, by decomposing complex phenotypes into modular and interpretable axes, link genetic perturbations to specific subcellular architectural outcomes that would be obscured in aggregate analyses, generating mechanistic hypotheses with direct relevance to disease-associated processes.

## DISCUSSION

Our study addresses two current limitations of Optical Pooled Screening: limited understanding of how individual markers contribute to morphological embeddings, and limited ability to translate high-dimensional morphological profiles into interpretable biology. By combining highly multiplexed data from CellMAPP with systematic *in silico* panel subsetting, we show that markers enable detection of distinct types of perturbation-relationships and, when combined, help broaden the range of information recovered. CHART leverages this rich signal by resolving imaging features into distinct marker-level morphological archetypes, linking those archetypes to biological interpretation through MLLM-based annotation, and integrating them via sparse PCA into morphological programs that capture coordinated phenotypic changes across markers. Together, this enables identification of regulators of defined organelle states and mapping of co-functional gene modules to defined morphological changes that support mechanistic interpretation, transforming high-dimensional image features from ’black box’ embeddings into interpretable descriptions of what changed, where it changed, and what changes co-occur.

These biological and interpretive gains relied upon a robust preprocessing framework. OPS feature tables are especially sensitive to segmentation errors, registration failures across staining rounds and well-level technical effects^61–63^, all of which can distort downstream embeddings. Our pipeline combined object- and feature-level filtering with quantile normalization rather than z-scoring. Although such steps are often treated as secondary to downstream modeling, they can substantially affect biological recovery and should be carefully considered during data analysis.

Our *in silico* panel-subsetting analysis addressed a key practical question for OPS: how marker choice and panel complexity shape the biological signal recovered by morphological profiling. Marker performance varied across evaluation metrics, markers were frequently sensitive to perturbations beyond those directly targeting the structures they imaged, and differed considerably in the types of biological relationships they recovered (*e.g.*, complex *vs*. pathway-based). This motivates use of higher-plexity panels that capture a broader range of effects, without requiring them to be defined *a priori*, an important consideration for hypothesis generating screens. Notably, some smaller panels performed exceptionally well (**Supplementary Table 3**), suggesting compact high-performing assays may be achievable. While optimal panel composition requires investigation across a broader range of models and conditions, our results indicate they should, at a minimum, include markers sensitive to complementary biological relationships (e.g. Ki67 and TOM20/Golgin97). Moreover, panel design could benefit from exploration of a broader marker space, since markers such as CAT and pRPS6 (rare in standard panels like Cell Painting) were enriched in top-performing minimal panels.

The added interpretability provided by CHART made it possible to resolve both discrete organelle phenotypes and coordinated cross-compartment responses. For example, CHART isolated micronucleus-associated states that were difficult to recover from aggregate multichannel features alone. Since micronuclei have recently been implicated in suppression of extrachromosomal DNA (ecDNA)^64^, a key feature of ER⁺ high-risk patients^65^, the regulators identified here provide candidates for testing whether micronuclear sequestration influences ecDNA activity in ecDNA-positive breast cancer models.

While coordinated changes across subcellular compartments are poorly characterized, as experiments often focus on one or few structures at a time, the morphological programs we identified using this highly multiplexed data reveal novel patterns of cross-compartment organization (e.g. coordinated changes across peroxisome, mitochondria, Golgi and endosomes in P16). Since both peroxisomes^66–69^ and membrane trafficking^70,71^ have been implicated in cellular adaptation to numerous stressors, P16 suggests that cells frequently couple changes in these systems to better handle and process altered signals. Our analysis also uncovered an unexpected link between ACIN1 and endosomal trafficking, demonstrating that similar organelle repertoires can be reorganized into distinct spatial architectures with different, likely functional, consequences. Thus, CellMAPP and CHART enable tracking of cross compartment alteration at scale, mapping how perturbations reorganize the cell’s physical components into functional states.

Modular combinations of morphological programs also provided insight into distinct mechanisms of cellular adaptation. Disruption of the ribosomal machinery (M4) and of cell-cycle regulators (M16) both relied on metabolic, oxidative, and trafficking changes (P10, P16), but the former additionally led to a decreased biosynthetic capacity and shift toward a quiescent-like state (P17), while the latter maintained a more active cell state (P2). The pathways captured by these modules are the targets of approved and investigational cancer therapies, including CDK inhibitors, translation-initiation/eIF inhibitors, and ribosome-targeting agents^72–74^, but intrinsic and acquired resistance limits durable responses^72,75^. The distinct adaptation modes we identify may help define context-specific resistance mechanisms and inform rational combination strategies.

Because CHART operates on marker-resolved phenotypes rather than a specific feature representation, it should be transferable across cell lines, marker sets and future vision-based embeddings. It should also improve as MLLMs continue to advance. Similarly, the CellMAPP panel was largely designed around cellular structures present across most cell types, supporting its use in additional models and conditions. However, several limitations remain. Our library was enriched for genes expected to produce strong morphological phenotypes and all experiments were performed in one cell type; as such, its performance across a broader context remains to be assessed. In addition, the morphological programs identified are based on perturbation-level aggregate effects and thus do not resolve heterogeneous responses within perturbations. Future analysis can learn such programs from single cell weights for a more direct capture of co-occurring morphological changes. Even so, these results establish a practical framework for moving optical pooled screens beyond phenotypic similarity maps toward interpretable cell biology.

## METHODS

### Cell culture

MCF-7 cells were obtained from ATCC and were maintained in DMEM supplemented with 10% fetal bovine serum (FBS; vol/vol), 100 U/ ml penicillin, 100 μg/ml streptomycin and 1× GlutaMAX. A doxycyclin-inducible Cas9 cassette was introduced into MCF-7 cells by piggybac transposon, followed by blasticitin selection. Single-cell sorting was performed and, after clonal expansion, knockout efficiency was evaluated by transducing CD81 guide and amplicon PCR of CD81 genomic loci. A clone with high knockout efficiency and high consistency with the parental line’s growth rate and estrogen signaling activity was selected for further screening (**Supplementary Fig. 1c,d)**.

### Lentiviral transduction

For virus production HEK293T cells were plated in 6-well plates at a density of 1 million cells per well. The following day, cells were transfected with the expression plasmid, delta8.9 and pCMV-VSV-G at a molar ratio of (1:2.3:0.2) using Lipofectamine 3000 (Thermo Fisher Scientific, L3000015). Opti-MEM (Gibco, 31985062) was exchanged to culture medium after 6 h. Viral supernatant was collected 2 days after transfection and filtered through a 0.45-μm filter.

MCF-7-Cas9 (A08) cells were transduced with the appropriate volumes of lentiviral supernatant overnight to achieve a Multiplicity of Infection (MOI) of <0.2. Cells were selected with 2 μg/mL puromycin (Gibco, A1113803) for 3 days. Cas9 expression was induced with 2 μg/mL doxycycline for 3 days, and the cell library was seeded into glass-bottom plates that tolerate high heat (Cellvis, P061.5HN) at a density of 300,000 cells per well 48 hours prior to fixation in media still containing doxycycline.

### Imaging

Imaging was performed on a Nikon Ti2 microscope equipped with Nikon lenses, a Teledyne Photometrics Kinetix sCMOS and a Lumencor CELESTA light engine. IBEX immunofluorescence images were acquired with a ×20/0.75-NA CFI Plan Apo Lambda objective with the following conditions: DAPI (365 nm, FF01-441/511/593/684/817), CF430 (446 nm, excitation 432/523/702, emission FF01-470/28, dm 459/526/596), GFP (477 nm, FF01-511/20), AF532 (545 nm, excitation FF01-432/523/702, emission FF01-563/9, dm 555-Di03), AF647 (637 nm, FF01-680/42) and AF750 (748 nm, emission FF01-792/64).

ISS cycles were imaged using a ×10/0.45-NA CFI Plan Apo Lambda objective with the CELESTA-DA/FI/TR/Cy5/Cy7 excitation filter and FF01-391/477/549/639/741 dichroic filter (Semrock) unless noted otherwise. Laser excitation wavelength and emission filters were set up for each channel: DAPI (408 nm, FF01-441/511/593/684/817), base G (545 nm, FF01-575/19), base T (545 nm, FF01-615/24), base A (637 nm, FF01-680/42) and base C (637 nm, FF01-732/68).

### Multiround immunofluorescence

Cells were fixed with 4% PFA (Electron Microscopy Sciences, 15714) in PBS for 30 min and washed three times with PBS with 0.05% Tween (PBST). Cells were then permeabilized with 0.3% Triton X-100 in PBS for 10 min, washed three times with PBST and blocked with 2% BSA in PBS for 1 h at room temperature. Cells were stained with fluorescent conjugated antibodies for 1h at room temperature, followed by three washes with PBST. Immunofluorescence was repeated for four cycles using IBEX^20^ iterative bleaching for signal removal. Complete signal removal was confirmed between rounds and staining order was optimized such that antibodies that showed greater change in signal after successive bleaching rounds were included in earlier rounds (1-2), and those which displayed minimal change in signal bleaching were included in later rounds (3-4). Antibodies used are listed in Supplementary Table 4.

### ISS

The PerturbView *in situ* sequencing (ISS) protocol was used, as previously described^21^. Specifically, cells were permeabilized with 70% ethanol for 30 min, followed by 75% volume exchange with PBST three times (to prevent drying) and two complete PBST washes. Next, heat decrosslinking was performed in 0.1 M sodium bicarbonate pH to 8.5 and 0.3 M NaCl in water at 65 °C for 4 h. After three washes with PBST, *in vitro* transcription (IVT) was conducted using a HiScribe T7 Quick High Yield RNA Synthesis kit (New England Biolabs, E2050S), 1× T7 RNA Polymerase Mix, NTP Buffer Mix at 10 mM and the addition of RiboLock RNase inhibitor (Thermo Fisher Scientific, EO0384) at 0.4 U /μl at 37 °C for 12 h. A primer cross-linking amine-modified primer was hybridized at 1 μM in PBST for 30 min at room temperature. After the hybridization and before reverse transcription, cells were fixed with 3.2% PFA and 0.1% glutaraldehyde (Electron Microscopy Sciences, 16120) in PBST for 30 min. The reaction was quenched by adding 0.2 M Tris-HCl (pH 8), and cells were washed three times with PBST. The reverse transcription reaction mix (1× RevertAid RT buffer, 250 μM dNTPs (New England Biolabs, N0447L), 0.2 mg /ml recombinant albumin (New England Biolabs, B9200S), 1 μM reverse transcription primer, 0.4 U /μl Ribolock RNase inhibitor and 4.8 U /μl RevertAid H minus reverse transcriptase (Thermo Fisher Scientific, EP0452)) was added to the samples, followed by incubation at 37 °C overnight. After reverse transcription, cells were washed three times with PBST and fixed again in 3.2% PFA and 0.1% glutaraldehyde in PBST for 30 min. Cells were then washed with PBST six times to remove residual dNTPs. After this step, cells were incubated in gap-fill reaction mix (1× Ampligase buffer, 0.4 U /μl RNase H (Enzymatics, Y9220L), 0.2 mg /ml recombinant albumin, 100 nM padlock probe, 0.02 U /μl TaqIT polymerase (Enzymatics, P7620L), 0.5 U /μl Ampligase (Lucigen, A3210K) and 50 nM dNTPs) for 5 min at 37 °C and 90 min at 45 °C. Cells were washed with PBST three times, and rolling circle amplification was performed by adding the reaction mix (1× Phi29 buffer, 250 μM dNTPs, 0.2 mg /ml recombinant albumin, 5% glycerol and 1 U /μl Phi29 DNA polymerase (Thermo Fisher Scientific, EP0091)) and incubating at 30 °C for 24 h. ISS steps were conducted according to the OPS protocol^76^ with the primer sequences listed in Supplementary Table 5.

### Library construction

For the library, four sgRNAs were designed per gene for 524 genes of interest, 50 olfactory receptor genes (as non-expressed gene controls), and another 20 different nontargeting sgRNAs. Spacer sequences were designed to be resolvable by eight cycles of ISS using an internal implementation of crisprVerse^77^. The oligonucleotide pool was synthesized by Twist Bioscience, amplified by dial-out PCR using Ex Premier DNA Polymerase (Takara Bio, RR370A), cloned into a PerturbView vector^21^ with Golden Gate assembly using BsmBI (New England Biolabs, E1602S), transformed into Stellar electrocompetent cells (Takara Bio, 636765) and packaged into a lentivirus library while maintaining >1,000× coverage at every step. Guide sequences are listed in Supplementary Table 6.

### Image preprocessing: registration, segmentation, barcode calling and feature extraction

First, raw image tiles were corrected for non-uniform illumination using the algorithm described by Singh et al^78^. Images from each well and cycle were then stitched into whole-well images using ASHLAR (v0.18)^79^. SCALLOPS^22^ was used for all subsequent processing steps. Whole-well images were aligned into a common coordinate system across cycles and across magnifications, using image registration to maximize similarity of corresponding features and correct stage jitter and objective-dependent misalignment.

To extract relevant information from cells and nuclei, image segmentation was performed in two sequential steps. Nuclei were first segmented using SCALLOPS StarDist wrapper with StarDist v0.9.1 and default StarDist inference parameters, and the resulting nuclei served as seeds for a second step of cytoplasmic segmentation using propagation. Segmentation information was stored in Zarr format.

For ISS analysis, spot detection was performed to identify candidate ISS foci. Aligned images were transformed using a Laplacian-of-Gaussian filter with σ=1. The resulting filtered images were then subjected to a max filter with a neighborhood size of 3 pixels. Subsequently, the standard deviation across cycles was calculated, followed by the mean across channels. The resulting array was used to identify local maxima within a 5 × 5 pixel neighborhood. These maxima represent candidate ISS spot locations that are later decoded during base calling. Spot intensities were extracted and corrected for cross-talk by linear transformation. Reads were then identified by calling the base with the highest intensity at each candidate spot in each cycle, with per-base quality scores calculated from the softmax probability of the brightest base.

A table of cellular, cytoplasmic, and nuclear phenotypes was generated based on the previously segmented phenotypic images. Features were extracted across four categories: (1) intensity features capturing the distribution and magnitude of signal, including percentiles (25th, 75th), median absolute deviation, maximum, median, mean, standard deviation, total intensity (sum), and radial intensity distributions from the centroid; (2) texture features quantifying spatial patterns of pixel intensities using Haralick descriptors (contrast, dissimilarity, homogeneity, angular second moment, correlation, energy) computed from gray-level co-occurrence matrices; (3) geometry features describing morphological properties including area, perimeter, eccentricity, solidity, convexity, circularity, aspect ratio, and seven Hu moments providing translation, scale, and rotation-invariant shape descriptors; and (4) multi-channel features characterizing relationships between fluorescent channels through Pearson’s and Spearman’s correlation coefficients and Manders’ colocalization coefficients (M1, M2).

ISS results were merged with phenotypic imaging data based on their common cell label. The resulting table contained all the features listed above for each channel of the phenotype image, along with the label of each cell, well identity, barcodes identified and barcode-associated information.

Different development versions of SCALLOPS (pre-release) were used for different steps of the image processing pipeline. ISS registration across cycles and nuclear segmentation were run with SCALLOPS pre-relase v.0.0.1. Cell segmentation, label transfer, spot detection and read calling were run with SCALLOPS pre-release v.0.0.2. Phenotypic registration across rounds and feature extraction were run with SCALLOPS pre-release v.0.0.3.

### Quality control of engineered features

Tabular engineered features obtained from SCALLOPS were subjected to quality control aiming to remove segmentation and registration errors. First, obvious segmentation errors were removed by setting a maximum threshold for cytosol / nuclear area of 15. Next, registration quality was assessed as the correlation between DAPI intensity values across rounds, since all 4 phenotype rounds and all 8 sequencing rounds share a DAPI stain. Phenotype registration quality was assessed as the correlation between DAPI intensity in round 1 to intensities in rounds 2-4 within each cell’s bounding box. Cells with an average correlation across rounds < 0.8 (5.8% of total cells with mapped barcodes) were discarded. Phenotype-to-sequencing registration quality was similarly assessed as the DAPI intensity correlation between the last phenotype round and the first sequencing round. Cells with correlation < 0.8 (43.1% of total cells with mapped barcodes) were discarded. All QC steps were performed for each well separately.

### Preprocessing of engineered features

After QC, cells across all 6 wells were concatenated and quantile normalization was performed to correct batch effects related to well ID. To this end, the distribution of each feature in each well was quantile normalized to the distribution of concatenated values across all 6 wells, using linear interpolation to account for differences in distribution size. After normalization, low-variance (robust CV < 0.01) and NA-rich (> 100,000 missing values) features (3.4% of total 11,804) were removed. Cells with missing values after these feature filters (1.08% of the remaining cells after per-well QC filters) were removed. Outlier detection was then performed across cells receiving the same gene perturbation, using Isolation Forest^80,81^ (*IsolationForest* function from *sklearn* Python package with *n_estimators=100* and *contamination=’auto’*).

Next, normalized feature profiles were aggregated (median) across all the cells receiving the same guide and outlier guide detection was performed. First, data were subsetted to control guides (NTC or targeting olfactory receptors (ORs)). Then, outlier detection was performed using Isolation Forest as previously described, as well as Local Outlier Factor^82^ (*LocalOutlierFactor* function from *sklearn* Python package with *n_neighbors=20* and *contamination=’auto’*) and Elliptic Envelope^83^ (*EllipticEnvelope* function from *sklearn* Python package with default parameters) on the PCA space derived from the aggregated features (PC kept up to 90% variance explained). Guides were considered outliers if identified by 2 out of these 3 methods. After outlier guides were removed (5.9% of control guides), the remaining guides targeting ORs and NTCs were retained as controls for downstream use. Cells from control guides were used downstream as ‘control cells’.

Perturbation guide outliers were identified by calculating the cosine similarity between their pre-processed feature profiles and those of other guides targeting the same gene. Firstly, the PCA space of all guides was centered on the previously identified control guides. Then, pair-wise cosine similarity was computed between the profiles of all guides targeting the same gene. Guides with mean cosine similarity to others < 0.2 were excluded (21.4% of perturbation guides). Finally, guides with < 50 cells were excluded (2.6% of remaining control and perturbation guides).

Overall, 874,882 cell profiles across 1,595 perturbation and 207 control guides were retained for downstream analyses.

### Gene-level embedding and clustering

After preprocessing, cell-level feature quantifications were aggregated (median), first across all cells receiving the same gRNA and then across all gRNAs targeting the same gene. Incremental PCA^84^ (*IncrementalPCA()* function from *sklearn* Python package with batch_size=2,000) was then run on these pseudo-bulked profiles, and PCs explaining the first 90% of the variance in the data were retained.

Leiden clustering was performed on the *k*-NN graphs built from the PCA space, using *scanpy* functions *sc.pp.neighbors()* with *nn=5* and *sc.tl.leiden()* with *res=2*.

### Panel evaluation by sub-sampling

#### Local similarity to full panel

Local similarity between a query embedding for a subset of the panel and a reference embedding of the full panel was quantified using the Area Under the Recall Curve (AURC). To generate each embedding (reference or query), a *k*-nearest neighbor (*k*-NN) graph was constructed in the PCA space using a Euclidean distance, implemented via *scanpy.pp.neighbors* with PCs explaining 90% of total variance. *k*-NN graphs were computed up to a maximum neighborhood size *k_max_=30*. For each cell *i* and each neighborhood size *k = 1, …, k_max_*, neighborhood recall was defined as the fraction of each cell’s *k* nearest neighbors in the reference (full panel) graph that are also among its *k* nearest neighbors in the query graph. The AURC for each cell was then computed as the area under the recall-*vs*-k curve using trapezoidal integration (*numpy.trapz*), normalized by the maximum possible area (i.e., perfect recall at all *k*) yielding a score in [0, 1], where 1 indicates perfect neighborhood preservation.

To assess robustness to the choice of similarity metric, this analysis was also performed using cosine similarity instead of Euclidean distance with *scanpy.pp.neighnbors(metric=’cosine’, …)*, resulting in a similar increasing trend across subsampled panels (**Supplementary Fig. 19**).

#### Recovery of ground truth relationships

Evaluation of ground truth relationship recovery followed the recall computation method described by Celik *et al.*^85^. Briefly, pairwise cosine similarity was computed between the mean profiles for all perturbed genes. Relationships below the 5th and above the 95th similarity percentiles of the distribution of all pairwise similarities (excluding self-links) were selected as ‘predicted links’. Ground truth associations between genes were then obtained from different databases: CORUM^26^, StringDB^25^ (no confidence filters), HuMAP^27^, SIGNOR^24^ and Reactome^23^. Recall at 95.0 was computed as the ratio between predicted true links over all links. AURC was computed as the average of recall at similarity thresholds 95.0, 96.0, 97.0, 98.0 and 99.0.

#### Confounding variable evaluation

The single-cell profiles in PCA space were aggregated to the median for each guide-well combination, and batch effects were computed as the Kolmogorov-Smirnov test statistic (*ks*) between two distributions: a foreground distribution of cosine distances in PCA space between pairs within the same batch (well) and a background distribution of distances in PCA space between randomly selected pairs. Low *ks* values indicate low separability of wells in the PCA space, *i.e.* successful batch correction.

#### Perturbation effect consistency evaluation

Single-cell profiles in PCA space were aggregated to the median for each guide-well combination, and perturbation effect consistency was computed as the Kolmogorov-Smirnov test statistic (*ks*) between two distributions: a foreground distribution of cosine distances in PCA space between pairs receiving the same perturbation (gene or guide) and a background distribution of distances in PCA space between randomly selected pairs. High *ks* values indicate stronger similarity of profiles receiving the same perturbation compared to random pairs, *i.e.* consistent perturbation effects.

#### Phenoprint rate evaluation

Phenoprint rate evaluation adapted the perturbation signal magnitude computation method described by Celik *et al.*^85^. Single-cell profiles in PCA space were used to compute leave-one-out average cosine similarity, defined as

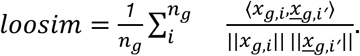

where *x_g_*_,*i′*_represents the average of all except the *i*th profiles.

Foreground similarities were computed among cells receiving the same guide (or gene) perturbation, and background similarities were computed among a random sample of control cells matching the number of perturbed cells. Statistical significance was calculated as the fraction of times foreground similarities were higher than background similarities across 10,000 repeats. Final perturbation (gene or guide level) results were obtained by aggregating to the median across wells.

#### Per-marker archetype analysis

The complete feature table was subsetted to contain features extracted exclusively from each marker, excluding geometry features that are not marker specific, such as cell and nucleus size. Embeddings were generated from this subset feature matrix as above (incremental PCA and retention of PCs explaining 90% of variance). The PC space was then further reduced to the top PCs using the elbow method, and then aggregated to the metacell level using the function *SCimplify_from_embedding()* from the SuperCell R package, with *k.nn=10* and *gamma=30* for metacell generation. Archetype analysis was performed across 8 metacell subsets of 12K metacells each, generated by enforced sampling of the full metacell space via Farthest-Point Sampling and enforced sampling from sparse regions via weighing of sampling probabilities by metacell rarity. For each subset, archetypes were generated using the function *stepArchetypes()* from the archetypes R package, with *k* ranging from 6 to 20. For each *k* in each subset, metacell reconstruction error on an evaluation set was computed, and the final number of archetypes was decided based on the RMSE trend over *k*.

#### Comparison of perturbation relationships between the engineered feature and archetype spaces

Perturbations were filtered to exclude the control cluster identified in the engineered feature space (cluster 3). Pairwise cosine similarity was computed both in the engineered feature space (PCA of gene-level aggregated profiles, centered on control genes, as described above) and the archetype space. Archetypes corresponding to artifacts (as judged by visual inspection) were filtered, cell weights per channel were re-weighted to sum to 1 and mapped into the Euclidean space with an isometric log-ratio (ILR) transform. Per-channel ILR-transformed weights at the single-cell level were median-aggregated to gene level and centered on control genes, following the same procedure as above. Cosine similarity matrices in these two spaces were compared using the Mantel test. For visualization purposes, perturbations were clustered on both similarity matrices as follows: first, each similarity matrix was z-transformed over all entries. The average over these two matrices was then used for hierarchical clustering with average linkage. The hierarchical clustering results were used to order perturbations in the split heatmap of **Extended Data Fig. 4**, where raw cosine similarity values were plotted.

#### Automated image annotation

Morphological archetypes were annotated using a multi-stage MLLM-based visual reasoning pipeline implemented in Python using LangChain and the OpenAI Azure API with GPT-5. All prompts used in each step are provided in supplementary files.

For each archetype, the 10 cells with highest membership weights were extracted and arranged into 5-cell grid images with overlaid cell and nuclear boundaries. Images were analyzed through a four-stage process. First, each grid was independently analyzed with GPT-5 vision using structured prompts restricted to directly observable features, including fluorescence intensity, intracellular localization, signal texture, cell and nuclear shape, and technical artifacts. Second, contrastive baseline analysis was performed hierarchically: each archetype was compared against the 1 or 2 most abundant archetype for that marker (**Supplementary Fig. 20**), which approximates baseline staining in the cell line used as assessed by manual inspection. This generated per-archetype notes describing relative differences in staining intensity, localization, texture, size, shape, and background signal. Third, archetype-level reports were synthesized with a prompt in a new MLLM iteration, which combined the representative grid-level annotation with the corresponding baseline comparison note to produce a structured summary, candidate archetype names, and a functional hypothesis. Fourth, semantic similarity among the initial archetype summaries was assessed using SentenceTransformers (all-MiniLM-L6-v2). For each marker, summary embeddings were compared pairwise using cosine distance, and archetype pairs with cosine distance < 0.1 were subjected to targeted bidirectional side-by-side comparison. The resulting contrastive notes were used to refine the summary section. Step-specific reasoning settings for GPT5 were used to balance descriptive consistency in step 1 and 2 (reasoning level ‘low’) and higher-level synthesis in step 3 (reasoning level ‘medium’) and step 4 (reasoning level ‘high’). Completion length was capped at each stage to limit verbosity and token use.

Across all stages, prompts instructed the model to avoid unsupported biological inference during primary visual description, to separate true cellular signal from background or neighboring-cell signal, and to report uncertainty when visual distinctions were ambiguous.

Pipeline outputs were manually reviewed by a human scientist for accuracy of the visual descriptions and candidate archetype labels. The pipeline generated three candidate names for each archetype. The most appropriate name was manually selected and used in the final archetype dashboard and downstream analyses. Overall, 17/188 of archetypes were found to be incorrectly described and were subsequently corrected. Mis-annotations most commonly occurred in cases with extreme or largely homogeneous staining patterns, including very high fluorescence intensity, largely absent signal, elevated background, or apparent spillover between nuclear and cytosolic compartments, where compartment-specific signal assignment was visually ambiguous. Representative examples of corrected annotations are shown in **Supplementary Fig. 21**.

#### Perturbation effect quantification

Archetype weights per cell were CLR-transformed across archetypes for each marker and concatenated across markers. Archetypes associated with segmentation or staining errors were identified through visual inspection and removed. Cells from each perturbation were compared to control cells previously identified, using 1-D Wasserstein distance (w_1_). Empirical p-values were calculated by re-shuffling perturbation labels (n=1,000). For quantification of aggregate effects of perturbations over groups of archetypes (e.g., micronuclei), CLR-transformed archetype weights were z-transformed for each marker and aggregated to the mean. Perturbation effects were computed on this vector using 1-D Wasserstein distance as above.

#### Identification of morphological programs

The matrix of the effect of each perturbation on each archetype W (n_perturbations_ × n_archetypes_) was centered by archetype and factorized using sparse PCA with scikit-learn *SparsePCA*. Sparse PCA decomposes W ≈ S L^T^, yielding a score matrix S (n_perturbations_ × K) and a sparse loading matrix L (n_archetypes_ × K), where K is the number of latent morphological programs. Each morphological program corresponds to a sparse linear combination of archetypes - potentially spanning multiple imaging channels - that are co-regulated across the perturbations. The model was fitted with *max_iter*=500, *tol*=1e-4, using an *L*1 penalty to enforce sparsity in the loading vectors. *K* ∈ {14, 16, 18, 20, 22} were evaluated and *K*=18 was selected as it offered a favorable trade-off between reconstruction fidelity and program interpretability. The per-archetype means were then projected onto the loadings and added back as a scalar shift per program. This resulted in a score matrix reflective of the absolute effect of perturbations on programs, *S^abs^*, rather than relative to the perturbed population.

#### Morphological program annotation

Morphological programs were annotated using an iterative human-AI collaborative workflow. Three inputs were used for each program: (1) per-archetype visual reports generated by the automated image annotation workflow; (2) archetype-program weight matrices from the archetype analysis; and (3) representative cell image grids for each program, to enable direct visual cross-validation. Annotation was performed using an AI agent (Ona, Anthropic Claude 4.6 Opus), operating within a structured analytical framework. A user-defined prompt (supplementary files) instructed the agent to: (i) rank archetypes by absolute weight to establish interpretation priority; (ii) retrieve and cross-reference the automated archetype report for each contributing archetype; (iii) independently examine the program image grids to verify or challenge report-derived descriptions against directly observable morphology; (iv) integrate observations across all contributing archetypes to identify coherent biological themes; and (v) generate a structured program annotation comprising a program composition summary, per-archetype recap with image-grounded descriptions, and an overall biological interpretation. The agent was instructed to prioritize directly observable morphological features over inferred biological function, and to flag discrepancies between archetype reports and image grid observations.

Program annotations were then subjected to iterative user-driven quality control and refinement. The human user reviewed each annotation against the source images and archetype reports, corrected misinterpretations, challenged inferences, and introduced domain-specific interpretive frameworks. These interpretive corrections, once established for one program, were propagated as constraints to subsequent program annotations, creating a cumulative interpretive framework that improved annotation consistency across programs.

#### Projection of programs onto single cells

Programs were projected onto archetype space using SPCA loadings L (n_archetypes_ × K). For a single cell *c*, its score for program *k* was calculated as

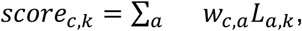

where *w_c_*_,*a*_ is the CLR-transformed weight of archetype *a* for cell *c*.

#### Perturbation clustering from morphological program scores

Perturbations were clustered into co-functional modules in the *K*-dimensional score space *S* using a consensus approach to average over nuisance variation in the clustering procedure (perturbation subsetting and clustering parameters) while keeping the SPCA dimensionality fixed at K=18. 150 resampled runs were performed, each drawing 85% of perturbations without replacement. For each subsample, SPCA was refitted using one of 10 random seeds (varied across runs), and the resulting scores were aligned to a reference loading matrix via the Hungarian algorithm^86^ on the absolute correlation matrix. Each program column was z-transformed prior to clustering so that all programs contributed equally to Euclidean distances. Each resampled score matrix was then clustered under a grid of algorithms and hyperparameters: (i) agglomerative hierarchical clustering with Ward and average linkage, cut at each of {12, 14, 16, …, 28} clusters; and (ii) Leiden community detection on a *k*-nearest neighbor graph with *k* ∈ {5, 8, 10, 12} and resolution ∈ {0.8, 1.0, 1.2, 1.5, 1.8}. For each clustering instance, the pairs of perturbations that were co-assigned to the same cluster were recorded. A consensus co-clustering matrix *C* (n_perturbations_ × n_perturbations_) was computed as C_ij_ = (number of times *i* and *j* were co-clustered) / (number of clustering instances in which both *i* and *j* were present in the subsample). The consensus matrix was converted to a distance matrix *D* = 1 − *C*, and average-linkage hierarchical clustering was applied to *D*. The resulting dendrogram was cut at a specified number of clusters using scipy.cluster.hierarchy.fcluster with criterion=’maxclust’. Cluster robustness was assessed by computing, for each perturbation, the margin between its mean consensus with its assigned cluster and the highest mean consensus with any other cluster; perturbations with margin < 0.15 were flagged as weakly assigned.

For downstream analysis, the score matrix *S* and the clustering results were filtered to include only strong effects. Perturbations were retained if their maximum absolute effect across all archetypes was over 1.2 (∼50% quantile of absolute effects across all perturbations), and archetypes were retained if the maximum absolute effect of any perturbation on them was over 1.5 (threshold determined from the distribution of maximum absolute effects across all archetypes). A perturbation module containing only 2 genes was further removed. This retained 247 out of 471 perturbations and 150 out of 188 archetypes.

Perturbation modules were annotated using a combination of functional enrichment analysis using the Gene Ontology^87^, KEGG^88^ and Reactome^23^ databases, and LLM-assisted summarization.

#### Program contribution to perturbation effect

To attribute each perturbation’s measured effect to individual SPCA programs, we decompose every perturbation vector (signed 1D-Wasserstein distances per archetypes) into the sum of program-aligned components and an unmodelled residual:

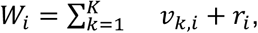

where

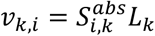

is the partial reconstruction vector of perturbation *i* by program *k*, *S^abs^* is the absolute score of perturbation *i* on program *k* after restoring the mean offset and *L_k_*is the loadings vector for program *k*. We then compute program contributions as the inner product between perturbation effect *W_i_* and the partial contribution vector *v_k_*_,*i*_, normalized by perturbation magnitude ||*W_i_*||:

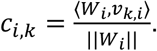

Contributions therefore measure the alignment between the partial reconstruction attributable to program *k* and the perturbation effect, weighted by the magnitude of the program’s contribution to the reconstruction. Perturbation-level contributions were aggregated to the median across perturbations in the same module.

#### Channel usage per program

Archetype loadings *L* (n_archetypes_ × K), resulting from the sparse PCA analysis, were squared and then summed for each channel *c*, resulting in a channel usage (n_channels_ × K) matrix. Channel usages were divided by total usage across the program and the normalized Shannon entropy was computed for each program *k*, defined as

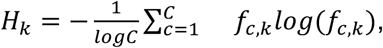

where *f_c_*_,*k*_ is the fraction of summed squared loadings from archetypes defined in channel *c* on program *k* over the total across all channels.

#### Module selectivity for focal programs

To investigate preferential activation of one program over a group of others, programs were restricted to P3, P4, P10 and P16, due to their common patterns of marker usage. For each program, we computed median scores across perturbations within each module and scaled them by the standard deviation of program scores across individual perturbations. To facilitate interpretation, we focused on positive program scores and calculated selectivity of module M for focal program f among programs K as

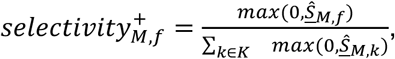

Where 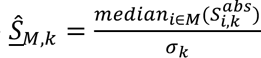 and *σ_k_* is the standard deviation of program k across perturbations.

## Data and Code Availability

All raw imaging data will be deposited into a public repository upon publication. The implementation of CHART along with tutorials is publicly available on GitHub (https://github.com/Genentech/CHART).

## Acknowledgements

We thank Leslie Gaffney for her help in figure making. We also thank Jose Sergio Hleap, Joshua Gould and Bo Li for their guidance in running scallops, and Jon Scherdin, Lindsay Liang, Jay Cowan and the BigDipIR team for their help deploying scallops at scale.

## Contributions

P.P.C. and A.C.L led and conceived the project. P.P.C., T.K., A.S. developed the experimental design, P.P.C., performed the experiments. P.W. generated and validated the cell line. A.C.L. developed and implemented the CHART framework, with input from P.P.C. A.C.L., H.Y. performed embedding evaluations. P.P.C. introduced LLM workflow with support of A.C.L and H.W. V.E., A. S., T. M., R.L., D.R. advised on and supported analysis. T.H. and L.G. advised on the interpretation of screens. P.P.C, A.C.L., A.S., J.M. and A.R. wrote the manuscript. A.S., J.M. and A.R. supervised the work.

## EXTENDED DATA FIGURE LEGENDS

**Extended Data Figure 1.**
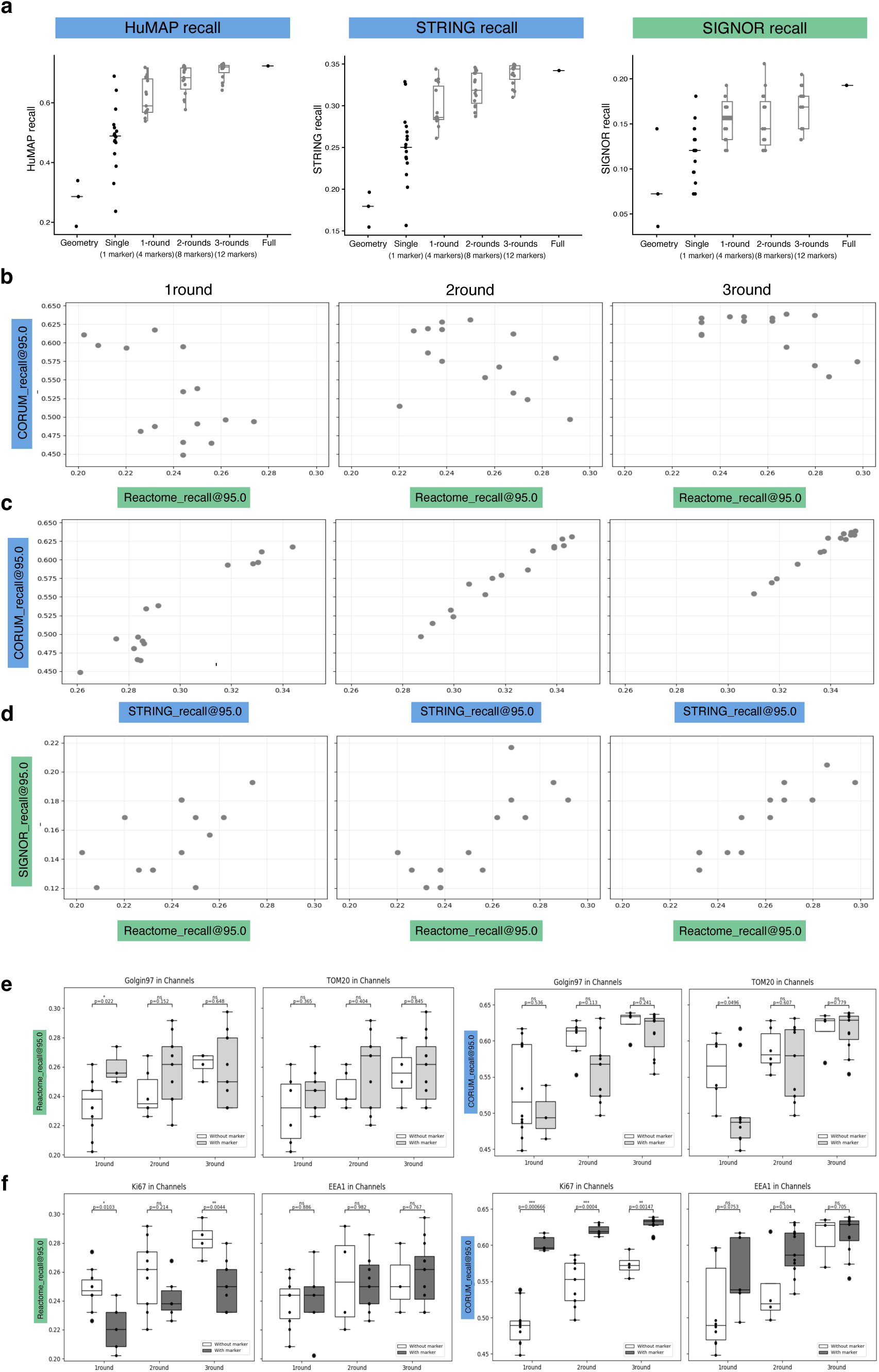
Database recall behaviour across sub-sampled panel embeddings. **a.** Performance of sub-sampled panel embeddings in recovering known biological relationships in different databases. Recall from each database (y axis, defined as in Fig. 2e) for 15 random panels were selected for the 1, 2 or 3 round categories (x axis). **b–d**. Pairwise relationship between database recall across random panels. Each dot is one of the 15 random panels selected for the 1-, 2- or 3-round category; axes show recall@95.0 for a pair of databases. **b,** CORUM (y) versus Reactome (x) recall, comparing a protein-complex database to a pathway database. **c,** CORUM (y) versus STRING (x) recall, comparing two interaction/complex-based databases. **d,** SIGNOR (y) versus Reactome (x) recall, comparing two signaling/pathway databases. Complex- and pathway-based recovery are weakly correlated. **e–f**. Effect of individual marker inclusion on complex recovery. CORUM recall@95.0 (y axis) for the 15 random panels of each round (x axis, 1/2/3 round), split by whether a panel contains (with marker, gray) or lacks (without marker, white) the indicated marker(s) (columns); dots, individual panels. e, Golgin97, TOM20, and TOM20 or Golgin97. f, Ki67, EEA1, and EEA1 or Ki67. Nominal two-sided Welch’s t-tests were used to compare panels with vs without the indicated marker(s) within each round.

**Extended Data Figure 2.**
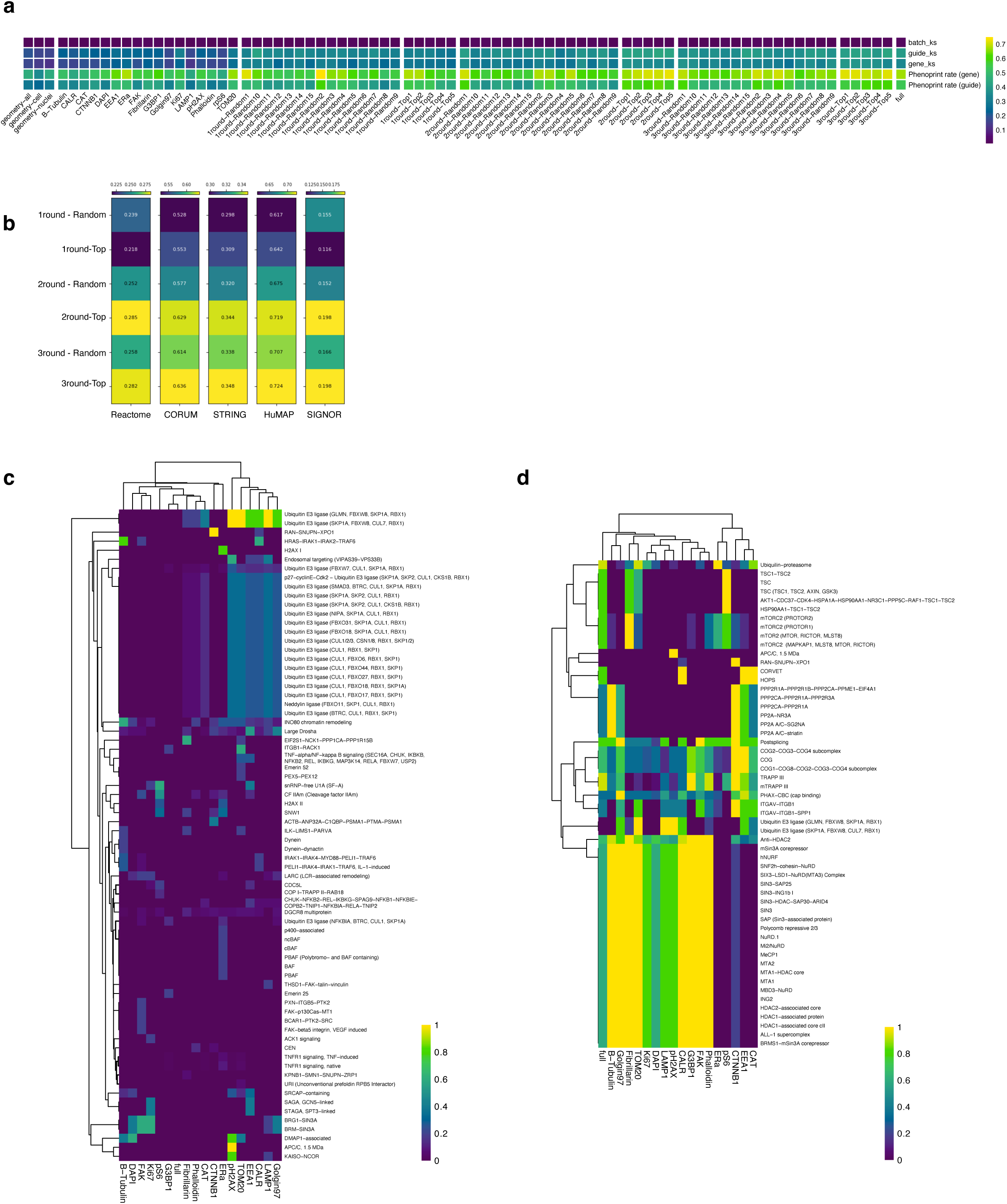
Subsampled panel performance and marker-specific recovery of biological relationships. Dashed horizontal line: corresponds to random recovery of 0.1. **a.** Consistency metrics for sub-panels. Consistency (color bar) for different subspanels (columns) including by geometry, representative top performing and randomly selected sub-sets across panel complexity, and the full panel, by different criteria (rows). *batch_ks, guide_ks, gene_ks*: Kolmogorov-Smirnov (KS) statistics between pairs of profiles with the same batch (well), guide or gene ID and randomly selected pairs. Low *batch_ks* values: successful batch effect removal. *Phenoprint rate (gene), Phenoprint rate (guide):* fraction of genes with distinct profiles from control as measured by energy-distance (at p-val < 0.05). **b.** Comparison of randomly sub-sampled embeddings to those with highest local similarity to full-panel embedding. Mean recall from each database (columns, defined as in Fig. 2e) for 15 random panels or 5 top performing panels (highest local similarity to full embedding) for the 1, 2 or 3 round categories (rows). **c,d.** Recovery of CORUM complexes by individual markers or all 17 markers. Recall at 95% (two sides, as in Fig. 2e, color bar) for CORUM complexes (rows) based on individual markers or the full panel (columns) for complexes were filtered by those captured in any single-marker panel but not by the full panel (recall=0, c), reflecting cases of complete loss of biological relationships when incorporating additional markers, or for complexes captured fully by at least one single-marker panel (recall=1) but not captured fully by the full panel, reflecting cases of partial loss of biological relationships when incorporating additional markers (**d**).

**Extended Data Figure 3.**
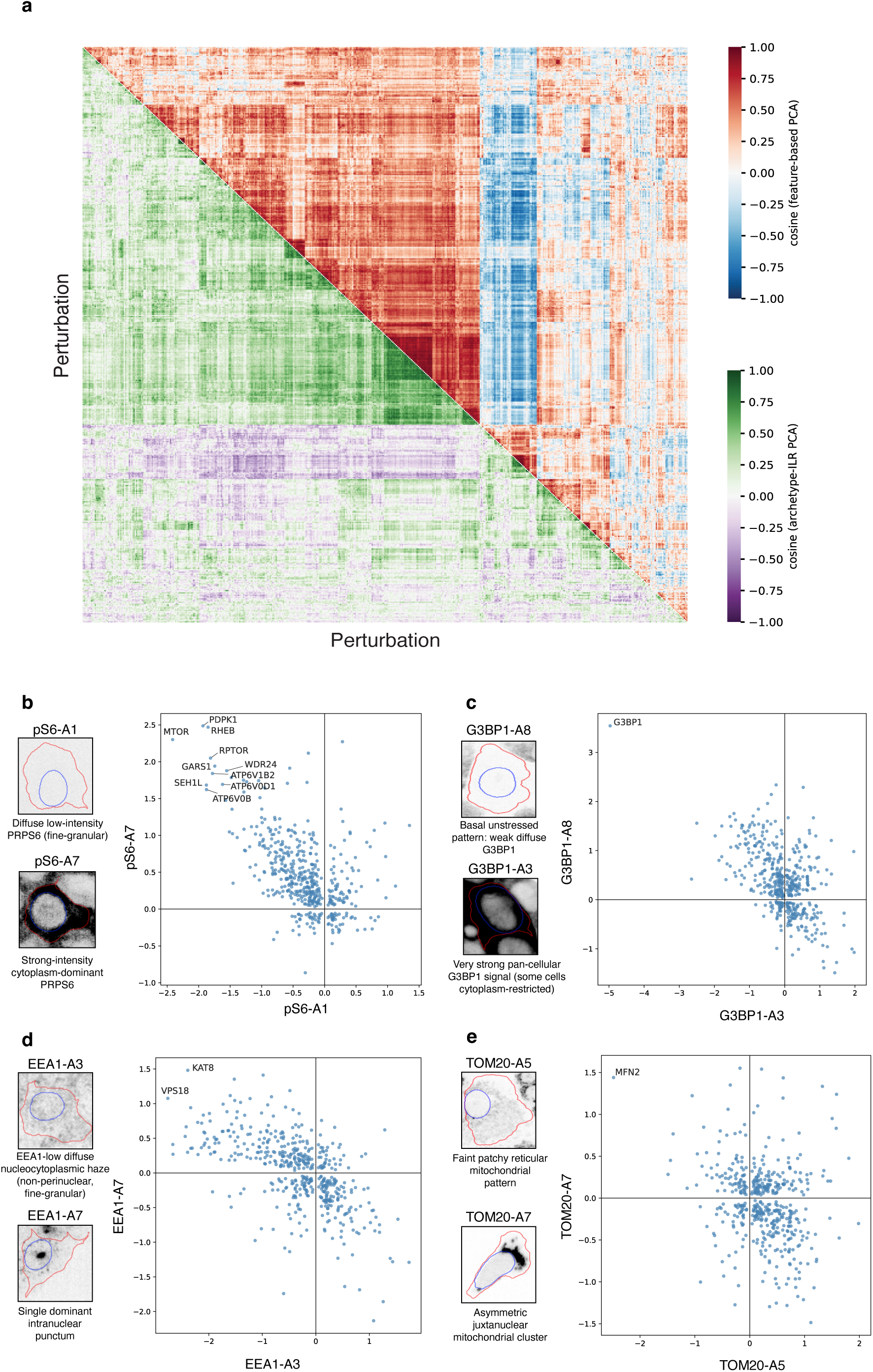
Archetype profiles capture perturbation relationships and expected marker phenotypes. **a.** Comparison of perturbation relationships captured by engineered features and archetypes. Pairwise cosine similarity (color bars) between perturbations profiles defined across engineered features (blue-red scale) or ILR-transformed archetype weights (purple-green scale), ordered by hierarchical clustering performed on the average between the two z-scored matrices (**Methods**). **b-e.** Example archetypes from selected markers. Effects (x and y axes) of each perturbation (dot) on each archetype. Left: Representative images of each archetype. Labeled genes: perturbations with expected effect. Crosshairs: zero effect relative to control.

**Extended Data Figure 4.**
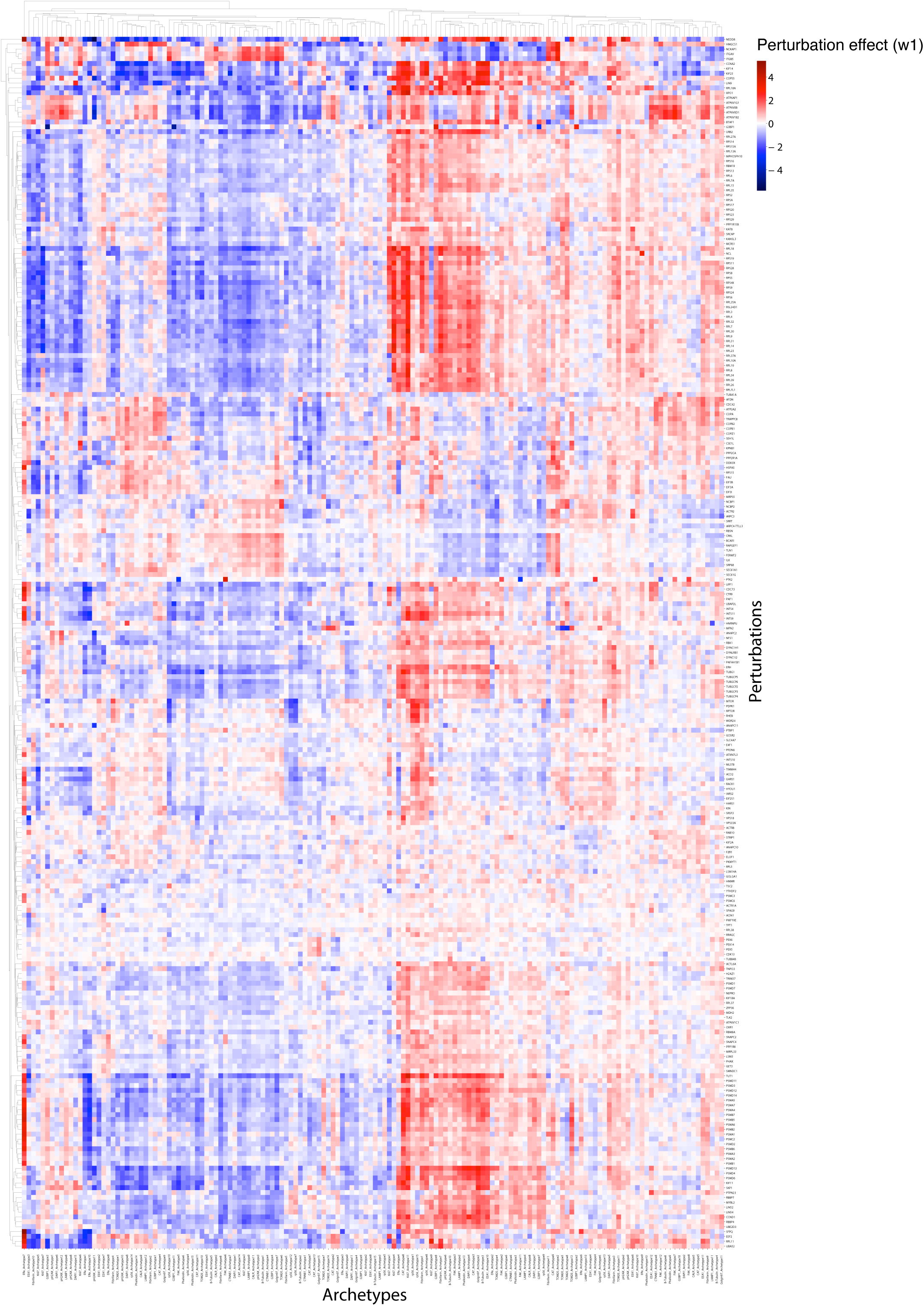
Modular perturbation effects on archetypes. Signed effects (color bar) of perturbations (rows) on archetypes (columns) represent shifts towards (positive values) or away from (negative values) the archetypal phenotype compared to the control population of cells.

**Extended Data Figure 5.**
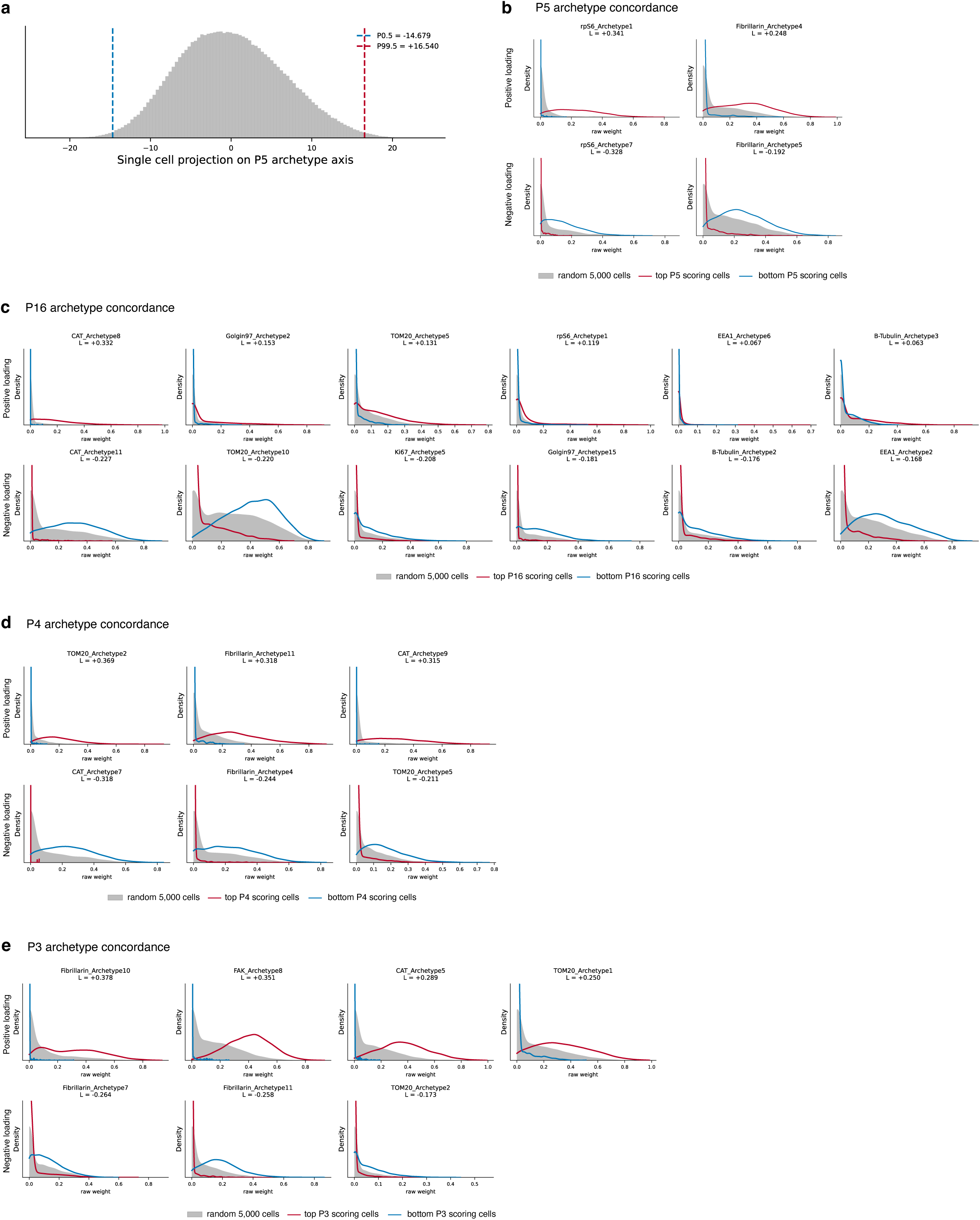
Single-cell-level concordance of archetype shifts across top-contributing members of the same program. **a.** Distribution of single cell projections along program axes (x-axis) for P5, obtained through archetype loadings on programs (**Methods**). Top positive and negative scoring cells are chosen above the 99.5 (red vertical dashed line) and below the 0.05 (blue vertical dashed line) percentiles. **b.-e.** Distribution of archetype weights (x-axis) for top positive projections (red line), top negative projections (blue line) and 5,000 randomly sampled cells (grey area). Archetype weights are shown for selected top contributing archetypes in programs P5 (**b.**), P16 (**c.**), P14 (**d.**) and P3 (**e.**), with either strongest positive (top rows) or strongest negative (bottom rows) loadings.

**Extended Data Figure 6.**
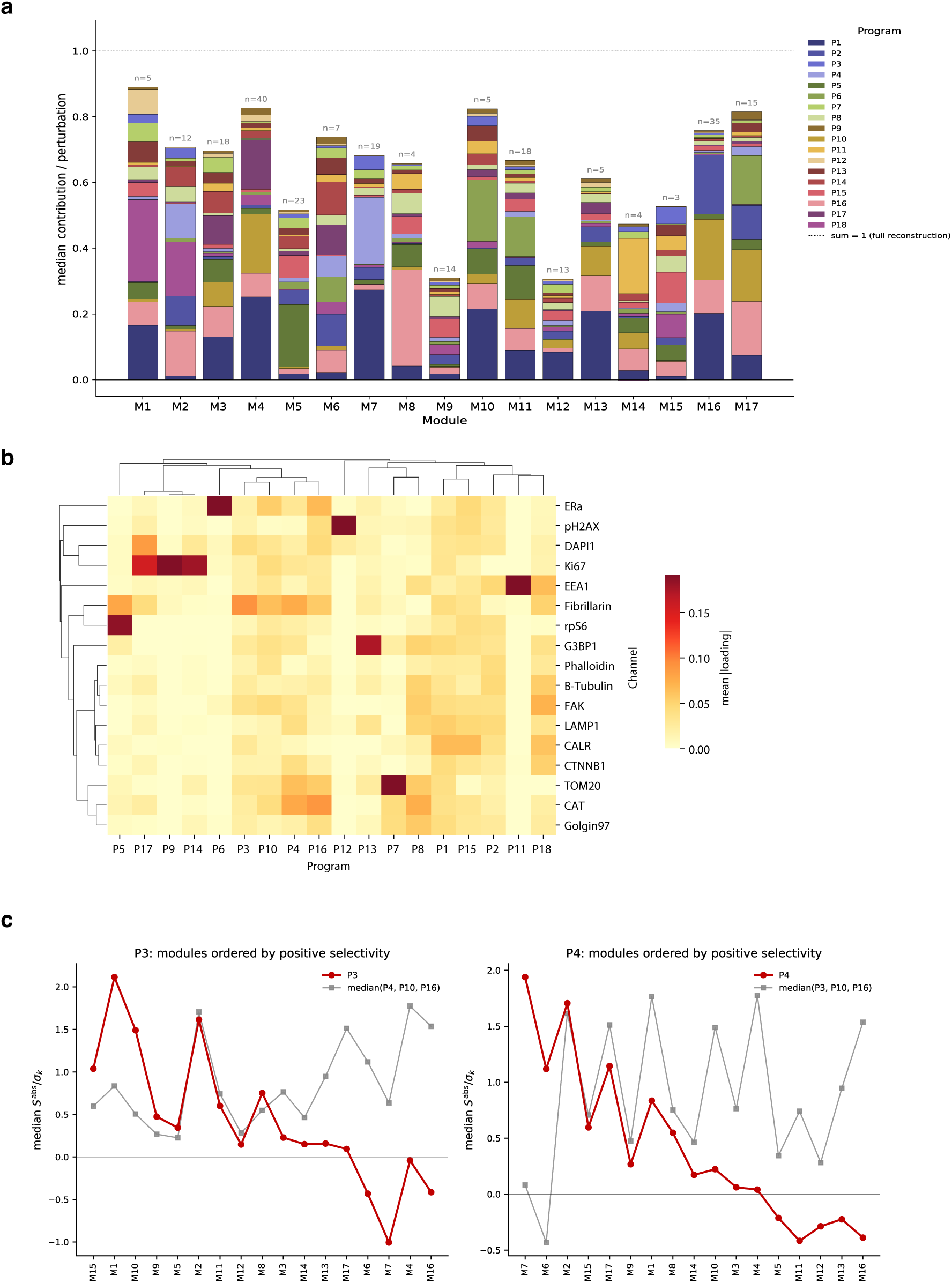
Program contributions, marker usage and perturbation selectivity across morphological programs. **a.** Program contribution to the reconstruction of perturbation module effects. Contributions (y-axis) reflect the share of perturbation effect that is attributable to each program (**Methods**) and are here aggregated to the median across perturbations within each module. Stacked contributions reflect the total recovered perturbation effect across programs (perfect recovery shown by horizontal line at 1). **b.** Aggregate marker loadings per program. Mean absolute loadings (color bar) across archetypes of each marker (row) on each program (column), showing patterns of marker usage across programs (hierarchical clustering on rows and columns). **c.** Perturbation selectivity within a program set with similar marker usage patterns (P3, P4, P10 and P16). Program score aggregated to the median across perturbations in the same module and scaled by program score standard deviation across perturbations (y-axis), across perturbation modules (x-axis). Scores are shown separately for focal programs (red dots) P3 (right) and P4 (left), against the median across the remaining 3 programs (grey dots). Modules (x-axis) are ordered by decreasing selectivity for the highlighted program, computed as the fraction of median scaled scores for the focal program among the median scaled scores across the program set.

## REFERENCES

1. Lawson, M. J. et al. In situ genotyping of a pooled strain library after characterizing complex phenotypes. Mol Syst Biol 13, MSB177951 (2017).

2. Emanuel, G., Moffitt, J. R. & Zhuang, X. High-throughput, image-based screening of pooled genetic-variant libraries. Nat Methods 14, 1159–1162 (2017).

3. Feldman, D. et al. Optical Pooled Screens in Human Cells. Cell 179, 787–799.e17 (2019).

4. Walton, R. T., Singh, A. & Blainey, P. C. Pooled genetic screens with image-based profiling. Molecular Systems Biology 18, e10768 (2022).

5. Carlson, R. J., Leiken, M. D., Guna, A., Hacohen, N. & Blainey, P. C. A genome-wide optical pooled screen reveals regulators of cellular antiviral responses. Proceedings of the National Academy of Sciences 120, e2210623120 (2023).

6. Funk, L. et al. The phenotypic landscape of essential human genes. Cell 10.1016/j.cell.2022.10.017 (2022) doi:10.1016/j.cell.2022.10.017.

7. Le, A., Biederer, T. & Blainey, P. C. High-content image-based pooled screens reveal regulators of synaptogenesis. Cell Reports 44, 115889 (2025).

8. Camsund, D. et al. Time-resolved imaging-based CRISPRi screening. Nat Methods 17, 86–92 (2020).

9. A genome-wide atlas of human cell morphology | Nature Methods. https://www.nature.com/articles/s41592-024-02537-7.

10. Seal, S. et al. Cell Painting: a decade of discovery and innovation in cellular imaging. Nat Methods 22, 254–268 (2025).

11. Seal, S., et al. From pixels to phenotypes: Integrating image-based profiling with cell health data as BioMorph features improves interpretability. MBoC 35, mr2 (2024).

12. Stirling, D. R. et al. CellProfiler 4: improvements in speed, utility and usability. BMC Bioinformatics 22, 433 (2021).

13. Sivanandan, S. et al. A pooled Cell Painting CRISPR screening platform enables de novo inference of gene function by self-supervised deep learning. Nature Communications 17, 77 (2025).

14. Kraus, O. et al. Masked Autoencoders for Microscopy are Scalable Learners of Cellular Biology. in 2024 IEEE/CVF Conference on Computer Vision and Pattern Recognition (CVPR) 11757–11768 (2024). doi:10.1109/CVPR52733.2024.01117.

15. Yao, H., Hanslovsky, P., Huetter, J.-C., Hoeckendorf, B. & Richmond, D. Weakly Supervised Set-Consistency Learning Improves Morphological Profiling of Single-Cell Images. Preprint at 10.48550/arXiv.2406.05308 (2024).

16. Wang, Z. J., et al. Multi-ContrastiveVAE disentangles perturbation effects in single cell images from optical pooled screens. bioRxiv 2023.11.28.569094 (2024) doi:10.1101/2023.11.28.569094.

17. Makino, T., et al. Supervised Contrastive Block Disentanglement. Preprint at 10.48550/arXiv.2502.07281 (2025).

18. Bray, M.-A. et al. Cell Painting, a high-content image-based assay for morphological profiling using multiplexed fluorescent dyes. Nat Protoc 11, 1757–1774 (2016).

19. von Coburg, E. et al. Cell Painting PLUS: expanding the multiplexing capacity of Cell Painting-based phenotypic profiling using iterative staining-elution cycles. Nat Commun 16, 3857 (2025).

20. Radtke, A. J. et al. IBEX: A versatile multiplex optical imaging approach for deep phenotyping and spatial analysis of cells in complex tissues. Proceedings of the National Academy of Sciences 117, 33455–33465 (2020).

21. Kudo, T. et al. Highly multiplexed, image-based pooled screens in primary cells and tissues with PerturbView. 2023.12.26.573143 Preprint at 10.1101/2023.12.26.573143 (2023).

22. Gould, J. et al. SCALLOPS: a scalable, integrated computational framework for Optical Pooled Screens. 2026.05.22.727250 Preprint at 10.64898/2026.05.22.727250 (2026).

23. Reactome Pathway Knowledgebase 2024 | Nucleic Acids Research | Oxford Academic. https://academic.oup.com/nar/article/52/D1/D672/7369850?login=false.

24. Lo Surdo, P., et al. SIGNOR 4.0: the 2025 update with focus on phosphorylation data. Nucleic Acids Res gkaf1237 (2025) doi:10.1093/nar/gkaf1237.

25. Szklarczyk, D. et al. The STRING database in 2025: protein networks with directionality of regulation. Nucleic Acids Res 53, D730–D737 (2025).

26. Steinkamp, R. et al. CORUM in 2024: protein complexes as drug targets. Nucleic Acids Res 53, D651–D657 (2025).

27. Fischer, S. N. et al. hu.MAP3.0: atlas of human protein complexes by integration of >25,000 proteomic experiments. Mol Syst Biol 21, 911–943 (2025).

28. Cutler, A. & Breiman, L. Archetypal Analysis. Technometrics 36, 338–347 (1994).

29. Seth, S. & Eugster, M. J. A. Probabilistic archetypal analysis. Mach Learn 102, 85– 113 (2016).

30. Eugster, M. J. A. & Leisch, F. Weighted and robust archetypal analysis. Computational Statistics & Data Analysis 55, 1215–1225 (2011).

31. Eugster, M. J. A. & Leisch, F. From Spider-Man to Hero — Archetypal Analysis in R. Journal of Statistical Software 30, 1–23 (2009).

32. Baran, Y. et al. MetaCell: analysis of single-cell RNA-seq data using K-nn graph partitions. Genome Biology 20, 206 (2019).

33. Bilous, M. et al. Metacells untangle large and complex single-cell transcriptome networks. BMC Bioinformatics 23, 336 (2022).

34. Ma, C., Trinh, L. T., Bucci, M., Regev, A. & Wang, H. Orion: Towards Lab Automation with Computer-Using Agents. 2026.06.13.732095 Preprint at 10.64898/2026.06.13.732095 (2026).

35. Magnuson, B., Ekim, B. & Fingar, D. C. Regulation and function of ribosomal protein S6 kinase (S6K) within mTOR signalling networks. Biochem J 441, 1–21 (2011).

36. van, der B. J., et al. Loss of the HOPS complex disrupts early-to-late endosome transition, impairs endosomal recycling and induces accumulation of amphisomes. Molecular Biology of the Cell 35, ar40 (2024).

37. Bach, D. et al. Mitofusin-2 Determines Mitochondrial Network Architecture and Mitochondrial Metabolism: A NOVEL REGULATORY MECHANISM ALTERED IN OBESITY*. Journal of Biological Chemistry 278, 17190–17197 (2003).

38. Kwon, M., Leibowitz, M. L. & Lee, J.-H. Small but mighty: the causes and consequences of micronucleus rupture. Exp Mol Med 52, 1777–1786 (2020).

39. Duan, H. et al. Micronuclei: origins, assays, mechanisms, diseases and treatments. Sig Transduct Target Ther 11, 114 (2026).

40. Cross, R. A. & McAinsh, A. Prime movers: the mechanochemistry of mitotic kinesins. Nat Rev Mol Cell Biol 15, 257–271 (2014).

41. Derenzini, M. et al. Nucleolar function and size in cancer cells. Am J Pathol 152, 1291–1297 (1998).

42. Derenzini, M. et al. Nucleolar size indicates the rapidity of cell proliferation in cancer tissues. The Journal of Pathology 191, 181–186 (2000).

43. Buchwalter, A. & Hetzer, M. W. Nucleolar expansion and elevated protein translation in premature aging. Nat Commun 8, 328 (2017).

44. Balderhaar, H. J. kleine & Ungermann, C. CORVET and HOPS tethering complexes – coordinators of endosome and lysosome fusion. J Cell Sci 126, 1307–1316 (2013).

45. Olesen, C. W. et al. Trafficking, localization and degradation of the Na+,HCO3− co-transporter NBCn1 in kidney and breast epithelial cells. Sci Rep 8, 7435 (2018).

46. Severin, M. et al. Dynamic localization of the Na+-HCO3− co-transporter NBCn1 to the plasma membrane, centrosomes, spindle and primary cilia. J Cell Sci 136, jcs260687 (2023).

47. Abshire, E. T. & Maquat, L. E. Gene regulation through exon junction complex modularity. Nat Struct Mol Biol 32, 2387–2397 (2025).

48. Haberman, A. S., Akbar, M. A., Ray, S. & Krämer, H. Drosophila acinus encodes a novel regulator of endocytic and autophagic trafficking. Development 137, 2157–2166 (2010).

49. Vasanthakumar, T. & Rubinstein, J. L. Structure and Roles of V-type ATPases. Trends in Biochemical Sciences 45, 295–307 (2020).

50. Barua, S., Berger, S., Pereira, E. M. & Jobanputra, V. Expanding the phenotype of ATP6AP1 deficiency. Cold Spring Harb Mol Case Stud 8, a006195 (2022).

51. Casey, J. R., Grinstein, S. & Orlowski, J. Sensors and regulators of intracellular pH. Nat Rev Mol Cell Biol 11, 50–61 (2010).

52. Marshansky, V. & Futai, M. The V-type H+-ATPase in vesicular trafficking: targeting, regulation and function. Current Opinion in Cell Biology 20, 415–426 (2008).

53. Praschberger, R. et al. Mutations in Membrin/*GOSR2* Reveal Stringent Secretory Pathway Demands of Dendritic Growth and Synaptic Integrity. Cell Reports 21, 97–109 (2017).

54. Liu, Y. et al. Myosin Vb controls biogenesis of post-Golgi Rab10 carriers during axon development. Nat Commun 4, 2005 (2013).

55. Liu, S. et al. RAB-10 cooperates with EHBP-1 to capture vesicular carriers during post-Golgi exocytic trafficking. J Cell Biol 224, e202410003 (2025).

56. Nik Nabil, W. N., et al. Towards a Framework for Better Understanding of Quiescent Cancer Cells. Cells 10, 562 (2021).

57. Melk, A. et al. Expression of p16INK4a and other cell cycle regulator and senescence associated genes in aging human kidney. Kidney International 65, 510–520 (2004).

58. Braun, S. et al. Nuclear Dynamics in Quiescent Cells: Conserved Mechanisms from Yeasts to Mammals. Biomolecules 16, 203 (2026).

59. Pathak, R. U., Soujanya, M. & Mishra, R. K. Deterioration of nuclear morphology and architecture: A hallmark of senescence and aging. Ageing Research Reviews 67, 101264 (2021).

60. Tiku, V. & Antebi, A. Nucleolar Function in Lifespan Regulation. Trends in Cell Biology 28, 662–672 (2018).

61. Caicedo, J. C. et al. Data-analysis strategies for image-based cell profiling. Nature Methods 14, 849–863 (2017).

62. Qiu, M. et al. A cell-level quality control workflow for high-throughput image analysis. BMC Bioinformatics 21, 280 (2020).

63. Arevalo, J. et al. Evaluating batch correction methods for image-based cell profiling. Nature Communications 15, 6516 (2024).

64. Brückner, L., et al. Extrachromosomal DNA micronucleation constrains tumour fitness and improves patient survival. bioRxiv 2025.04.15.648906 (2026) doi:10.1101/2025.04.15.648906.

65. Houlahan, K. E. et al. Complex rearrangements fuel ER+ and HER2+ breast tumours. Nature 638, 510–518 (2025).

66. Okumoto, K., Tamura, S., Honsho, M. & Fujiki, Y. Peroxisome: Metabolic Functions and Biogenesis. in Peroxisome Biology: Experimental Models, Peroxisomal Disorders and Neurological Diseases (ed. Lizard, G.) 3–17 (Springer International Publishing, Cham, 2020). doi:10.1007/978-3-030-60204-8_1.

67. Shukla, N. et al. TOR and heat shock response pathways regulate peroxisome biogenesis during proteotoxic stress. Nat Commun 16, 10743 (2025).

68. Bajdzienko, J. & Bremm, A. Mammalian pexophagy at a glance. J Cell Sci 137, jcs259775 (2024).

69. Han, J., Zheng, D., Liu, P.-S., Wang, S. & Xie, X. Peroxisomal homeostasis in metabolic diseases and its implication in ferroptosis. Cell Commun Signal 22, 475 (2024).

70. Da Graça, J., Delevoye, C. & Morel, E. Morphodynamical adaptation of the endolysosomal system to stress. FEBS J 292, 248–260 (2025).

71. López-Hernández, T., Haucke, V. & Maritzen, T. Endocytosis in the adaptation to cellular stress. Cell Stress 4, 230–247.

72. Morrison, L., Loibl, S. & Turner, N. C. The CDK4/6 inhibitor revolution — a game-changing era for breast cancer treatment. Nat Rev Clin Oncol 21, 89–105 (2024).

73. Gandhi, V., Plunkett, W. & Cortes, J. E. Omacetaxine: A Protein Translation Inhibitor for Treatment of Chronic Myelogenous Leukemia. Clin Cancer Res 20, 1735–1740 (2014).

74. Rosen, E. et al. 367P Phase I/II dose escalation study evaluating first-in-class eIF4A inhibitor zotatifin in ER+ metastatic breast cancer. Annals of Oncology 35, S372 (2024).

75. Gao, T. et al. CDK4/6 inhibitors in breast cancer therapy: mechanisms of drug resistance and strategies for treatment. Front. Pharmacol. 16, (2025).

76. Feldman, D. et al. Pooled genetic perturbation screens with image-based phenotypes. Nat Protoc 17, 476–512 (2022).

77. Hoberecht, L., Perampalam, P., Lun, A. & Fortin, J.-P. A comprehensive Bioconductor ecosystem for the design of CRISPR guide RNAs across nucleases and technologies. Nat Commun 13, 6568 (2022).

78. Singh, S., Bray, M.-A., Jones, T. & Carpenter, A. Pipeline for illumination correction of images for high-throughput microscopy. J Microsc 256, 231–236 (2014).

79. Muhlich, J. L. et al. Stitching and registering highly multiplexed whole-slide images of tissues and tumors using ASHLAR. Bioinformatics 38, 4613–4621 (2022).

80. F. T. Liu, K. M. Ting, & Z. -H. Zhou. Isolation Forest. in 2008 Eighth IEEE International Conference on Data Mining 413–422 (2008). doi:10.1109/ICDM.2008.17.

81. Liu, F. T., Ting, K. M. & Zhou, Z.-H. Isolation-Based Anomaly Detection. ACM Trans. Knowl. Discov. Data 6, (2012).

82. Breunig, M. M., Kriegel, H.-P., Ng, R. T. & Sander, J. LOF: Identifying Density-Based Local Outliers.

83. Rousseeuw, P. & Driessen, K. A Fast Algorithm for the Minimum Covariance Determinant Estimator. Technometrics 41, 212–223 (1999).

84. Ross, D. A., Lim, J., Lin, R.-S. & Yang, M.-H. Incremental Learning for Robust Visual Tracking. Int J Comput Vis 77, 125–141 (2008).

85. Celik, S. et al. Building, benchmarking, and exploring perturbative maps of transcriptional and morphological data. PLOS Computational Biology 20, e1012463 (2024).

86. Kuhn, H. W. The Hungarian method for the assignment problem. Naval Research Logistics Quarterly 2, 83–97 (1955).

87. Ashburner, M. et al. Gene Ontology: tool for the unification of biology. Nature Genetics 25, 25–29 (2000).

88. Kanehisa, M. Toward understanding the origin and evolution of cellular organisms. Protein Science 28, 1947–1951 (2019).

