## Supplementary Figures and Legends for "Archetype analysis of multiplexed antibody imaging reveals interpretable morphological programs in optical pooled screens"

#### SUPPLEMENTARY FIGURE LEGENDS

**Supplementary Figure 1. CellMAPP panel design, and MCF-7-iCas9 clone validation**

**a, b.** Comparison of the MCF-7-iCas9 high-efficiency knockout clone A08 with the parental MCF-7 cell line. **a.** Immunofluorescence staining of estrogen receptor α (ERα-594) and the ER target GREB1-647 confirms baseline signal and comparable signal reduction after ER inhibition (Ggiredestrant) in clonal MCF-7-iCas9-A08, bulk MCF-7-iCas9 cells and the parental MCF-7 line**. b.** Giredestrant dose–response viability assay showing comparable sensitivity between lines**.**

**c.** Panel rounds. Representative images from multiplexed immunofluorescence staining of markers across imaging rounds 1–4, each covering DAPI and four different antibodies.

**d.** Validation of Consistency in localization and staining quality across iterative imaging cycles with signal removal using IBEX[20] (Iterative Bleaching EXtends multiplexing). Representative single-channel images after staining in each round (top) and in corresponding IBEX controls (bottom).

**Supplementary Figure 2. Pre-processing pipeline**

**a.** Feature preprocessing pipeline overview. From left: Cells were filtered based on nuclear size, registration quality, and label transfer accuracy. Low-variability and NA-rich features were removed, followed by quantile normalization and incremental PCA for embedding computation. Outlier cells and guides were excluded prior to re-embedding, yielding normalized single-cell and gene-level feature tables (**Methods**).

**b.** Effective batch correction. UMAP of the embedding of single-cell profiles (dots) based on morphological features from all 17 markers, colored by batch (well).

**Supplementary Figure 3. Automated archetype annotation**

**a.** MLLM-based archetype annotation workflow. From left: Representative images from high-scoring cells (top left) were used to generate initial archetype descriptions (1), which were refined by contrastive comparison (2) with a user-defined baseline archetype from the same marker (bottom left) . Refined annotations were combined into marker-level reports (3), evaluated for semantic similarity and, when necessary, re-analyzed side-by-side using the original images (4) to generate final archetype reports (right).

**Supplementary Figure 4. Examples of annotated morphological archetypes recovered**
Representative archetype image selected from among the top five cells with the highest archetype weight (left) and MLLM-defined archetype names (right) for three archetypes capturing distinct morphological states across eight markers.

**Supplementary Figure 5. Representative archetype reports**

Representative examples from select archetype reports for Fibrilarin and Tubulin. Each example shows the archetype name (top), images from the five cells with the highest archetype weights used for annotation (middle), and excerpt from the final archetype summary (bottom).

**Supplementary Figure 6. Single-cell co-occurrence of archetypes within morphological programs**

**a–d.** Single-cell weights for archetypes with non-zero loadings in morphological programs P3 (**a.**), P4 (**b.**), P5 (**c.**) and P16 (**d.**). Archetype loadings for each program are shown above (color bars). Cells with the highest positive and negative projections along each program axis were selected above the 99.5th percentile and below the 0.05th percentile, respectively. Archetype-weight distributions are shown for high-scoring cells (red half-violin plots; upper half of heatmaps), low-scoring cells (blue half-violin plots; lower half of heatmaps) and all cells (grey half-violin plots). The vertical white stripe observed for rpS6-A1 in P5 reflect the competition with rpS6-A15, the highest loading archetype for this program; the stripe observed for Fibrillarin-A3 in P16 reflects archetype weights that are close to zero across nearly all cells, as also indicated by the corresponding grey reference distributions.

**Supplementary Figure 7. Detailed annotation for P7**

Morphological program 7 (P7) archetype loadings are shown at the top. Representative images for positively and negatively weighted archetypes are selected from among the five cells with the highest archetype weights and grouped by marker. AI-generated textual summaries for each marker or marker group are shown below the corresponding image panels. The AI-generated biological interpretation of P7 is shown at the bottom.

**Supplementary Figure 8. Detailed annotation for P11**

Morphological program 11 (P11) archetype loadings are shown at the top. Representative images for positively and negatively weighted archetypes are selected from among the five cells with the highest archetype weights and grouped by marker. AI-generated textual summaries for each marker or marker group are shown below the corresponding image panels. The AI-generated biological interpretation of P11 is shown at the bottom.

**Supplementary Figure 9. Detailed annotation for P5**

Morphological program 5 (P5) archetype loadings are shown at the top. Representative images for positively and negatively weighted archetypes are selected from among the five cells with the highest archetype weights and grouped by marker. AI-generated textual summaries for each marker or marker group are shown below the corresponding image panels. The AI-generated biological interpretation of P5 is shown at the bottom.

**Supplementary Figure 10. Detailed annotation for P16**

Morphological program 16 (P16) archetype loadings are shown at the top. Representative images for positively and negatively weighted archetypes are selected from among the five cells with the highest archetype weights and grouped by marker. AI-generated textual summaries for each marker or marker group are shown below the corresponding image panels. The AI-generated biological interpretation of P16 is shown at the bottom.

**Supplementary Figure 11. Detailed annotation for P10**

Morphological program 10 (P10) archetype loadings are shown at the top. Representative images for positively and negatively weighted archetypes are selected from among the five cells with the highest archetype weights and grouped by marker. AI-generated textual summaries for each marker or marker group are shown below the corresponding image panels. The AI-generated biological interpretation of P10 is shown at the bottom.

**Supplementary Figure 12. Detailed annotation for P4**

Morphological program 4 (P4) archetype loadings are shown at the top. Representative images for positively and negatively weighted archetypes are selected from among the five cells with the highest archetype weights and grouped by marker. AI-generated textual summaries for each marker or marker group are shown below the corresponding image panels. The AI-generated biological interpretation of P4 is shown at the bottom.

**Supplementary Figure 13. Detailed annotation for P3**

Morphological program 3 (P3) archetype loadings are shown at the top. Representative images for positively and negatively weighted archetypes are selected from among the five cells with the highest archetype weights and grouped by marker. AI-generated textual summaries for each marker or marker group are shown below the corresponding image panels. The AI-generated biological interpretation of P3 is shown at the bottom.

**Supplementary Figure 14. Single cell representative images of P3 and P4 morphological programs.**

**a,b.** Representative single-cell images for cells with high positive projections onto P3 (a) or P4 (b), selected from the top-scoring cells based on the single-cell program projections quantified in Extended Data Fig. 5. For each program, three representative cells are shown across seven channels with positive archetype loadings, colored as indicated. The final column shows the merged overlay, illustrating co-occurring marker-level morphological changes within the same individual cells.

**Supplementary Figure 15. Detailed annotation for P18**

Morphological program 18 (P18) archetype loadings are shown at the top. Representative images for positively and negatively weighted archetypes are selected from among the five cells with the highest archetype weights and grouped by marker. AI-generated textual summaries for each marker or marker group are shown below the corresponding image panels. The AI-generated biological interpretation of P18 is shown at the bottom.

**Supplementary Figure 16. Detailed annotation for P6**

Morphological program 6 (P6) archetype loadings are shown at the top. Representative images for positively and negatively weighted archetypes are selected from among the five cells with the highest archetype weights and grouped by marker. AI-generated textual summaries for each marker or marker group are shown below the corresponding image panels. The AI-generated biological interpretation of P6 is shown at the bottom.

**Supplementary Figure 17. Detailed annotation for P17**

Morphological program 17 (P17) archetype loadings are shown at the top. Representative images for positively and negatively weighted archetypes are selected from among the five cells with the highest archetype weights and grouped by marker. AI-generated textual summaries for each marker or marker group are shown below the corresponding image panels. The AI-generated biological interpretation of P17 is shown at the bottom.

**Supplementary Figure 18. Detailed annotation for P2**

Morphological program 2 (P2) archetype loadings are shown at the top. Representative images for positively and negatively weighted archetypes are selected from among the five cells with the highest archetype weights and grouped by marker. AI-generated textual summaries for each marker or marker group are shown below the corresponding image panels. The AI-generated biological interpretation of P2 is shown at the bottom.

**Supplementary Figure 19. Cosine-based local similarity of subset panels to full**

Distribution of local similarity (cosine-based) to the full panel embedding (y axis, partial AURC@30 of full embedding neighbor recall) for embeddings generated on each class of panel subset (x axis).

**Supplementary Figure 20: Archetype abundance quantification**

**a.** Total archetype weight (sum over cells of untransformed archetype weights [0,1]).

**b.** Number of cells in which a given archetype is dominant. Dominant archetypes (highest weights) were identified separately for each marker.

**Supplementary Figure 21. Examples of corrected MLLM archetype annotations**
Representative examples of archetypes misannotated by the MLLM workflow. For each archetype, a grid containing the five cells with the highest archetype weights used for annotation is shown at the top, with the original MLLM-generated label and the corrected archetype label shown below. Misannotations most commonly occurred for archetypes with extreme or largely homogeneous staining patterns, elevated background, apparent nuclear–cytosolic spillover, or spillover from neighboring cells.

### Supplementary Figure 1

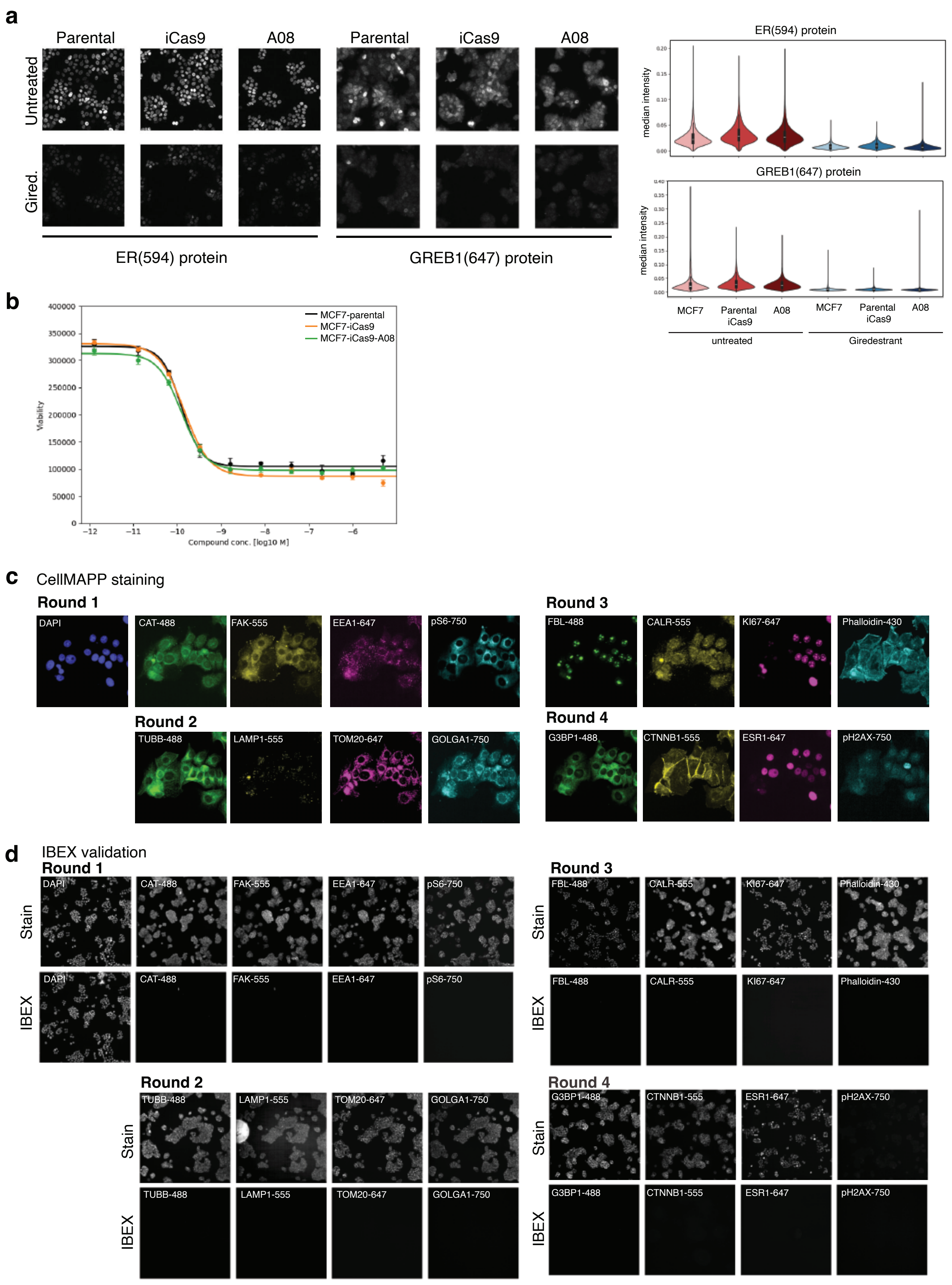

### Supplementary Figure 2

**a** Feature pre-processing pipeline

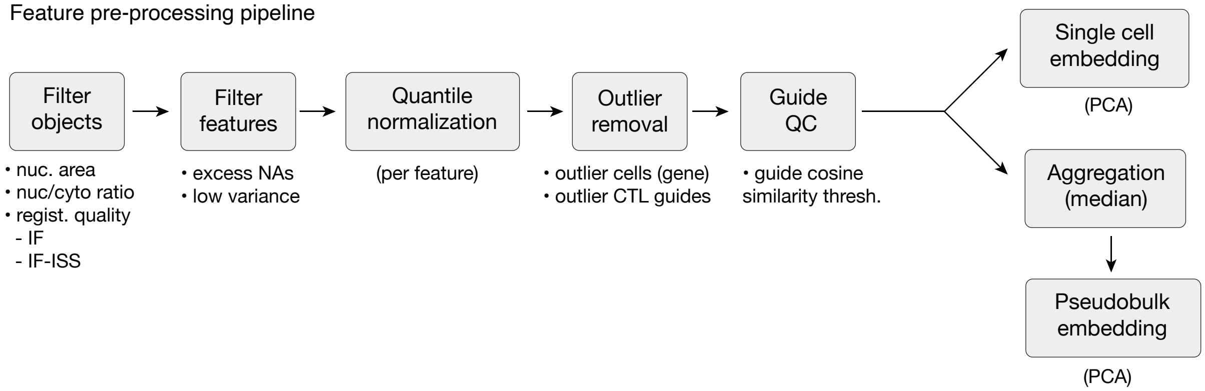


**b**

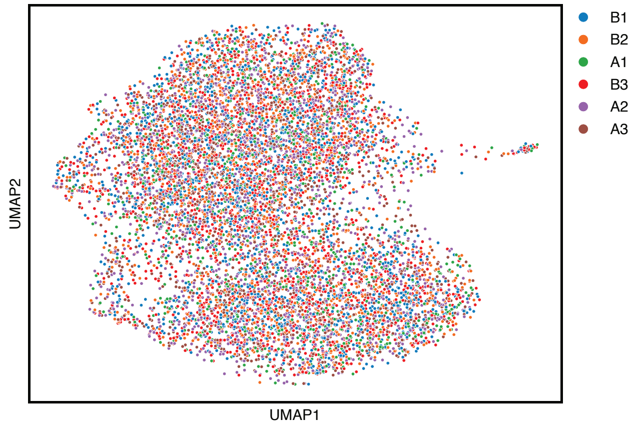

### Supplementary Figure 3

**a.**

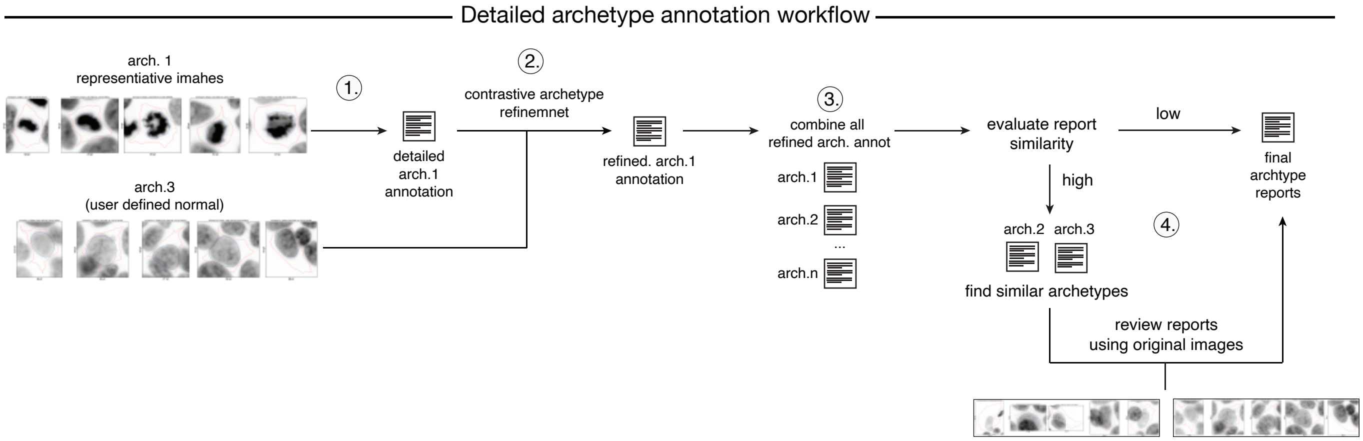

### Supplementary Figure 4

**Examples of archetypes captured across markers**

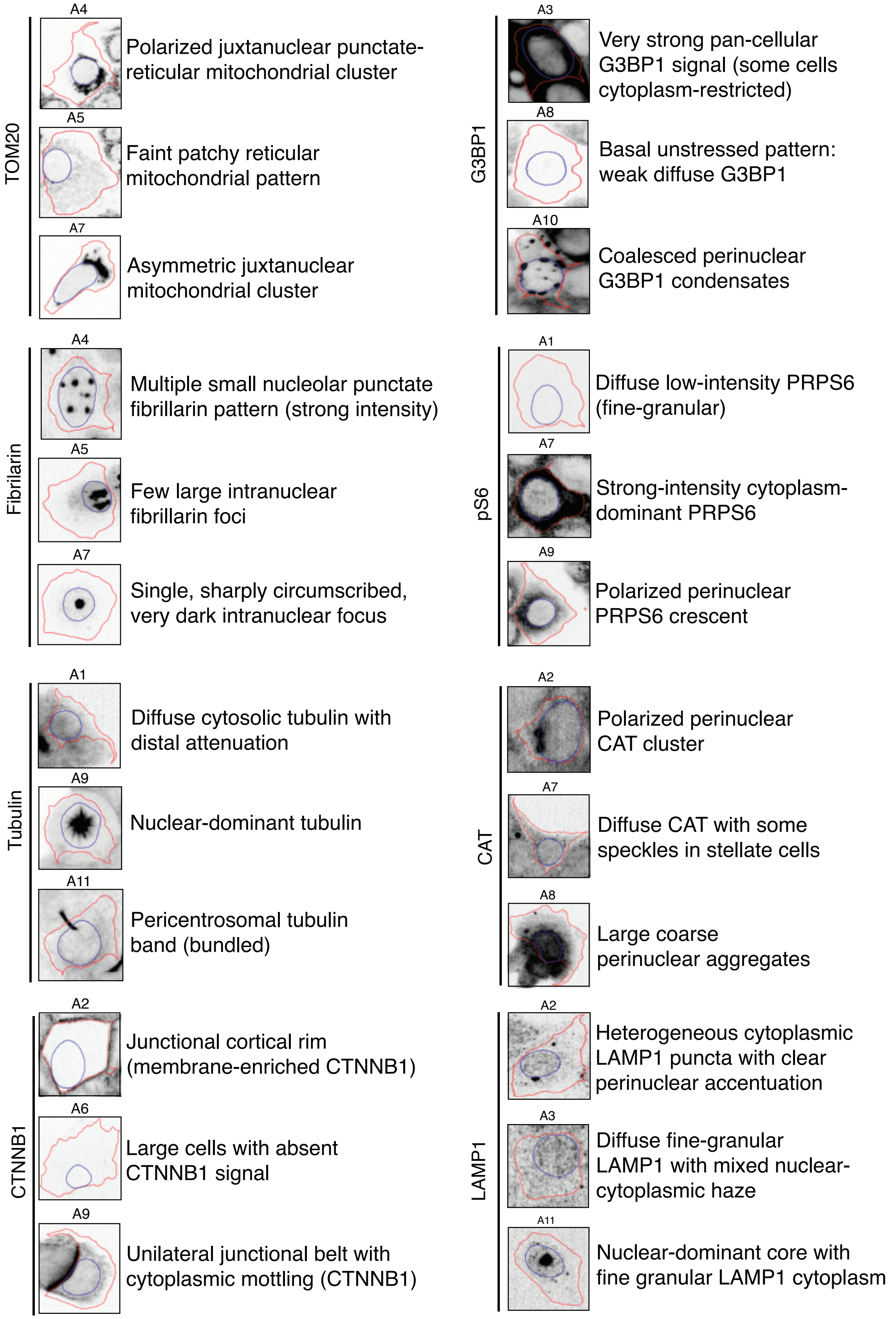

### Supplementary Figure 5

#### Select example of archetype MLLM archetype annotation

Representative archetype images depict top 5 cells with highest archetype weight used for archetype annotation

*Fibrillarin*

##### A4: Multiple small nucleolar punctate fibrillarin pattern (strong intensity)

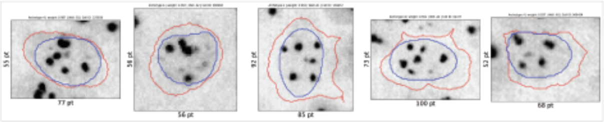

Fluorescence is predominantly strong, occasionally very strong, with consistent nuclear enrichment across cells.
**Signal manifests as multiple discrete nuclear puncta (coarse, rounded foci)** superimposed on a fine-granular nucleoplasmic haze. Cytoplasmic fluorescence is weak, diffuse, and uniform, without distinct puncta or perinuclear accumulations. **The puncta are confined to the nuclear compartment** with clear boundary adherence and no meaningful spillover.
Texture and distribution are characteristic: **several large nuclear foci per nucleus** with a fine granular background; **puncta are typically central without an obvious peripheral bias**.
Cytoplasm appears smooth to slightly speckled but remains low-intensity.

##### A7: Single, sharply circumscribed, very dark intranuclear focus

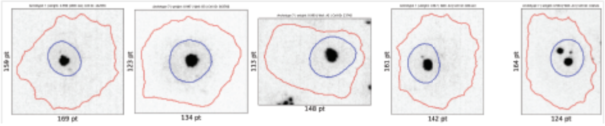

Archetype 7 shows a **dominant nuclear punctate pattern characterized by a single, sharply circumscribed, very dark intranuclear focus per cell**, set against low diffuse background. **Fluorescence intensity is strong to very strong at the focal punctum**, while the surrounding nucleoplasm displays faint, smooth-to-fine-granular haze and the cytoplasm exhibits only very weak, diffuse signal. C**onsistency is high: most cells contain one prominent nuclear punctum; occasional cells show a secondary small speckle** or up to three discrete foci, but all remain intranuclear. The **focal bodies are compact, smooth-edged, and well defined, positioned centrally** or slightly off-center without perinuclear or peripheral enrichment. No cytoplasmic puncta or edge-localized accumulations are observed. Texture and distribution are simple and clean, with no nuclear or cytoplasmic aggregates and no evident intensity gradients. **Cells are moderately large relative to baselines** 5 and 9, with smooth, **rounded/ovoid outlines** and minimal protrusions. Nuclei are moderate in size, round to mildly oval, and centrally placed.

*Tubulin*

##### A11: Pericentrosomal tubulin band (bundled)

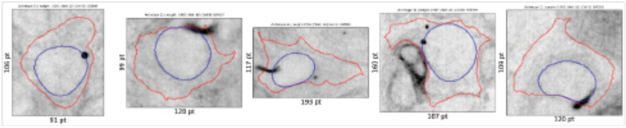

Fluorescence intensity shows a high dynamic range with strongly positive cytoplasmic regions contrasted against faint diffuse background. **Most cells display one or more intensely stained bands or compact foci,** while the remainder of the cytoplasm and the nuclear interior are weak to negative. Across cells, intensity is heterogeneous and often polarized to one side of the nucleus.
Localization is **predominantly cytoplasmic with perinuclear enrichment**: **dark, filamentous bands hug the nuclear margin and may extend toward the cell edge; occasional compact foci abut the nuclear border**. Nuclear signal is consistently low with no structured intranuclear pattern.
**Texture and distribution are characterized by bundled/striated elements consistent with filamentous structures, interspersed clumped aggregates, and scattered small puncta**. A gradient is common, with peak signal adjacent to the nucleus (and sometimes at the periphery) tapering to faint diffuse cytoplasm elsewhere.

##### A12: Edge-aligned cortical microtubule bundle

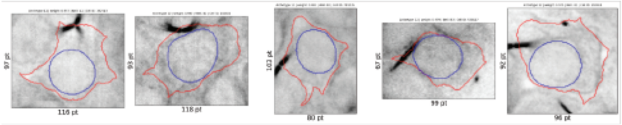

Fluorescence intensity is predominantly moderate with localized high-contrast foci: **most cytoplasm shows low, fine-granular signal, while one or more very dark, thick filamentous/linear bands along a single cell edge** exhibit peak intensity. Nuclear signal is consistently very weak to absent across cells.
**Localization is cytoplasmic with clear peripheral/cortical accentuation**. The **recurrent edge-tracking linear elements often abut and cross the cell boundary, continuing outside the cell outline; within the cell they form cortical arcs or bands**, while the remainder of the cytoplasm retains diffuse low signal. Perinuclear enrichment is not prominent in this grid.
Texture/distribution features a fine-granular diffuse background punctuated by **one to two thick linear bundles compatible with bundled microtubules;** puncta or coarse aggregates are not a dominant feature here. Signal heterogeneity arises chiefly from these high-contrast cortical bands.

##### A9: Nuclear-dominant tubulin

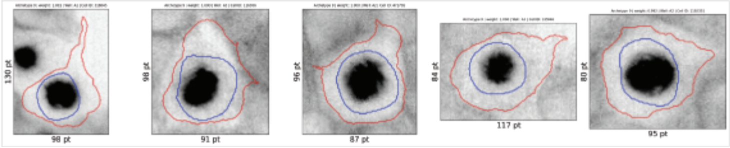

Archetype 9 displays uniformly high fluorescence with minimal cell-to-cell variability. Across all cells, **nuclear signal is the dominant feature—very strong, homogeneous, and smooth—forming the darkest region within the nuclear boundary**. Cytoplasmic fluorescence is consistently weaker to moderate and diffuse with fine granularity, showing clear perinuclear enrichment that tapers toward the cell periphery, producing a centripetal gradient. **No puncta, aggregates, or filamentous/reticular structures are evident; notably, the cytoplasm lacks discernible microtubule-like fibers**.

### Supplementary Figure 6

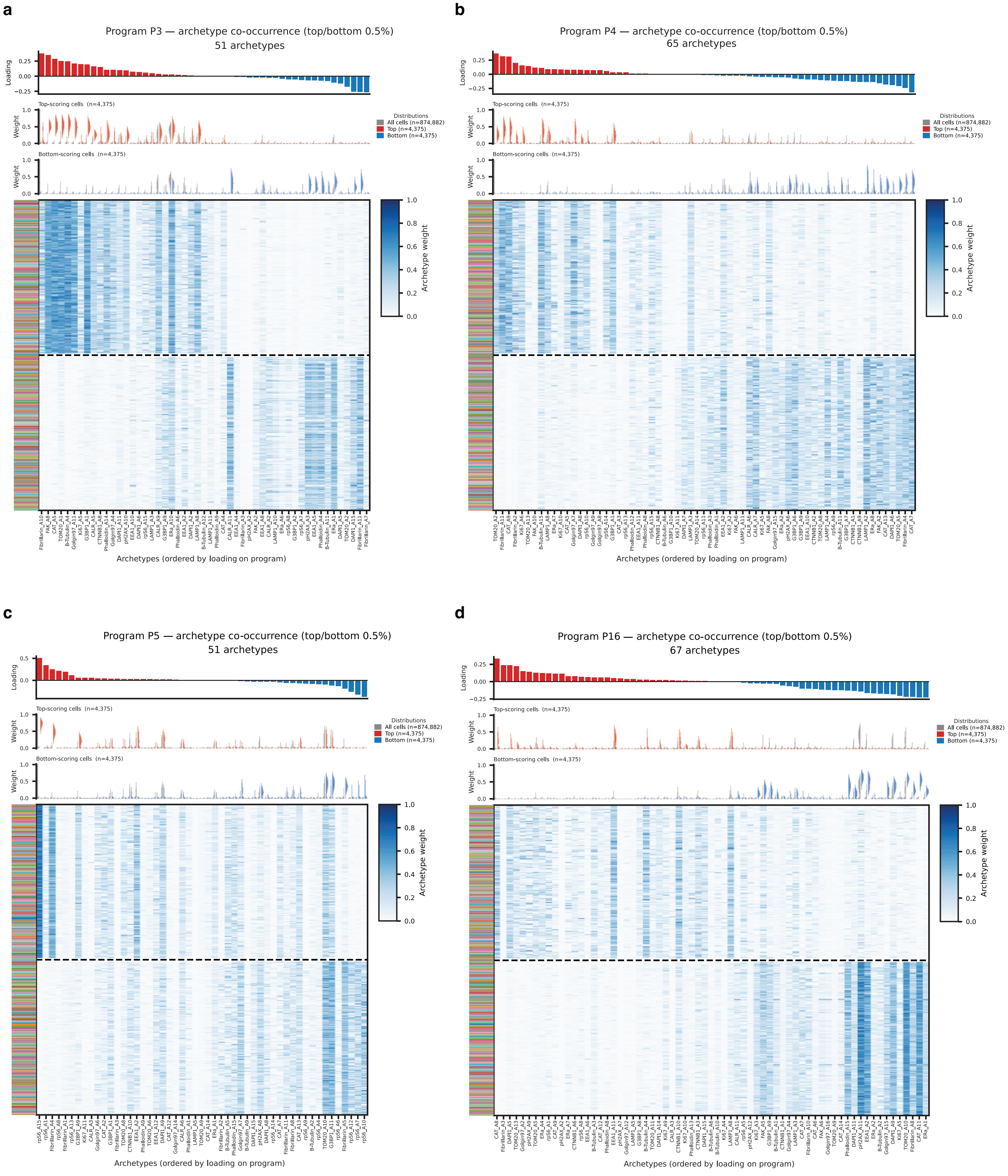

### Supplementary Figure 7

#### P7: Mitochondria dispersing into cytoplasmic reticular networks

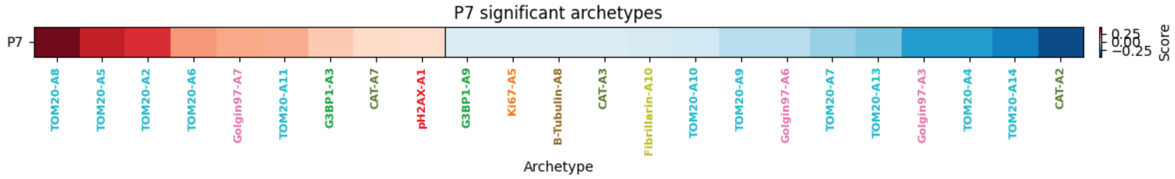


Representative images for archetypes in program:

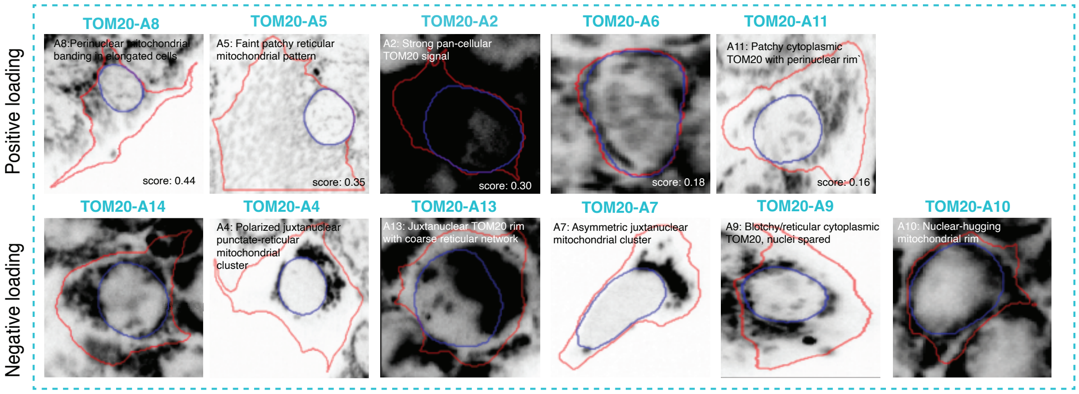


TOM20-dominated program that captures the axis of mitochondrial spatial distribution. The increase side promotes mitochondrial patterns where signal is distributed throughout the cytoplasm: reticular networks filling elongated cell bodies (A8), dispersed reticular filaments extending to the periphery (A5, the most reticular archetype), pan-cellular saturated signal (A2), and patchy cytoplasmic foci (A11). The decrease side features mitochondrial patterns where signal is tightly concentrated around the nucleus: polarized juxtanuclear clusters (A4, A7), tight perinuclear rims/rings (A10, A13), and dense one-sided accumulations (A7).

**Biological interpretation**: P7 captures a fundamental axis of mitochondrial network organization: dispersed/reticular vs. perinuclear-concentrated. Cells scoring high on P7 have mitochondria spread throughout the cytoplasm in reticular networks, while cells scoring low have mitochondria tightly clustered around the nucleus. This spatial redistribution has biological significance: perinuclear mitochondrial clustering is associated with quiescent or low-energy-demand states where mitochondria aggregate near the MTOC, while dispersed reticular networks are associated with active oxidative metabolism, mitochondrial fusion, and energy distribution to the cell periphery.

### Supplementary Figure 8

#### P11: Early Endosome Remodeling

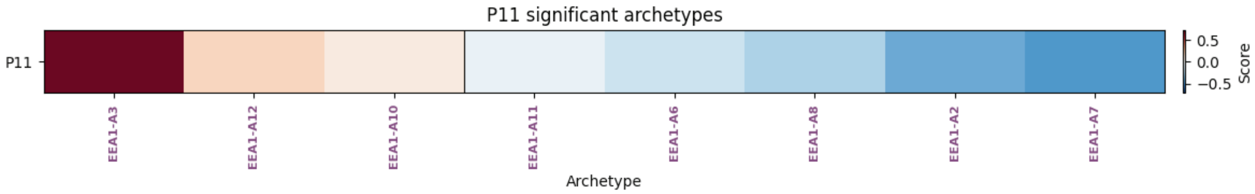


Representative images for archetypes in program:

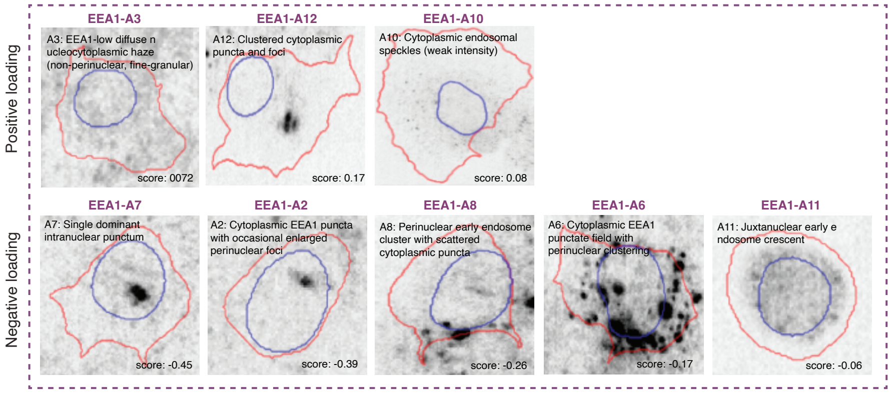


P11 is an EEA1-exclusive program that captures the **endosomal residence time / EEA1 occupancy axis**. Rather than simply "active vs. inactive" endosomal trafficking, P11 measures the abundance of EEA1+ early endosomal compartments — a quantity that can be altered by dysfunction at either end of the endosomal pathway. Both directions can represent pathological states; the "healthy" baseline of normal endosomal flux sits in the middle of the axis.

increasing (low EEA1 occupancy): EEA1-negative or EEA1-low cells. The dominant pattern (A3, 0.717) is essentially blank — no detectable early endosomes. The secondary contributor (A12, 0.169) shows a few large, coarse foci rather than numerous fine puncta, consistent with **enlarged, stalled endosomal compartments** — residual EEA1+ structures that have not been processed. The minor A10 contribution (0.079) shows sparse, weak speckles in well-spread cells. The coexistence of A3 (blank) with A12 (few swollen foci) is a dysfunction signature, not just a shutdown signature: normal trafficking has ceased, but residual compartments that persist are enlarged rather than cleared.

decreasing (high EEA1 occupancy): Cells with abundant EEA1+ compartments across a spectrum of patterns: dense cytoplasm-filling punctate fields (A6), perinuclear clusters (A8), distributed puncta with perinuclear foci (A2), organized perinuclear crescents (A11), and over-nuclear endosomal signal (A7). These represent high EEA1 membrane occupancy — either from active high-throughput endocytic trafficking or from impaired EEA1 release / endosomal maturation causing compartment accumulation.

**Biological interpretation**:
(1) P11 increase — recycling pathway disruption:** When the recycling arm of the endosomal pathway is impaired (e.g., disruption of Rab-family GTPases controlling tubular recycling endosome formation or cargo return to the plasma membrane), cells may compensate by reducing endocytic uptake — a feedback mechanism to maintain surface receptor homeostasis when recycling cannot return internalized receptors. This leads to fewer EEA1+ compartments forming, producing the A3 pattern (blank). The A12 pattern (few large stalled foci) represents residual endosomes that formed but cannot be cleared through the recycling arm — they swell and persist as enlarged, stalled compartments. Any perturbation that blocks endosomal recycling, impairs recycling endosome biogenesis, or reduces endocytic input is predicted to increase P11.
(2) P11 decrease — maturation/degradation pathway disruption:** When the early-to-late endosome maturation pathway is impaired (e.g., disruption of the PI(3)P PI(3,5)$P_2$ lipid conversion, Rab5-to-Rab7 switch, or MVB formation), PI(3)P accumulates on endosomal membranes. Since EEA1 binds PI(3)P via its FYVE domain, EEA1 becomes trapped — it cannot be released from endosomal membranes. Simultaneously, endosomes swell and accumulate because they cannot mature into late endosomes or fuse with lysosomes.

### Supplementary Figure 9

#### P5: Biosynthetic shutdown across nucleus and cytoplasm

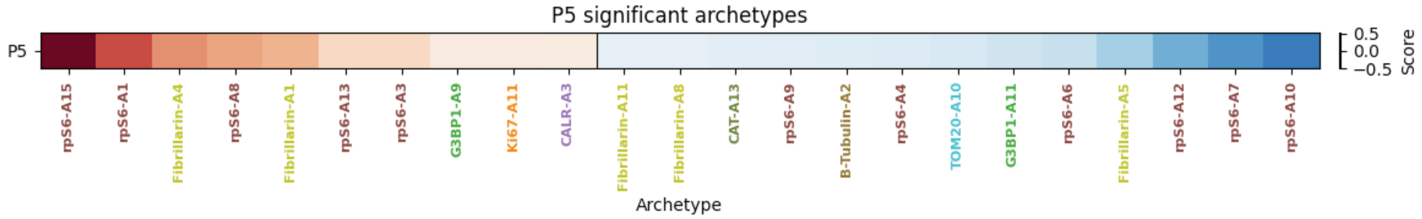


Representative images for archetypes in program:

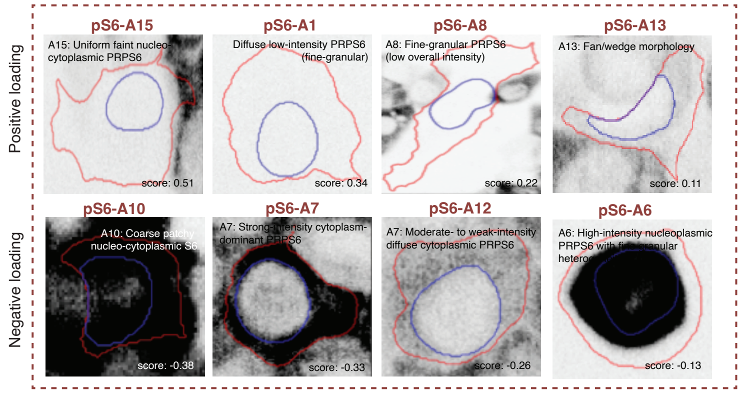


Near-absent or uniformly faint rpS6 signal across morphologically varied cells (rounded, spread, irregular).

Decreasing: All forms of strong phospho-rpS6 — cytoplasm-dominant perinuclear rings (A7), coarse patchy nucleo-cytoplasmic (A10), nuclear-dominant (A6), cortical patches (A4), perinuclear crescents (A9) — indicating comprehensive mTOR-S6K pathway inactivity

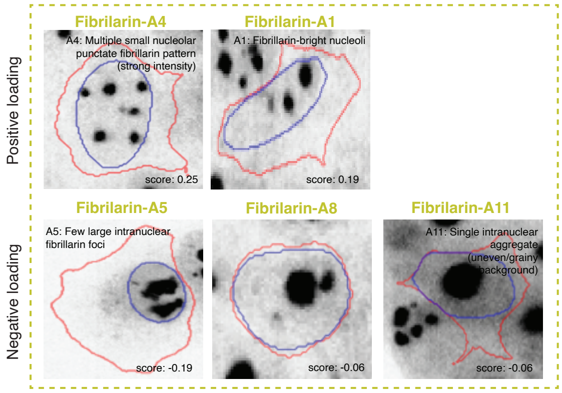


Multiple small nucleolar foci (Fibrillarin_A4: several discrete puncta per nucleus in small-to-medium cells) — nucleoli that have not coalesced into the large, high-through-put factories characteristic of actively growing cells.

Few large coalesced nucleoli (Fibrillarin_A5, A7, A11) — the mature interphase pattern associated with concentrated, high-output ribosome assembly.

**Biological interpretation:** P5 defines a translationally quiescent state where nucleolar organization and phospho-rpS6 status are coherently suppressed. In actively growing cells, nucleoli fuse via liquid-liquid phase separation into few large bodies that serve as high-throughput ribosome factories, and mTORC1 drives rpS6 phosphorylation to promote ribosomal protein mRNA translation. P5 opposes both: nucleoli remain small and dispersed (low coalescence, low rRNA output), and pS6 is absent (no translational drive). This coherence is notable because it demonstrates that single-cell imaging can resolve the coordinated downregulation of the ribosome biogenesis pipeline across two independent compartments — nuclear (nucleolar architecture) and cytoplasmic (pS6 status) — within the same morphological program.

### Supplementary Figure 10

#### P16: CAT+ stress-adaptive trafficking remodeling state

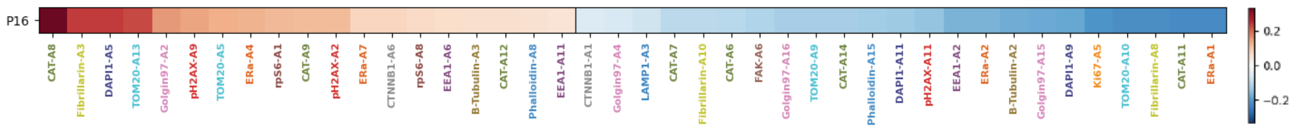


Representative images for archetypes in program:

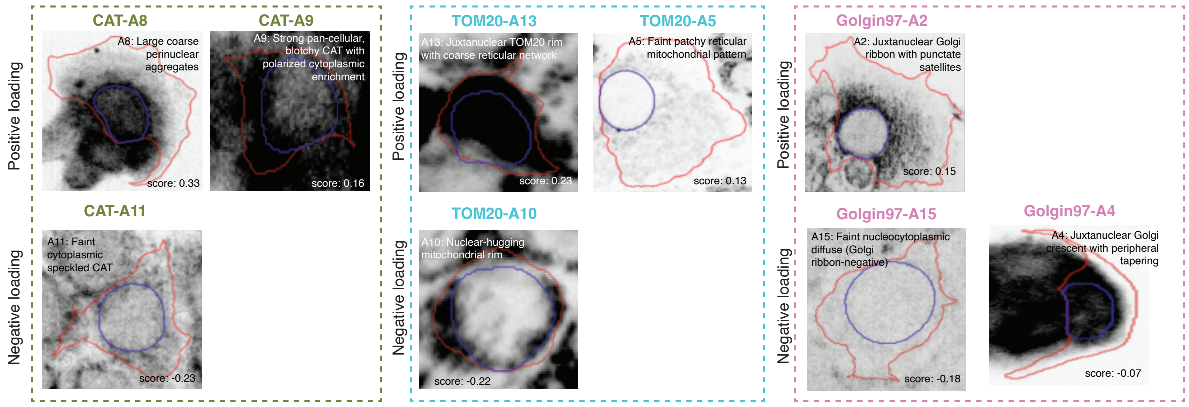


**Peroxisomes — massive expansion:** Three catalase archetypes increase (CAT_A8 at 0.332, CAT_A9 at 0.115, CAT_A12 at 0.062) while four low-intensity patterns decrease. The coarse perinuclear aggregates (A8) indicate physically enlarged/clustered peroxisomes, not just increased catalase expression. This is peroxisomal biogenesis — new peroxisomes being generated and concentrated perinuclearly.

**Mitochondria — network loosening:** The tight perinuclear band (TOM20_A10, decrease) opens into a looser reticular rim (TOM20_A13, increase) while maintaining network connectivity (both A13 and A5 show reticular morphology, not punctate/fragmented). The distributed cytoplasmic pattern (TOM20_A9) also decreases — mitochondria remain perinuclear but in a more extended, less tightly packed configuration. This loosening may create more surface area for ER-mitochondria contacts and for metabolic exchange between mitochondria and the expanding peroxisomal compartment.

**Golgi — maturation into full ribbon:** The compact crescent (Golgin97_A4, decrease) is replaced by the most organized ribbon pattern (Golgin97_A2, increase) — a mature juxtanuclear ribbon/cluster with associated transport vesicle satellites in larger, spread cells. This represents Golgi maturation and expanded secretory capacity, potentially to support the membrane and protein demands of peroxisomal and organelle biogenesis.

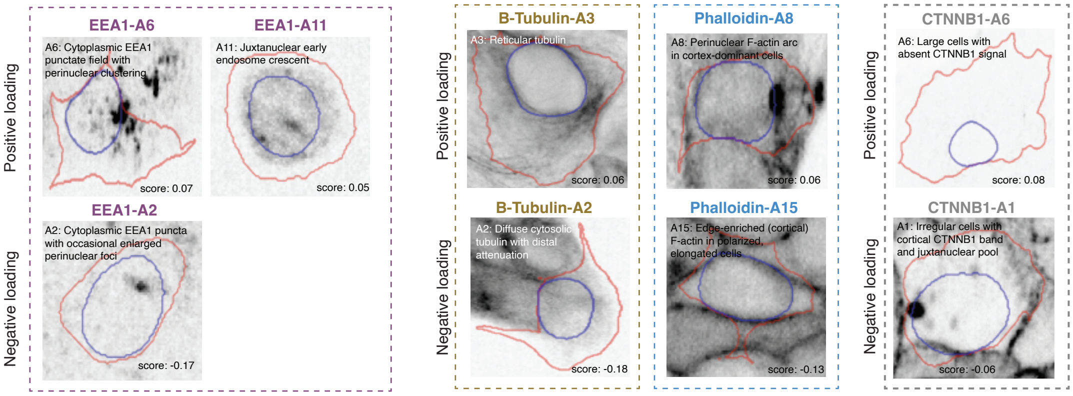


**Endosomes — shift to active high-throughput trafficking:** Enlarged, stalled perinuclear foci (EEA1_A2, decrease –0.168) are replaced by dense punctate fields with perinuclear clustering (EEA1_A6, increase 0.067) and organized juxtanuclear crescents (EEA1_A11, increase 0.053). The shift from few large endosomes to many small organized ones indicates more active, higher-throughput endosomal trafficking — consistent with increased membrane cycling demands from organelle biogenesis.

*Cytoskeleton — perinuclear support:** Cortical F-actin in elongated polarized cells (Phalloidin_A15, decrease) is replaced by perinuclear F-actin arcs (Phalloidin_A8, increase). Diffuse depolymerized tubulin (B-Tubulin_A2, decrease) is replaced by reticular organized microtubules (B-Tubulin_A3, increase). The cytoskeleton is reorganizing to support the perinuclear organelle expansion — actin concentrated perinuclearly, microtubules maintaining organized network for vesicular transport.

- **CTNNB1_Archetype6** (increase, 0.078): **Large cells with absent CTNNB1 signal.** Very low, diffuse β-catenin; large cells with eccentric nuclei. Indicates loss of adherens junctions.

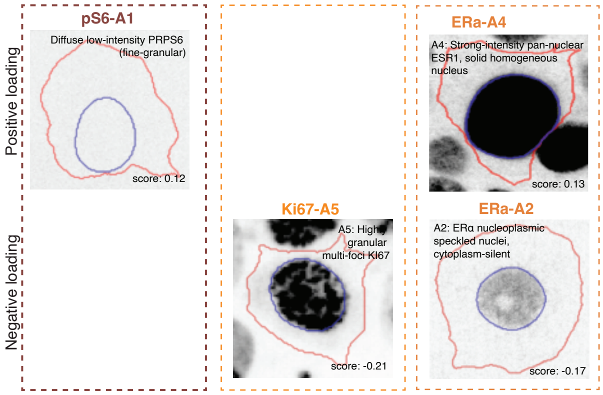


Strong ERα expression suggests hormone-responsive transcriptional engagement, while concurrent loss of Ki67 and phospho-rpS6 indicates exit from proliferation and translational downregulation

**Biological interpretation:** P16 captures a coordinated perinuclear organelle expansion and maturation program spanning numerous compartments. Peroxisomes expand massively (coarse catalase-positive aggregates), mitochondria loosen from a tight perinuclear band to a reticular rim (TOM20), the Golgi matures toward a full ribbon with transport vesicle satellites (Golgin97), and early endosomes shift from stalled foci to active punctate trafficking (EEA1). The cytoskeleton reorganizes in support — microtubules adopt a reticular network (β-tubulin) while F-actin forms a perinuclear arc (phalloidin). The cells are not dividing (Ki67 decrease) and have low translation (rpS6 low), suggesting the organelle expansion is occurring through redistribution and maturation of existing compo-nents rather than de novo protein synthesis. Together, P16 describes a cell state that centralizes trafficking and redox capacity while shifting from growth toward structural maintenance and detoxification

### Supplementary Figure 11

#### P10: Quiescent metabolic reprogramming

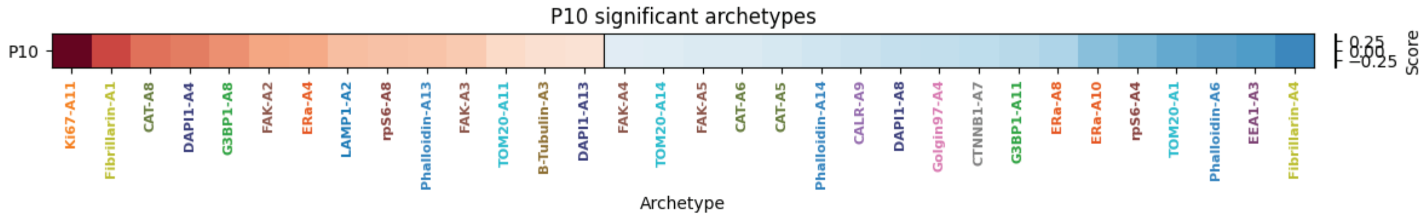


Representative images for archetypes in program:

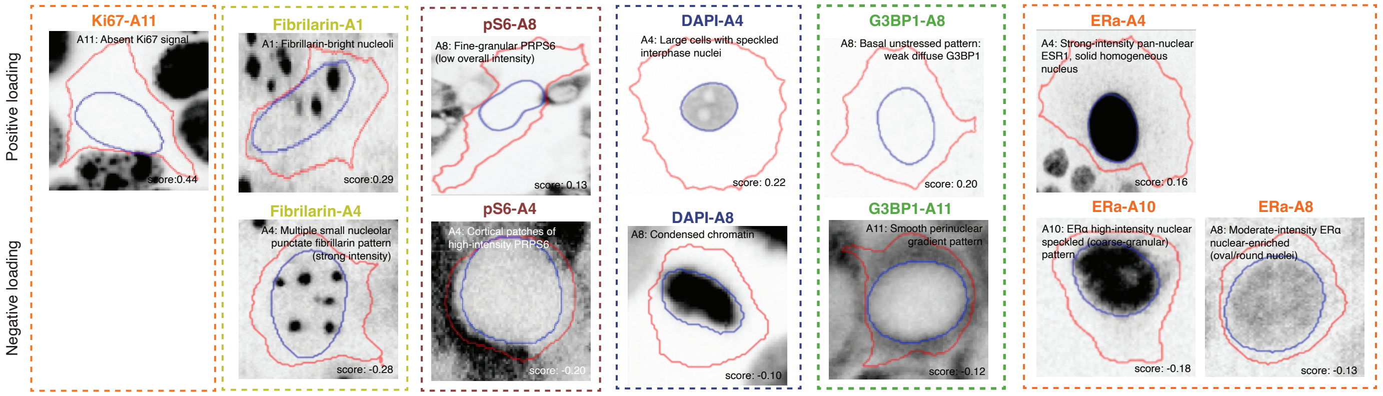


G0 quiescent large cells with few modest nucleoli, low translation (rpS6 low) cells are not actively

decrease in actively cycling small cells

absence of stress (G3BP1 essentially blank)

Smooth, homogeneous ERα fill (A4) — as opposed to the speckled/granular pattern (A10) — may reflect ERα uniformly distributed across the nucleoplasm in a tonically occupied or unliganded state, rather than concentrated at active transcriptional foci.

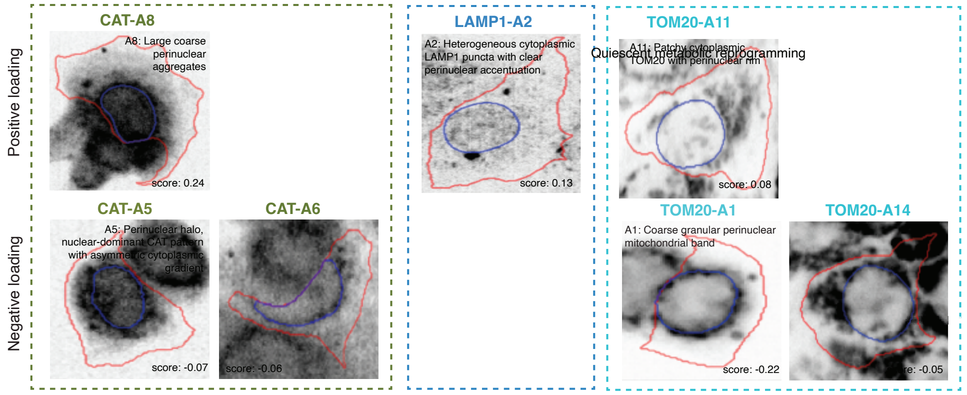


**Biological interpretation**: P10 captures a deeply quiescent, hormone-responsive cell state in which the growth machinery has been shut down but organelle-level housekeeping functions persist. Both ribosome biogenesis (few modest nucleoli) and translation (low rpS6) are suppressed — these cells are not actively making protein. Yet they maintain robust peroxisomal metabolism (catalase), an active lysosomal/autophagy system, and high ERα levels, suggesting they retain the capacity to respond to hormonal signals and manage oxidative/catabolic homeostasis. The cells are large and spread but with low FAK signaling (A2, A3), suggesting stable, mature adhesions that no longer require active focal adhesion turnover.

Organelle-level housekeeping functions persist. Cells maintain robust peroxisomal metabolism (catalase), an active lysosomal/autophagy system, suggesting they manage oxidative/catabolic homeostasis.
P10 cells lack the centrosome-driven radial organelle architecture associated with proliferation and migration. Instead, organelles are more loosely distributed.

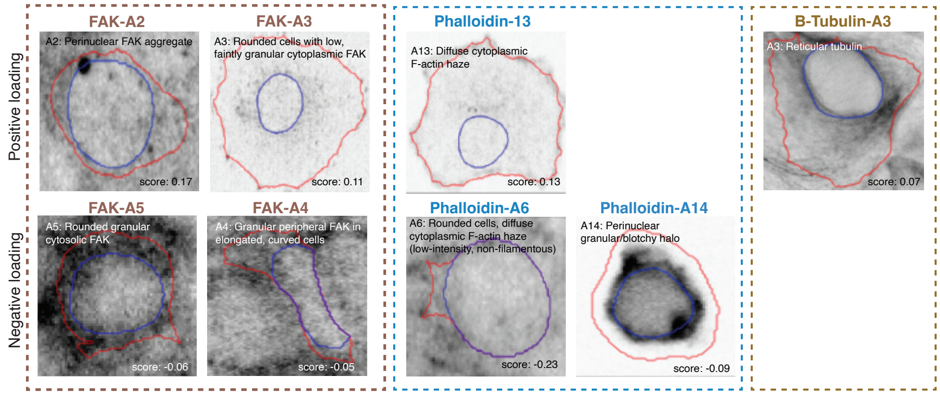


The low FAK despite large spread morphology suggests these cells have established stable, mature adhesions that no longer require active signaling — consistent with long-term quiescent cells that have settled into a stable substrate attachment

**Metabolic interpretation**: The marker profile suggests P10 cells have shifted from the anabolic glycolytic metabolism typical of proliferating cancer cells to a **catabolic maintenance state**. The combination of high peroxisomal mass (CAT_A8), low mTOR/translation (rpS6 low), and active lysosomes (LAMP1_A2) points to a metabolic program centered on fatty acid oxidation (FAO) and autophagy rather than glycolysis and biosynthesis. Peroxisomes perform β-oxidation of very long chain fatty acids, generating $H_2O_2$ that is detoxified by the abundant catalase; the resulting medium-chain products are shuttled to mitochondria (TOM20_A11, moderate and distributed) for complete oxidation via the TCA cycle. Low mTOR suppresses the anabolic arm (lipid synthesis, nucleotide synthesis, glycolytic gene expression) while permitting autophagy — the active lysosomes likely reflect constitutive quality-control autophagy recycling damaged organelles and long-lived proteins during prolonged quiescence. The absence of stress markers (G3BP1 blank, no strong γH2AX) indicates that ROS generated by peroxisomal FAO is effectively managed by the high catalase levels. In the context of cancer, this FAO-dependent, mTOR-low, autophagy-active, quiescent profile is consistent with **therapy-resistant persister cells** or **dormant disseminated tumor cells** — cells that survive proliferation-targeting therapies by exiting the cell cycle and switching to a catabolic fuel source.

### Supplementary Figure 12

#### P4: Loss of spatial compartmentalization with organelle compaction

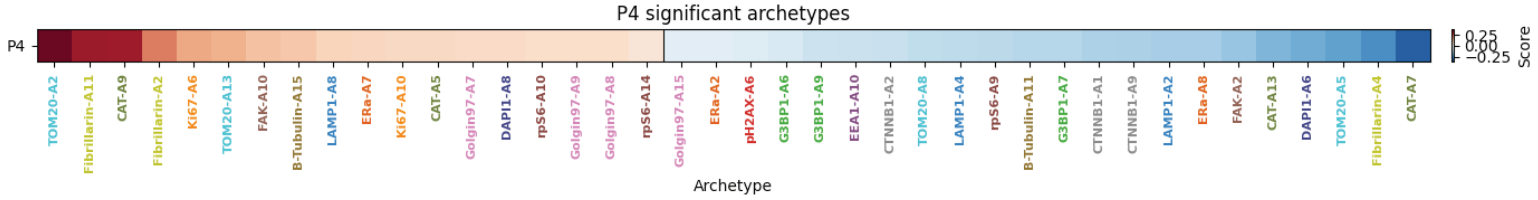


Representative images for archetypes in program:

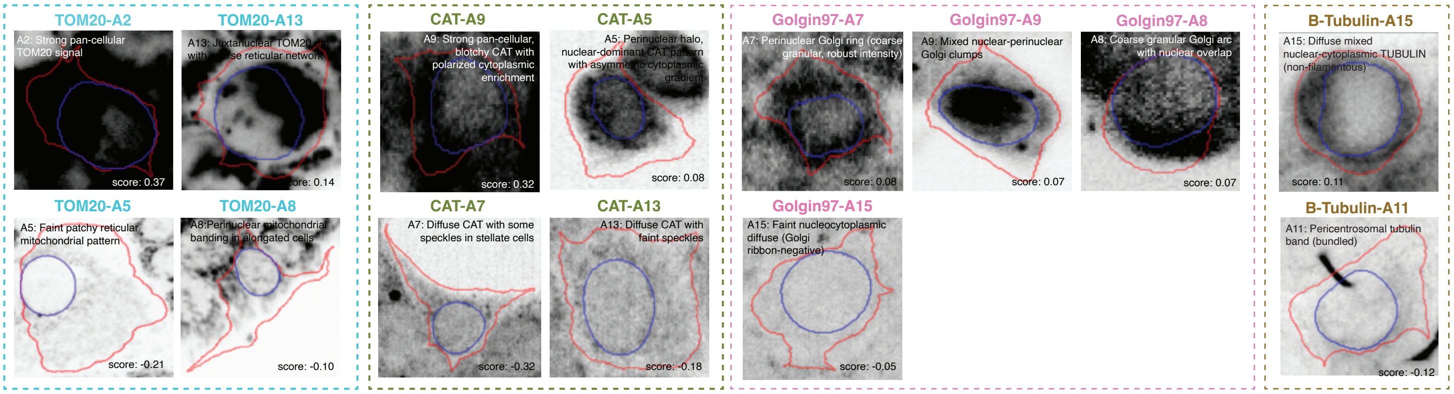


The defining morphological features — observed directly across the dominant archetypes — are: small, rounded cells with high N/C ratio; pan-cellular undifferentiated organelle signals (mitochondria, peroxisomes. Pan-cellular undifferentiated organelle signals — TOM20_A2 (uniform mitochondrial filling) and CAT_A9 (uniform peroxisomal filling) — reflecting loss of spatial organization rather than increased organelle content, as organelles redistribute uniformly when cells compact.

Diffuse non-filamentous tubulin (B-Tubulin_A15) — depolymerized microtubule network.

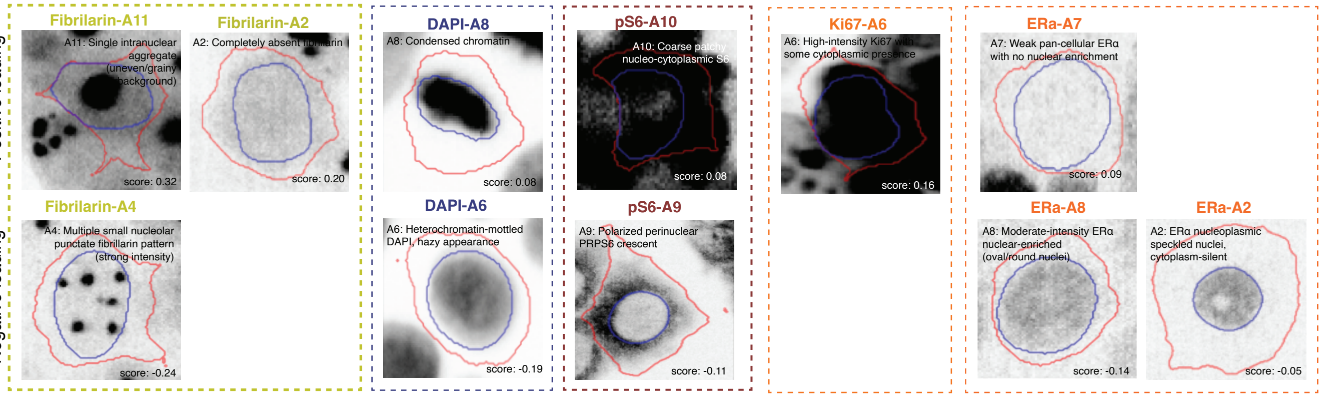


A two-stage nucleolar disassembly sequence: Fibrillarin_A11 (single condensed nucleolar aggregate — pre-dissolution compaction) followed by Fibrillarin_A2 (completely absent fibrillarin — full nucleolar dissolution).

chromatin condensation

Weak pan-cellular ERα without nuclear enrichment (ERa_A7).

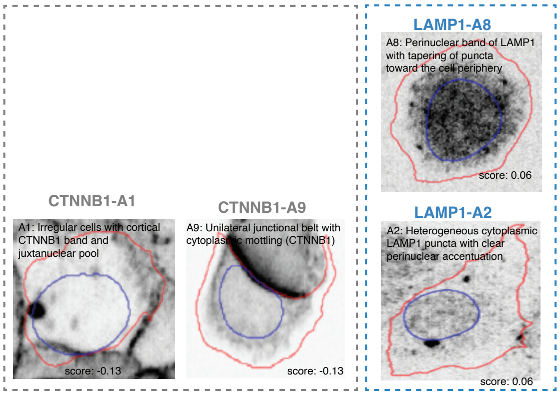


dissolution of cell-cell junctions and focal adhesions

Decreasing archetypes: All forms of interphase spatial organization — multiple active nucleoli (Fibrillarin_A4), organized pericentrosomal microtubule bundles (B-Tubulin_A11), junctional and cortical β-catenin in three distinct patterns (CTNNB1_A9, A1, A2), organized lysosomal distributions (LAMP1_A2, A4), polarized perinuclear rpS6 crescent (rpS6_A9), and spread stellate cell morphology (CAT_A7). The program comprehensively opposes the adherent, polarized, spatially structured interphase state.

**Biological interpretation**: The most parsimonious interpretation is that P4 captures a compact transitional state — cells that have rounded up, lost substrate adhesion and cell-cell junctions, dissolved their interphase organelle architecture, and shut down nucleolar transcription. Mitosis (prophase/prometaphase) is the leading biological interpretation: the nucleolar condensation-to-dissolution sequence corresponds to known prophase nucleolar behavior; chromatin condensation (DAPI_A8) is a hallmark of mitotic entry; Ki67 cytoplasmic redistribution (Ki67_A6) occurs when the nuclear envelope breaks down; diffuse non-filamentous tubulin (B-Tubulin_A15) represents the depolymerized state between interphase MT array disassembly and spindle formation; and junction dissolution is expected as cells round up for division. However, the dominant archetypes (TOM20_A2, CAT_A9, Fibrillarin_A11) describe the physical configuration — compact, organelle-filled, spatially undifferentiated — rather than the cell cycle phase directly. This same morphological signature could in principle also arise from de-adhesion, early apoptotic rounding, or other transitions that involve cell compaction and loss of interphase architecture. The Ki67_A6 and DAPI_A8 contributions favor mitosis over these alternatives, but they are moderate contributors (0.155 and 0.079) rather than the dominant signal.

### Supplementary Figure 13

#### P3: Proliferative-migratory activation with centrosomal polarity and secretory expansion

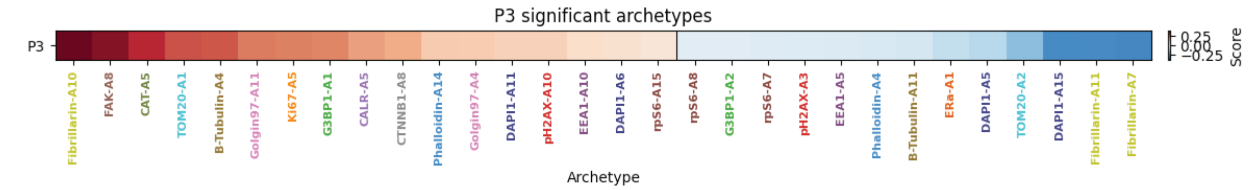


Representative images for archetypes in program:

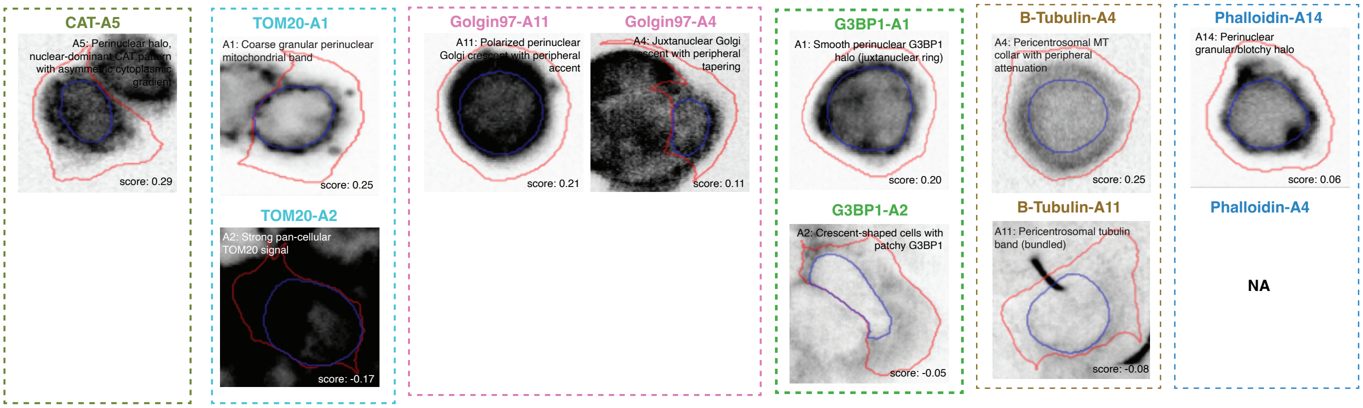


The defining feature is a **centrosomal organizing hub**:TOM20_A1, B-Tubulin_A4, CAT_A5, Phalloidin_A14, and G3BP1_A1 all show concentric perinuclear patterns that are independently consistent with MTOC-anchored radial organization. The Golgi hypertrophy (Golgin97_A11) was more dramatic than the report text suggested — near-saturated signal filling much of the cytoplasm rather than a compact ribbon.

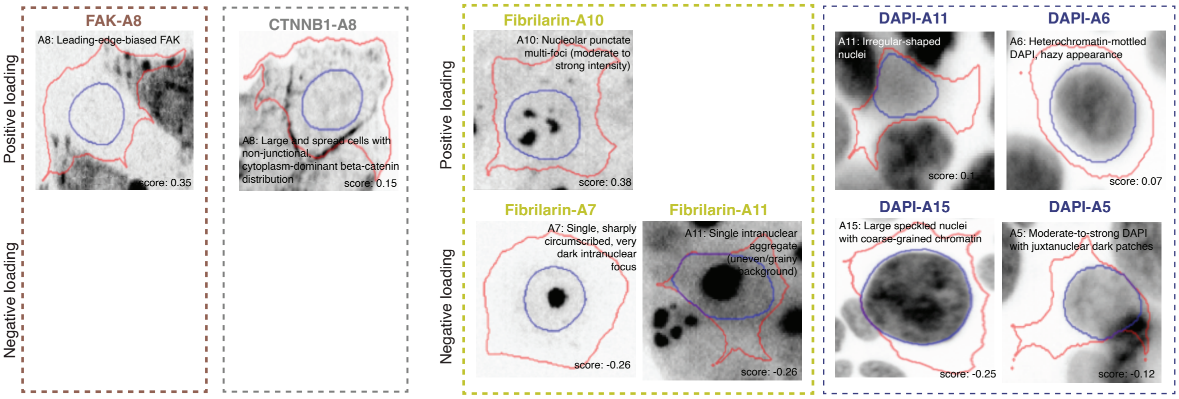


Leading-edge FAK polarization (directed migration) Non-junctional cytoplasmic β-catenin (loosened cell-cell contacts). Irregular/lobulated nuclear shapes (migration or S/G2).

Fibrillarin contrast (multi-foci vs. single focus)
What is decreasing: resting cells with single consolidated nucleoli, very large heterochromatin-rich nuclei. The program opposes both large quiescent cells (Fibrillarin_A7, DAPI_A15) and small compact quiescent cells (Fibrillarin_A11), suggesting the activated state it captures is distinct from quiescence regardless of cell size.

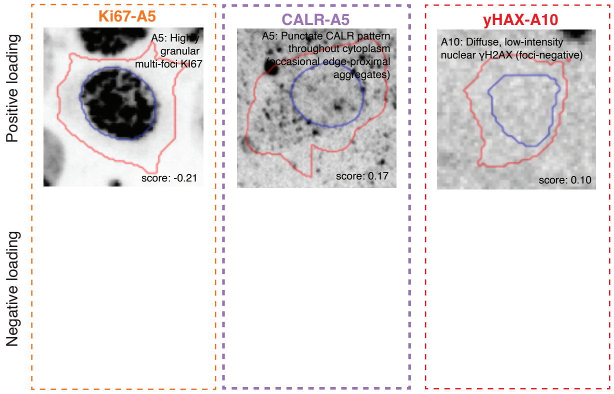


Ki67 macrofoci (S/G2 proliferation)

Ccattered ER exit site puncta

No DNA damage (foci-negative γH2AX)

**Biological interpretation**: P3 defines a proliferative-migratory activation state. The centrosomal organizing hub — where mitochondria, microtubules, peroxisomes, actin, and ribosomes are all arranged concentrically around the MTOC — is a hallmark of cells that have established functional centrosome-based polarity. The expanded, near-saturated Golgi (not a compact ribbon but a hypertrophic mass extending toward the periphery) suggests Golgi ministack dispersal or genuinely increased Golgi mass. The scattered CALR puncta (ER exit sites) feeding into this expanded Golgi complete a picture of an activated secretory pathway. The non-junctional cytoplasmic β-catenin indicates cells that have loosened adherens junctions — consistent with migratory cells that have partially disengaged from neighbors. The irregular/lobulated nuclear shapes may reflect nuclear deformation during confined migration or S/G2-associated nuclear remodeling. Notably, this activation occurs without detectable DNA damage (foci-negative γH2AX) or proteotoxic stress (no G3BP1 stress granules), suggesting a physiological proliferative-migratory program rather than a stress response. Strictly nuclear Ki67 macrofoci place these cells in late S or G2 phase. Overall, P3 likely captures an aggressive or activated epithelial phenotype combining cell cycle progression, directed migration, secretory hypertrophy, and centrosome-based polarity.

### Supplementary Figure 14

**a.**

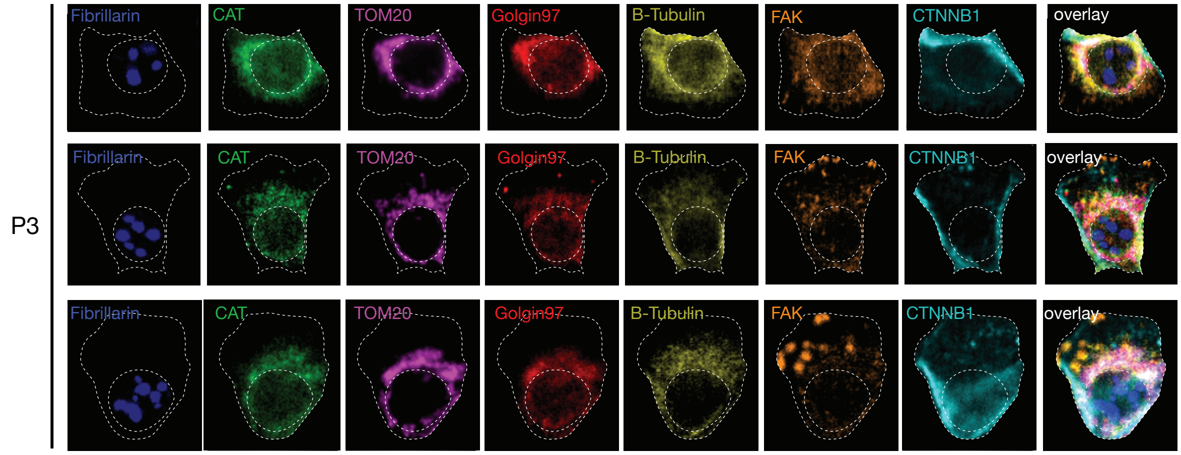


**b.**

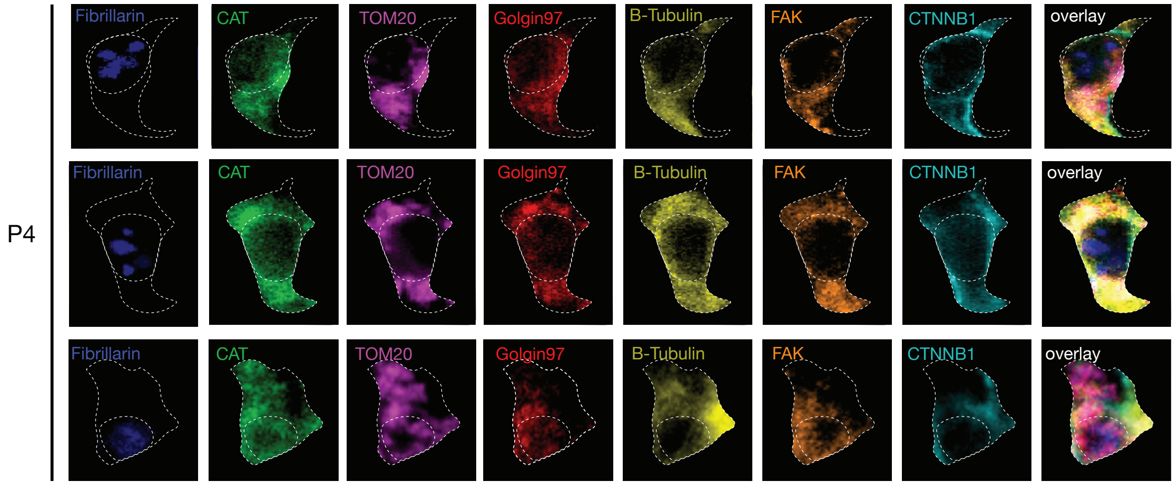

### Supplementary Figure 15

#### P18: Cortically organized stress-responsive state

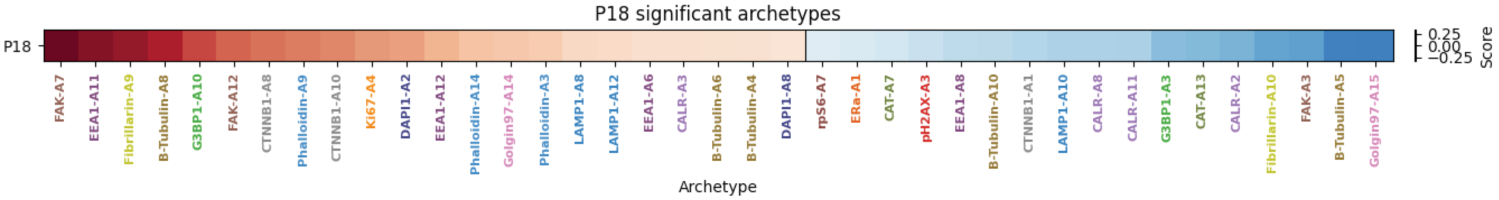


Representative images for archetypes in program:

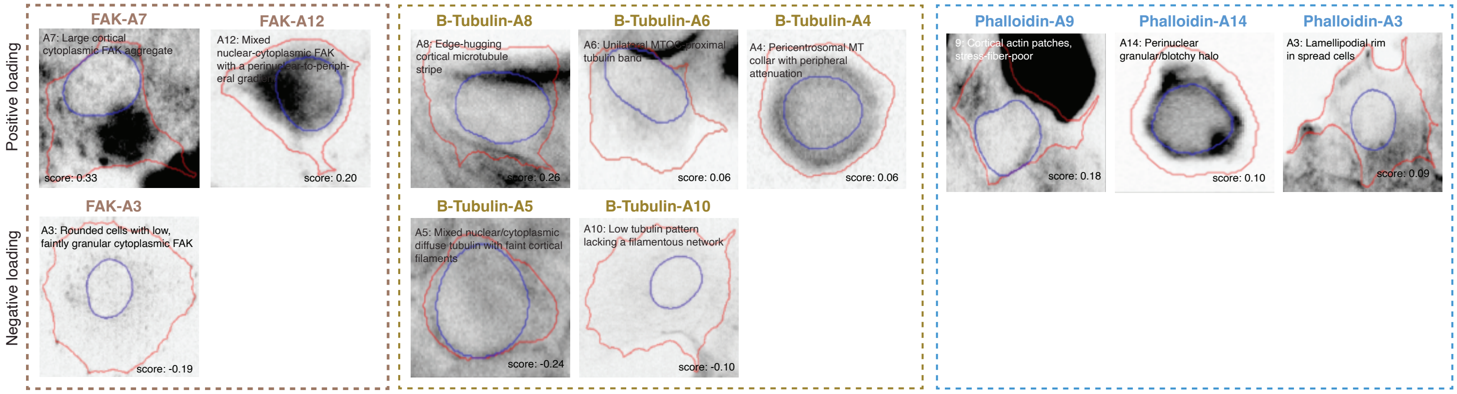


P18 captures a cell state characterized by cortical organization.
Iincrease of cortical FAK aggregates (A7), cortical microtubule stripes (B-Tubulin_A8), cortical actin patches (Phalloidin_A9),
The decrease of rounded cells with low FAK, spindle cells with low CALR/LAMP1, and diffuse tubulin indicates a departure from both compact and elongated morphologies toward a spread, cortically organized state.

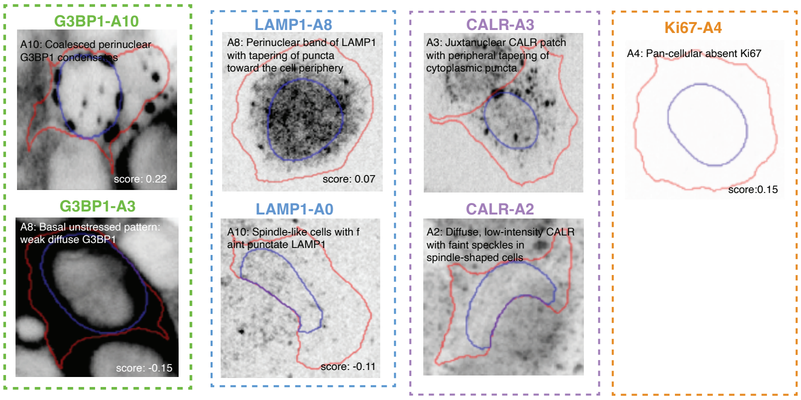


The G3BP1 condensates (A10) suggest stress granule formation

Opposition to spindle cell morphology (CALR_A2, LAMP1_A10)

Absent Ki67

**Biological interpretation**: P18 may capture a polarized, cortically organized cell state with features of cellular stress. The cortical enrichment of FAK, tubulin, and actin suggests cells with strong peripheral organization — possibly polarized for migration or responding to mechanical cues. The G3BP1 condensates are the most distinctive feature, indicating stress granule assembly. Combined with absent Ki67 this may represent quiescent cells under stress that have reorganized their cortical cytoskeleton and initiated stress granule formation. The opposition to spindle cell morphology (CALR_A2, LAMP1_A10) and to diffuse G3BP1 (A3) reinforces that this is a distinct, cortically organized, stress-responsive state.

### Supplementary Figure 16

#### P6: Estrogen receptor silencing

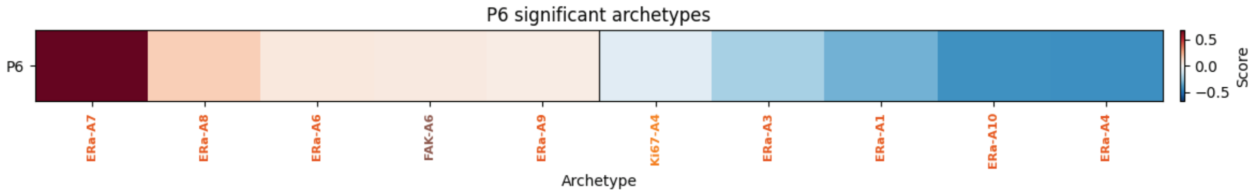


Representative images for archetypes in program:

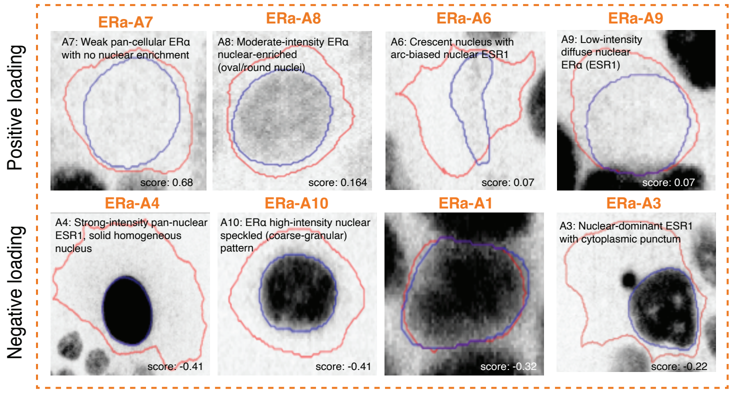


P6 is almost exclusively an ERα program, capturing the axis of estrogen receptor expression intensity. It promotes weak/absent ERα signal and opposes strong, nuclear-enriched ERα patterns. This is a single-marker program that cleanly separates ER-low from ER-high cells.

**Biological interpretation**: P6 directly captures the ERα expression gradient. Cells scoring high on P6 are ER-negative or ER-low, while cells scoring low are ER-high with active nuclear estrogen signaling. This is consistent with a fundamental axis of breast cancer cell heterogeneity (ER+ vs. ER– status).

### Supplementary Figure 17

#### P17: G0 / Quiesencent-like

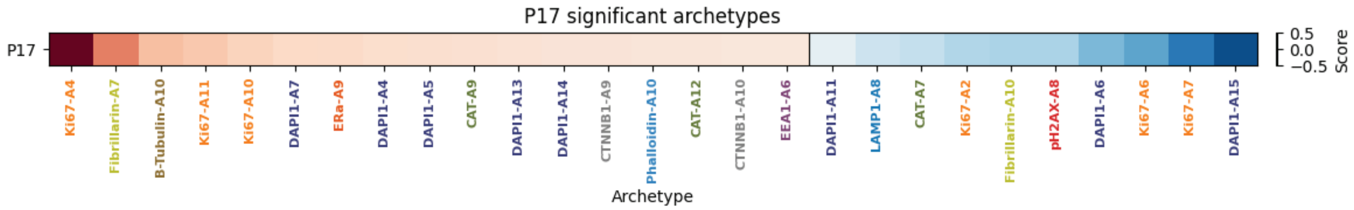


Representative images for archetypes in program:

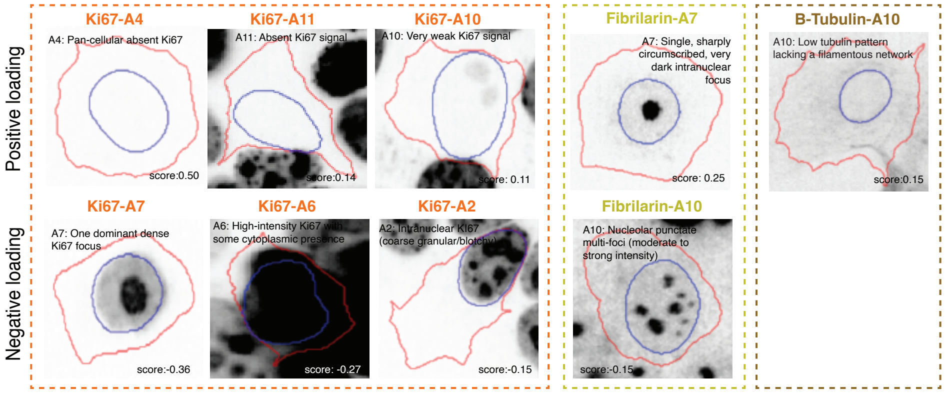


Three Ki67 archetypes indicating absent/very weak Ki67 are increased, while three Ki67 archetypes indicating active proliferation are decreased

Quiescent cells have single nucleoli (Fibrillarin_A7)

Depolymerized tubulin (no mitotic spindle, reduced cytoskeletal dynamics)

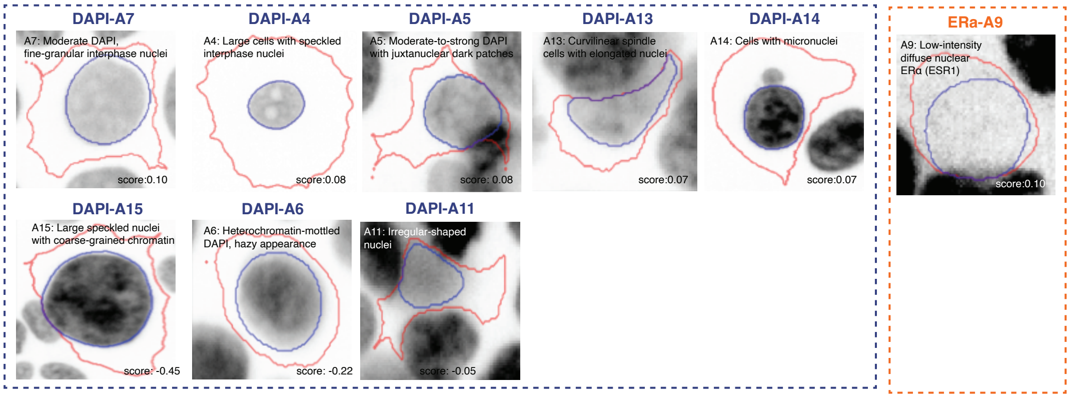


Fine-granular interphase chromatin

**Biological interpretation**: P17 captures the quiescence axis — specifically, cells that have exited the cell cycle into G0. The hallmarks are: absent Ki67 (no proliferation), single nucleolus (reduced ribosomal demand), depolymerized tubulin (no mitotic spindle, reduced cytoskeletal dynamics), and fine-granular interphase chromatin (relaxed, non-condensed). The opposition to large heterochromatin-rich nuclei (DAPI_A15) is interesting — quiescent cells here have moderate, fine-granular nuclei rather than large, coarse ones, suggesting these are small/compact quiescent cells rather than senescent cells (which typically have enlarged nuclei). The low ERα (A9) suggests ER-low quiescent cells. The presence of micronuclei (DAPI_A14) may indicate prior mitotic errors before cell cycle exit.

#### P2: Large spread interphase cells with active ERα and translation

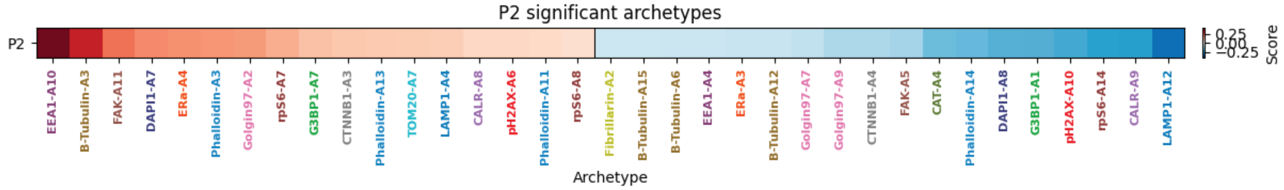


Representative images for archetypes in program:

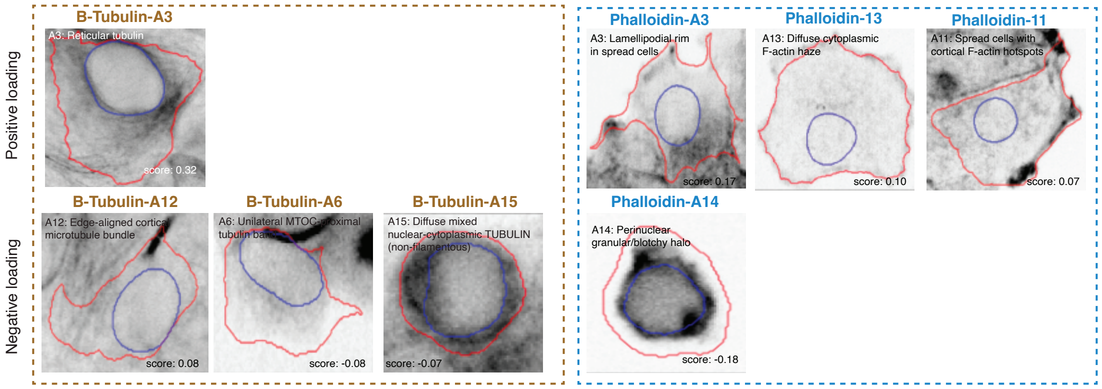


Large, well-spread interphase cells with organized microtubule networks, lamellipodial actin

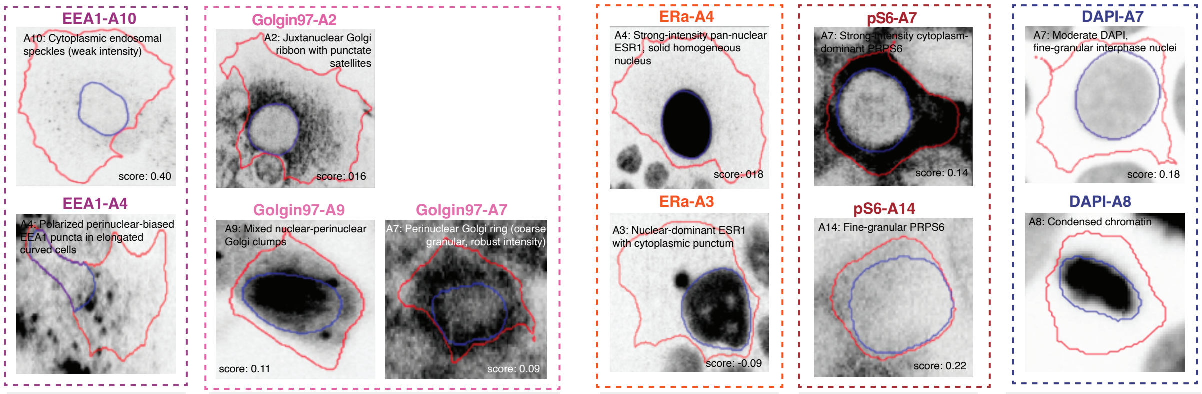


Active endocytic trafficking (EEA1 puncta)

Organized Golgi (juxtanuclear ribbon).

Robust estrogen receptor signaling

High translational activity (cytoplasmic rpS6)

Opposes compact, condensed-chromatin

**Biological interpretation**: P2 captures a fully spread, adherent, interphase cell state with active estrogen signaling, high biosynthetic capacity, and organized cytoskeletal and vesicular trafficking systems. The opposition to condensed chromatin (DAPI_A8) and compact morphology suggests this program distinguishes well-differentiated, actively signaling interphase cells from rounded or mitotic cells. This is consistent with an estrogen-responsive, proliferation-competent but interphase-locked epithelial phenotype.

### Supplementary Figure 19

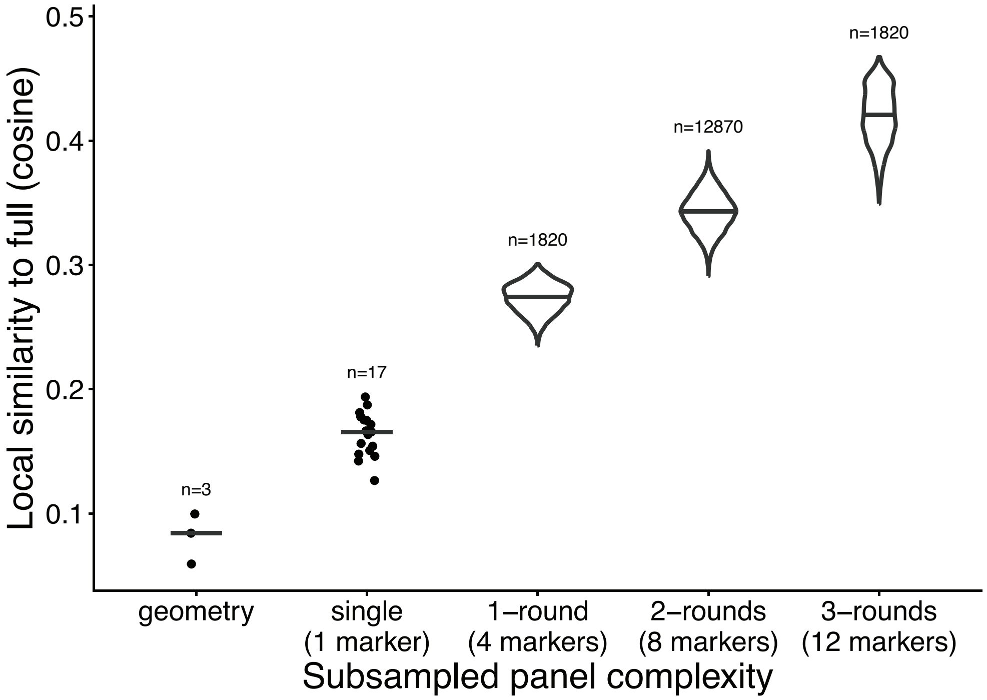

### Supplementary Figure 20

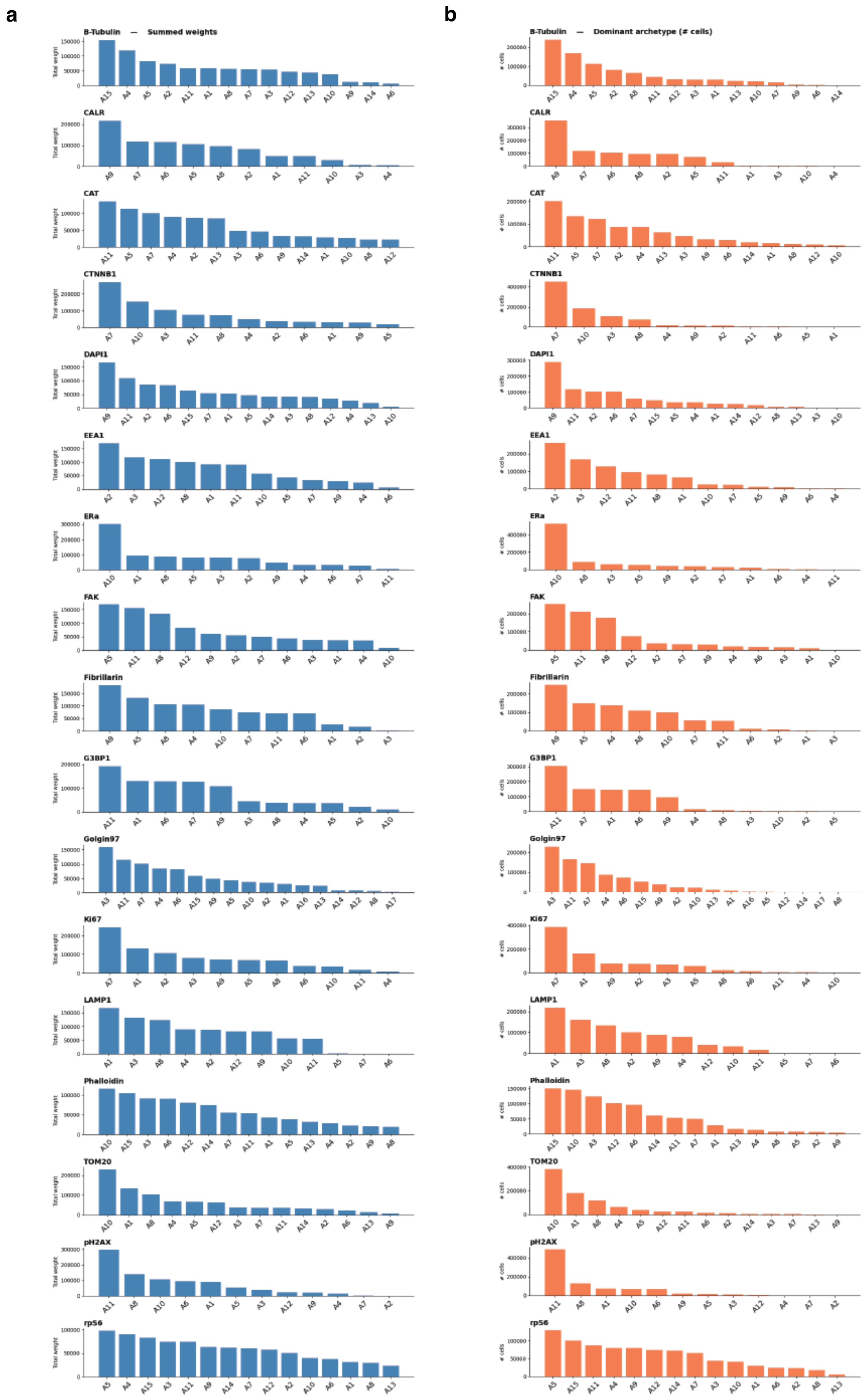

### Supplementary Figure 21

**TOM20-A2**

Representative archetype images:

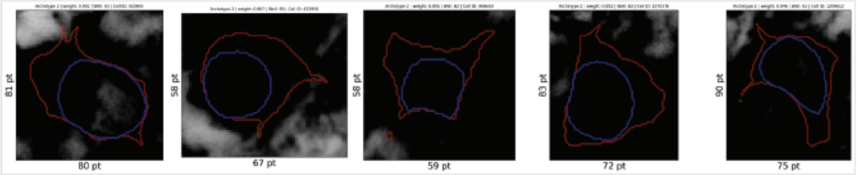

Missannotated MLLM label: Perinuclear-enriched TOM20 ring

Corrected archetype label: Strong pan-cellular TOM20 signal

**Fibrilarin-A2**

Representative archetype images:

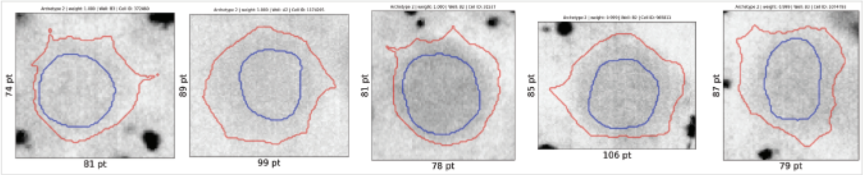

Missannotated MLLM label: Nucleolar puncta on fine nuclear speckle

Corrected archetype label: Completely absent fibrillarin

**Ki67-A6**

Representative archetype images:

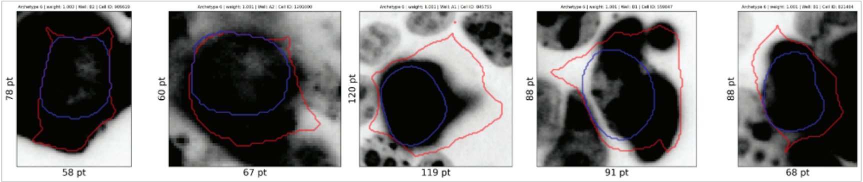

Missannotated MLLM label: Ki-67–high, nuclear-restricted (fine-granular) interphase nuclei

Corrected archetype label: High-intensity Ki67 with some cytoplasmic presence

**pS6-A4**

Representative archetype images:

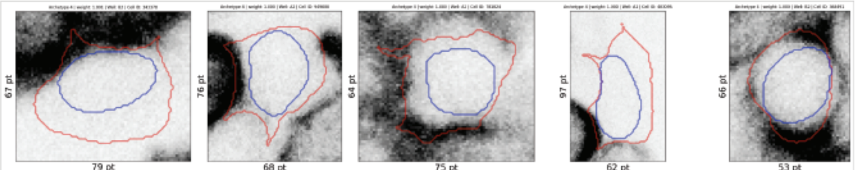

Missannotated MLLM label: Juxtanuclear PRPS6 halo with nuclear clearance

Corrected archetype label: Cortical patches of high-intensity PRPS6
